# Spectral and complexity-based signatures of advanced concentration absorption meditation: A high-density EEG study of jhana

**DOI:** 10.64898/2026.11.15.766880

**Authors:** Peter M. Skov-Andersen, Remko van Lutterveld, Sofie Paludanus, Thomas J. Cahaly, Ruby M. Potash, Andrés Canales-Johnson, Winson F. Z. Yang, Matthew D. Sacchet

## Abstract

Advanced meditation provides structured and reproducible alterations of consciousness that are increasingly used as model states for the neuroscience of consciousness. Jhana-type advanced concentration absorption meditation (ACAM-J) consists of eight progressively deepening absorption states (ACAM-J1–8) in which sensory and self-referential content gradually attenuate. We applied high-density EEG and a comprehensive multidimensional battery of spectral and complexity measures to characterize each ACAM-J state individually in 20 advanced meditators, together with phenomenological ratings. ACAM-J produced widespread posterior-central reductions in oscillatory and aperiodic power, posterior aperiodic flattening across all eight states, and convergent increases in entropy-based complexity over the posterior cortex, with power effects strengthening with phenomenological depth. Crucially, neural complexity remained elevated even in the late, most minimal and unified states, in opposition to the prevailing account that links higher neural entropy to richer phenomenal content. Between practitioners, lower power and higher entropy accompanied more visual and auditory but less bodily content, a redistribution rather than an enrichment of experience; within practitioners, only reduced low-frequency power tracked wider attention and less narrative thought. ACAM-J thus emerges as a non-pharmacological model state at the high-complexity end of the spectrum of altered consciousness, and reveals a previously unquantified dissociation between neural complexity and experiential richness for specific ACAM-J states.

## Introduction

Scientific investigations of meditation have recently begun to expand beyond basic mindfulness meditation practices into what can be referred to as the third wave of meditation research, focusing on advanced meditation ^1–3^. Advanced meditation can be viewed as a set of mental states or stages cultivated with increasing mastery along meditative development, which can give rise to profoundly altered conditions in the form of meditative states and endpoints ^4–6^. In contrast to basic forms of meditation, such as modern mindfulness-based interventions ^7–10^, advanced meditation presents structured and reproducible states that advanced meditators can reliably enter and classify, making it an increasingly important target for studying the neuroscience of consciousness ^2^.

Among the most structured meditative frameworks in Buddhism is jhana, a sequence of reportable absorption states marked by progressive attentional refinement. States based on jhana have been classified and operationalized as jhana-type advanced concentration absorption meditation (ACAM-J) ^11^. ACAM-J comprises eight consecutive states that emerge as absorption and attentional refinement deepen, commonly divided into four early states (J1– J4) and four late states (J5–J8). The early ACAM-J are marked by focused attention, unification of mind, and strong positive affective qualities such as joy, bliss, and equanimity. In contrast, the late ACAM-J involve progressively reduced sensory and self-referential content, accompanied by an experience of boundlessness that culminates in increasingly subtle perception, often described as neither perception nor non-perception. Progression through the ACAM-J is understood to occur through the gradual relinquishing of the dominant qualities of each preceding state, resulting in phenomenologically distinct states that experienced practitioners can reliably recognize and report ^11^.

Consciousness research is increasingly turning to altered states of consciousness to inform hypotheses about brain function ^1,2,12–16^. One framework used in this context is the study of complexity in neural signals. Broadly, entropy-based measures index the predictability of a signal over time, such that more unpredictable dynamics are taken to reflect higher informational complexity. Within this landscape, the entropic brain hypothesis (EBH) ^12,13^ proposes that richer or more unconstrained conscious states are associated with greater neural entropy. Much of this literature has attempted to characterize a spectrum of conscious states using brain complexity measures across conditions such as anesthesia, sleep, and psychedelics.

An initial placement of ACAM-J along this putative spectrum of phenomenological richness, often implicitly framed with psychedelics at one end and anesthesia at the other, is not straightforward. Attention becomes increasingly stable, sensory input fades, subject–object structure weakens, and experiential content radically diminishes. ACAM-J8 is often described in paradoxical terms as a state of neither *perception nor non-perception*, and the late ACAM-J have been discussed in relation to minimal phenomenal experience^17^. ACAM-J, therefore, presents a particularly challenging test case for complexity-based theories of consciousness ^11^. Moreover, increases in neural entropy are not specific to phenomenological richness and may also reflect vigilance, sensory processing, task engagement, or other non-specific factors ^18–21^. Findings of elevated neural complexity during extended cessation, a meditative state described as a complete absence of consciousness ^22^, further underscore the difficulty of assuming a simple one-to-one mapping between neural complexity and conscious content ^23^.

At the same time, “complexity” is not a unitary construct. Entropy-based measures and fractal-scaling measures capture different facets of neural dynamics and may diverge within the same state ^24^. A recent review of meditation research found substantial heterogeneity across metrics, expertise levels, meditation styles, and analytic pipelines, but a general tendency toward increased entropy-related measures in *advanced* meditators ^24–27^, while detrended fluctuation analysis (DFA) seemed to decrease. These patterns highlight the importance of measuring multiple, complementary dimensions of complexity rather than treating complexity as a single quantity.

Prior EEG studies of meditation have often emphasized oscillatory power, yet conventional band-limited estimates may conflate rhythmic oscillations with aperiodic background activity ^28–30^. The aperiodic component is now understood to carry physiologically meaningful information and has been linked to psychiatric and neurological health as well as altered states of consciousness ^31,32^. The aperiodic slope reflects the relative distribution of power across lower and higher frequencies and has been linked to the brain’s excitation–inhibition balance ^30^. Because entropy measures are sensitive to the shape of the power spectrum, separating oscillatory and aperiodic power makes it possible to test whether changes in complexity across ACAM-J reflect changes in oscillatory activity, in aperiodic structure, or in both.

In the present study, we investigate ACAM-J1–8 using a multi-measure EEG framework. We assess oscillatory and aperiodic power, aperiodic slope and offset, and complexity metrics spanning entropy (Lempel–Ziv complexity, permutation entropy, and spectral entropy) and fractal scaling (detrended fluctuation analysis and Higuchi fractal dimension), to determine whether they reveal convergent or dissociable signatures of advanced meditative absorption. Secondarily, we relate these measures to subjective phenomenological ratings obtained after each session, which cover attentional quality, sensory and narrative content, and the defining features of individual ACAM-J. We examine these associations both between practitioners and across states within practitioners. In doing so, we aim to better situate the unique conscious states observed in ACAM-J within contemporary complexity-based accounts of altered consciousness.

## Results

### Study overview

We recorded 128-channel EEG from 20 advanced practitioners while they progressed through ACAM-J1 to ACAM-J8 with eyes closed in a supine position, indicating each transition to a new state with a button press. Before the meditation session, participants completed two non-meditative control tasks under the same conditions: mentally counting down in steps and recalling recent events. For each state, we compared spectral power (total, oscillatory, and aperiodic), the aperiodic exponent and offset, and six complexity measures with the control condition, using linear mixed-effects models of whole-scalp values and cluster-based permutation tests of scalp topography. We then related these measures to state-level phenomenological ratings using partial least squares correlation (see Methods for details).

### Control condition and sample sizes

For the primary analyses, we used the average of the memory and counting conditions as the control condition, consistent with our previous work ^33–35^. This approach was intended to provide a balanced and parsimonious control condition that approximates a more everyday brain state, incorporating both goal-directed cognition and mind-wandering, in advanced meditators. A summary of the contrast between the memory and counting conditions showed differences in frontal theta and alpha power clusters (Supplementary Figure 1), but no differences in average global power. Sample sizes for the entropy and Irregularly Resampled AutoSpectral Analysis (IRASA) analyses are shown in Table 1.

**Table 1.** Sample size for each ACAM-J state for different epoch lengths (column 2-4) and the average time spent in individual ACAM-J states.

| **ACAM-J** | **Entropy measures (2-s epochs), *N*** | **IRASA spectral/aperiodic measures (5-s epochs), *N*** | **DFA α (10-s epochs), *N*** | **Duration, s (M ± SD)** |
|---|---|---|---|---|
| ACAM-J1 | 19 | 18 | 18 | 265 ± 130 |
| ACAM-J2 | 20 | 20 | 20 | 245 ± 131 |
| ACAM-J3 | 20 | 20 | 20 | 280 ± 113 |
| ACAM-J4 | 20 | 20 | 20 | 225 ± 79 |
| ACAM-J5 | 20 | 20 | 20 | 239 ± 99 |
| ACAM-J6 | 19 | 19 | 19 | 220 ± 85 |
| ACAM-J7 | 18 | 17 | 17 | 216 ± 88 |
| ACAM-J8 | 17 | 17 | 15 | 226 ± 135 |

### Power spectral density

We assessed changes in spectral power across ACAM-J1–8 in the delta, theta, alpha, low-beta, and high-beta bands using IRASA applied to 5-s epochs, which reduced residual mixing between oscillatory and aperiodic components compared with 2-s epochs (Supplementary Figure 2; see Methods). For transparency, results from 2-s epochs are shown in Supplementary Figures 3–5. Changes in oscillatory and aperiodic power are shown topographically in Figures 1 and 2 (Total power is shown in Supplementary Figure 6). Across the two power components, the topographies revealed similar broad central clusters of reduced power for most ACAM-J vs. control contrasts and across most frequency bands. High beta effects appeared somewhat more posteriorly distributed than those observed in the delta, theta, and alpha bands. Effects in the delta and theta bands appeared less pronounced for ACAM-J1 and ACAM-J2 than for later states. More generally, cluster size tended to increase from earlier to later ACAM-J states. Global power changes, assessed with linear mixed-effects model (LMM), showed a similar pattern, with progressively larger reductions in total, oscillatory, and aperiodic power across the ACAM-J progression. The same pattern is shown in effect sizes seen across ACAM-J states in Figure 3. Precise estimates and effect sizes for global total, oscillatory, and aperiodic power are reported in Supplementary Data 1.

**Figure 1.**
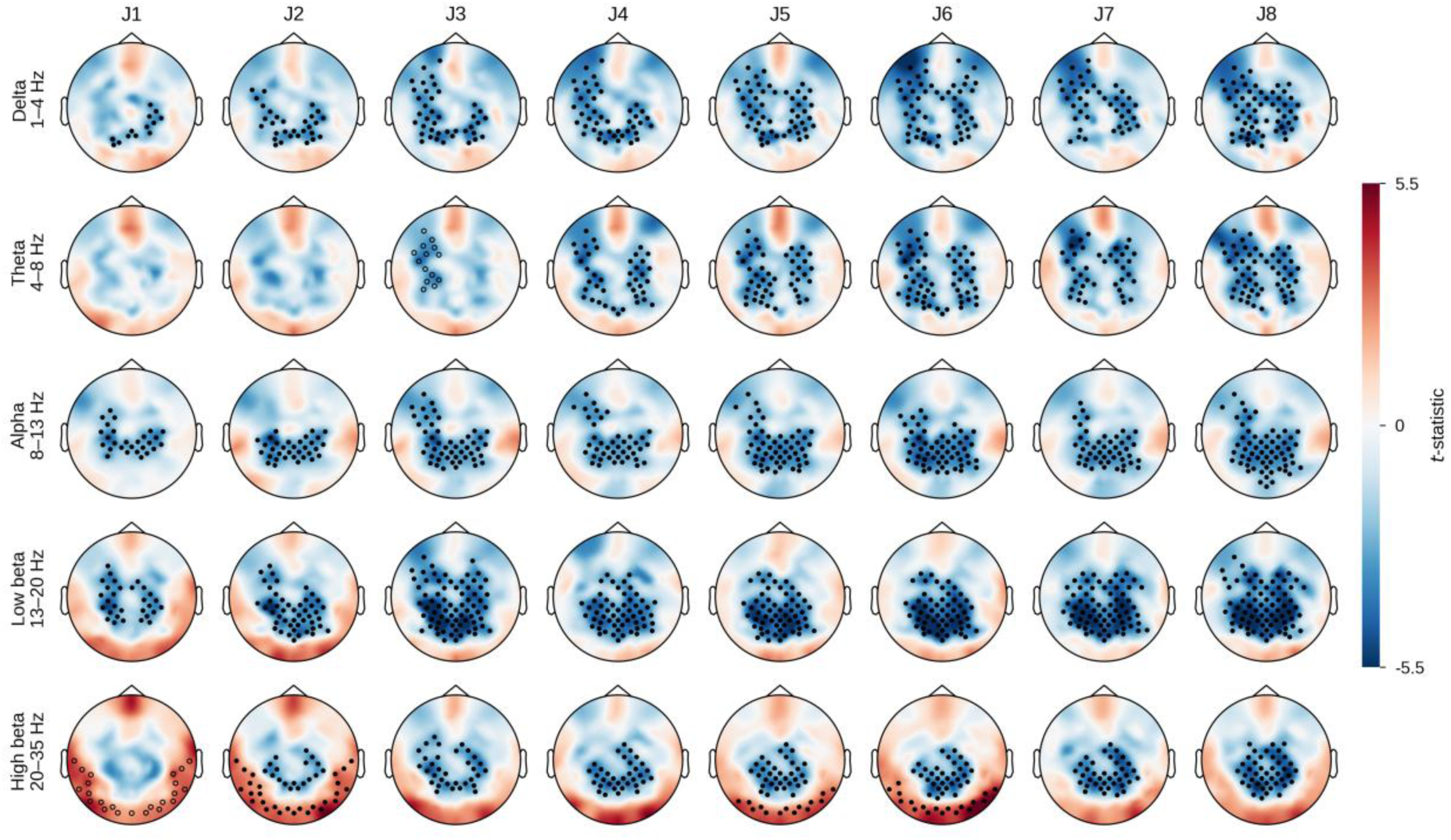
Topography of oscillatory spectral power during ACAM-J1–8. Scalp maps show channel-wise, paired t-statistics for each ACAM-J state minus the combined control condition (mean of memory and counting), for the oscillatory IRASA component estimated from 5-s epochs. Rows show frequency bands and columns show ACAM-J1 to ACAM-J8. Red indicates higher and blue lower power during ACAM-J; the color scale is shared across all maps. Markers show electrodes in significant clusters from a two-sided cluster-based permutation test (cluster-forming threshold P < 0.05; cluster-level P < 0.025; exact null distribution from all sign-flip permutations). Markers are drawn only for contrasts that also survived Benjamini–Hochberg FDR correction across the eight ACAM-J states within each band (q < 0.025). Filled markers show q < 0.01 and open markers show 0.01 ≤ q < 0.025. n per state is given in Table 1. Cluster statistics are given in Supplementary Data 2.

**Figure 2.**
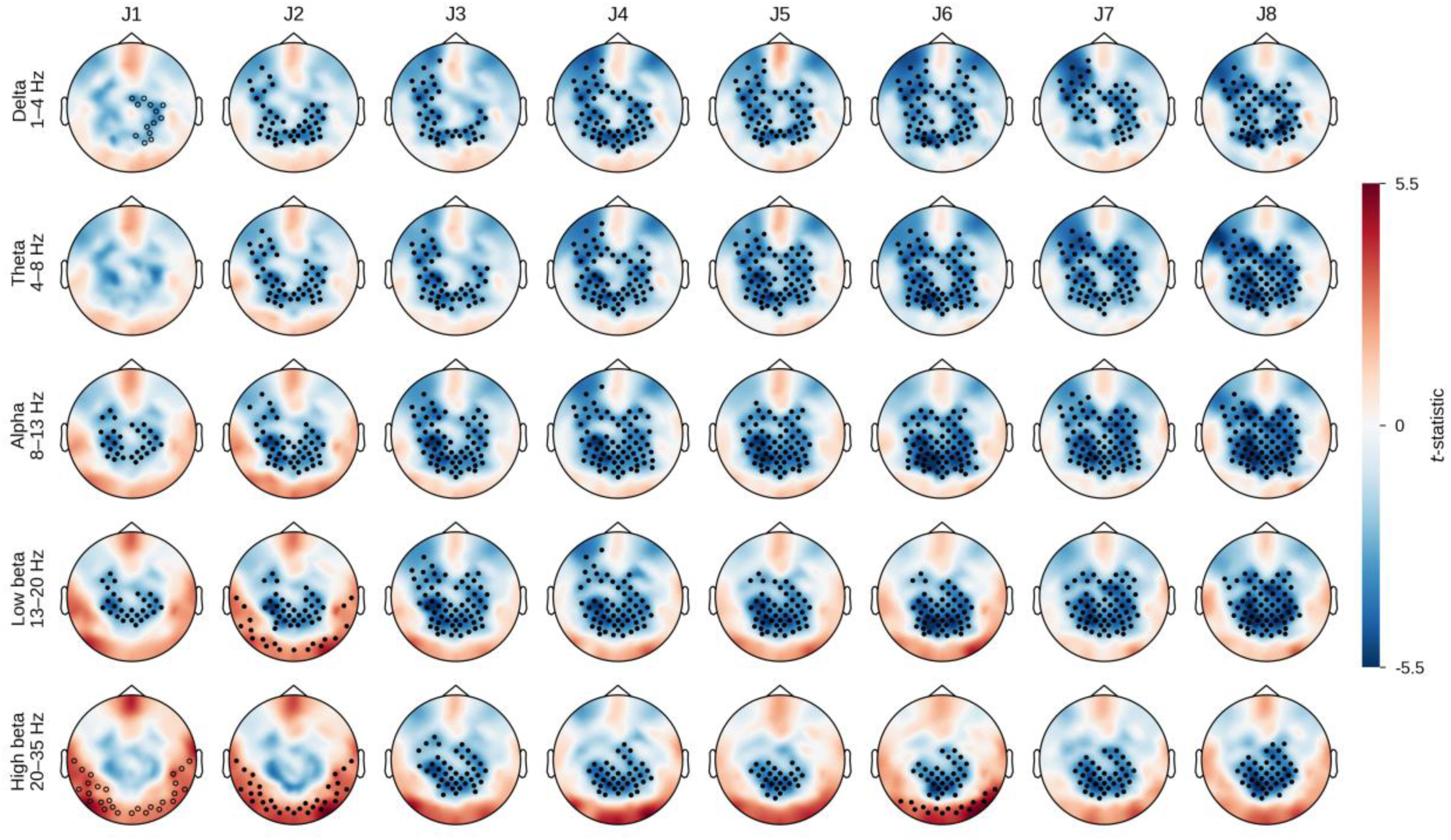
Topography of aperiodic spectral power during ACAM-J1–8. Scalp maps show channel-wise, paired t-statistics for each ACAM-J state minus the combined control condition (mean of memory and counting), for the aperiodic IRASA component estimated from 5-s epochs. Rows show frequency bands and columns show ACAM-J1 to ACAM-J8. Red indicates higher and blue lower power during ACAM-J; the color scale is shared across all maps. Markers show electrodes in significant clusters from a two-sided cluster-based permutation test (cluster-forming threshold P < 0.05; cluster-level P < 0.025; exact null distribution from all sign-flip permutations). Markers are drawn only for contrasts that also survived Benjamini–Hochberg FDR correction across the eight ACAM-J states within each band (q < 0.025). Filled markers show q < 0.01 and open markers show 0.01 ≤ q < 0.025. n per state is given in Table 1. Cluster statistics are given in Supplementary Data 2.

**Figure 3.**
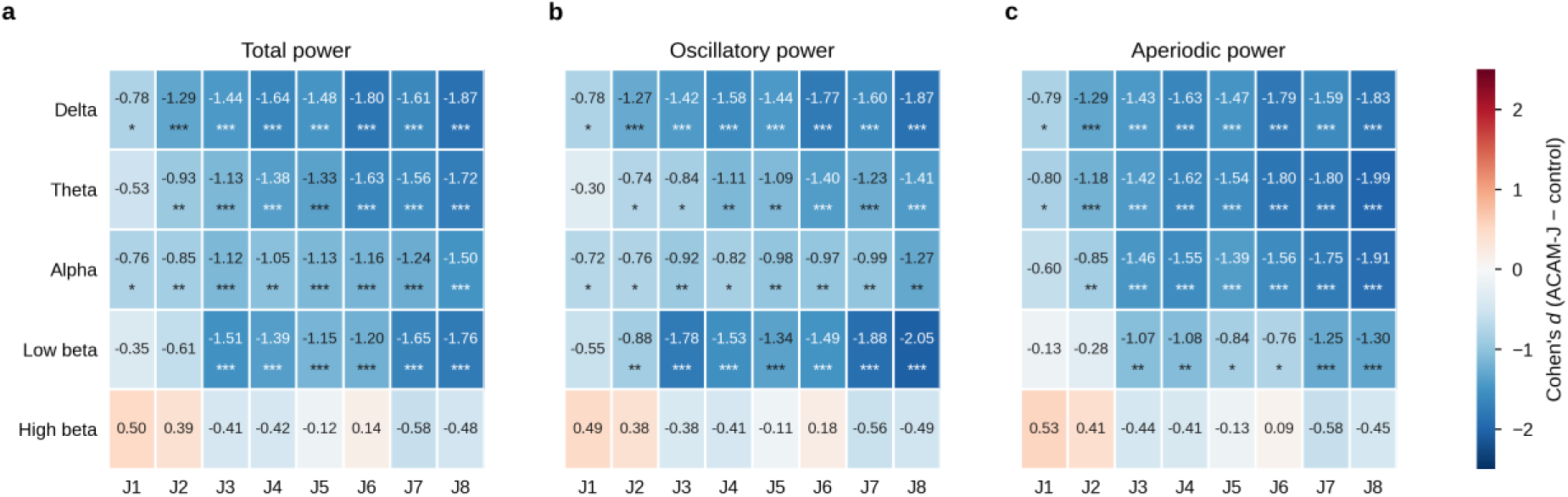
Global spectral power changes across ACAM-J states. Cohen’s d for the difference between each ACAM-J state (J1–J8) and the combined control condition, from linear mixed-effects models of whole-scalp **a**, total, **b**, oscillatory and **c**, aperiodic power (5-s epochs) in five frequency bands. Negative values (blue) indicate lower power during ACAM-J. Asterisks mark contrasts surviving FDR correction across the eight contrasts within each band and component (*P < 0.05, **P < 0.01, ***P < 0.001). Participant-level distributions are shown in Supplementary Figures 9–11. n per state is given in Table 1. Model statistics are given in Supplementary Data 1.

In the high-beta band, posterior clusters of increased power were observed for ACAM-J1, ACAM-J2, ACAM-J5, and ACAM-J6, although no significant posterior positive cluster was identified for the aperiodic component in ACAM-J5. Supplementary Figure 7 shows the full spectrum activity (1–35 Hz), with a pronounced reduction in the alpha peak in Supplementary Figures 7a and 7b and a progressively larger separation between the ACAM-J conditions and the control across all three components. To evaluate whether the posterior high-beta effects could reflect muscle artifact, we correlated the average total-power change in posterior positive clusters, and in centrally located negative clusters for comparison, with self-reported automatic movement during ACAM-J1 and ACAM-J2. No significant correlations were found (Supplementary Figure 8).

Taken together, these findings indicate that ACAM-J is characterized by widespread reductions in total, oscillatory, and aperiodic power, with effects generally becoming more pronounced across successive ACAM-J states.

### Aperiodic exponent and offset

Changes in the aperiodic exponent and offset (3–35 Hz) for ACAM-J1–8 relative to the control condition are shown in Figure 4. Clusters of reduced aperiodic exponent (β) were observed for all ACAM-J states and were located predominantly over posterior scalp regions. Reductions in aperiodic offset were likewise observed across all ACAM-J states, although these effects appeared more broadly distributed across the scalp. Results for low-and high-frequency aperiodic exponents estimated over 3–13 Hz and 13–30 Hz are presented in Supplementary Figure 12. No significant clusters were observed in the 3–13 Hz range, whereas more modest clusters of flattened aperiodic slope were found in the 13–30 Hz range. This suggests that the broadband reduction in aperiodic slope reflects a combination of reduced aperiodic power at lower frequencies below alpha frequencies and increased aperiodic power at higher frequencies.

**Figure 4.**
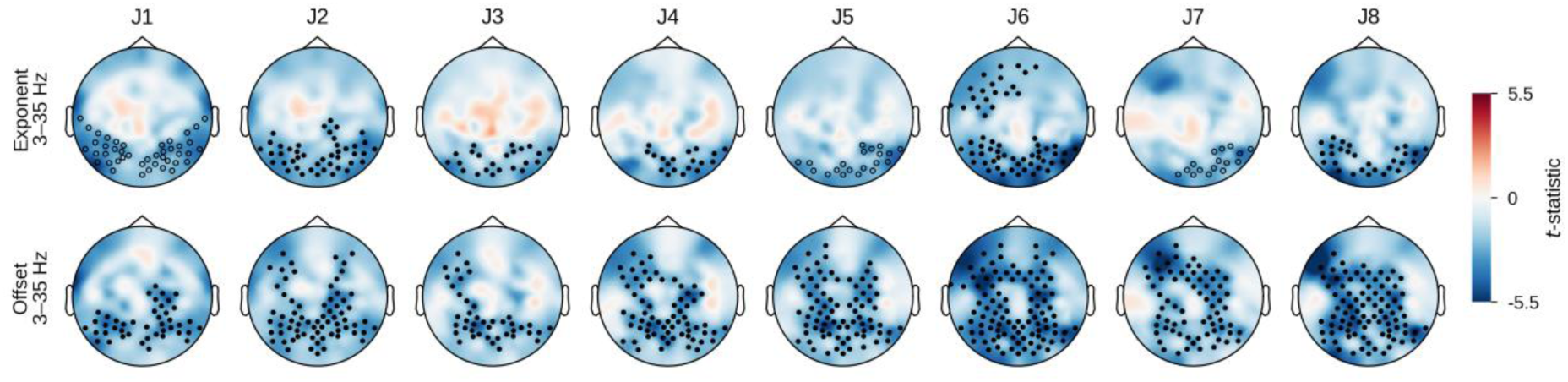
Topography of the aperiodic exponent and offset during ACAM-J1–8. Topographic maps of paired t-statistics for differences in aperiodic features between each ACAM-J state and the combined control condition. Rows show the aperiodic exponent β (3–35 Hz) and offset (3–35 Hz), and columns show ACAM-J1 to ACAM-J8. Positive values indicate higher values during ACAM-J than control. Bloack circles mark electrodes within cluster-significant effects (cluster p < 0.025) in contrasts surviving within-row FDR correction (p < 0.025); filled circles indicate p < 0.01, and hollow circles indicate 0.01 ≤ p < 0.025. n per state is given in Table 1. Cluster statistics are given in Supplementary Data 2.

### Global complexity

Global entropy measures assessed using LMM are shown in Figure 5. LZsum was significantly increased for ACAM-J1, ACAM-J2, ACAM-J5, and ACAM-J6, whereas LZsumN was significantly increased for ACAM-J2, ACAM-J4, ACAM-J5, ACAM-J6, and ACAM-J8. Permutation entropy (PeEn) was increased for all ACAM-J states, with particularly strong effects in ACAM-J1, ACAM-J2, ACAM-J5, ACAM-J6, and ACAM-J8. Overall, ACAM-J was associated with increased entropy as indexed by both compressibility-based measures (LZsum), irrespective of power correction (LZsumN), and Shannon entropy–based measures (PeEn).

**Figure 5.**
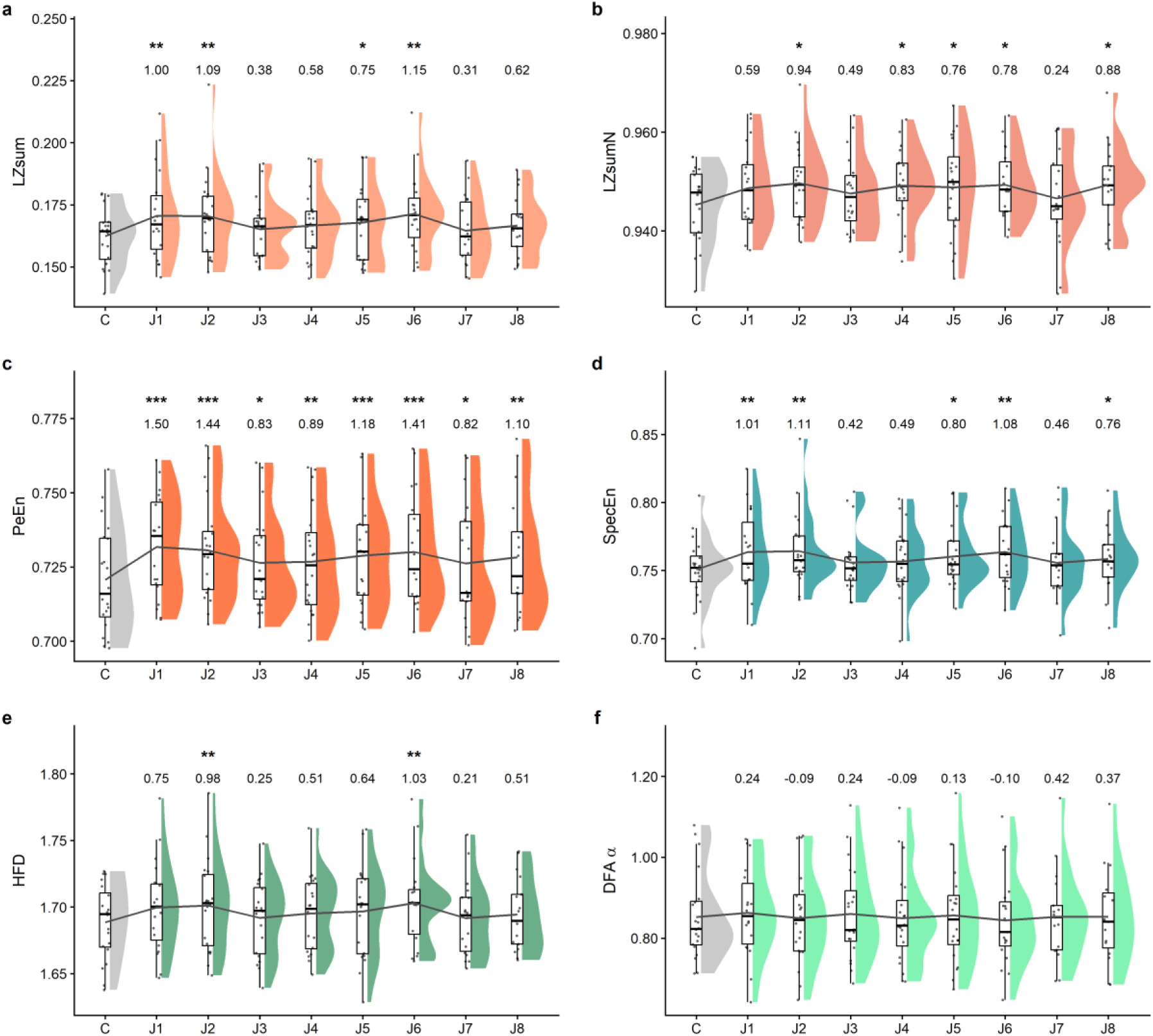
Global signal complexity across ACAM-J states. Raincloud plots of whole-scalp (128-channel average) complexity measures for the combined control condition (C) and ACAM-J1–8 (J1–J8): **a**, Lempel–Ziv complexity (LZsum); **b**, LZsum normalised to phase-randomised surrogates (LZsumN); **c**, permutation entropy (PeEn); **d**, spectral entropy (SpecEn); **e**, Higuchi fractal dimension (HFD); **f**, detrended fluctuation analysis exponent (DFA α). Measures in **a**–**e** were computed from 2-s epochs and DFA α from 10-s epochs. Points show individual participants. Half-violins show the distribution within each state. Boxplots show the median and interquartile range, with whiskers extending to 1.5 × the interquartile range. The grey line connects state means. Numbers above each state give Cohen’s d for the contrast with the combined control condition, estimated from the linear mixed-effects model (estimated mean difference divided by the model residual standard deviation). Asterisks mark contrasts that survive Benjamini–Hochberg FDR correction across the eight contrasts within each measure (*P < 0.05, **P < 0.01, ***P < 0.001). n per state is given in Table 1. Model statistics are given in Supplementary Data 1.

Similarly, global spectral entropy showed significant increases in ACAM-J1, ACAM-J2, ACAM-J5, ACAM-J6, and ACAM-J8. Changes in signal complexity as indexed by fractal dimension were assessed using HFD and DFA, with increased HFD and decreased DFA α taken to indicate increased complexity. HFD increased in ACAM-J2 and ACAM-J6, whereas no significant changes were observed for DFA α.

To investigate the spatial distribution of these complexity changes, we additionally performed cluster-based permutation analyses. These results are shown in Figure 6 and indicate that increases in entropy were primarily localized to posterior scalp regions, whereas central scalp areas were largely unaffected. The posterior distribution of the complexity changes appeared to partially mirror the posterior distribution of the power changes. An exception was observed for PeEn, for which some significant clusters extended into more central regions. Overall, there was good correspondence between the global effects identified with linear mixed models and the spatially resolved effects identified with cluster-based permutation testing, with only minor discrepancies.

**Figure 6.**
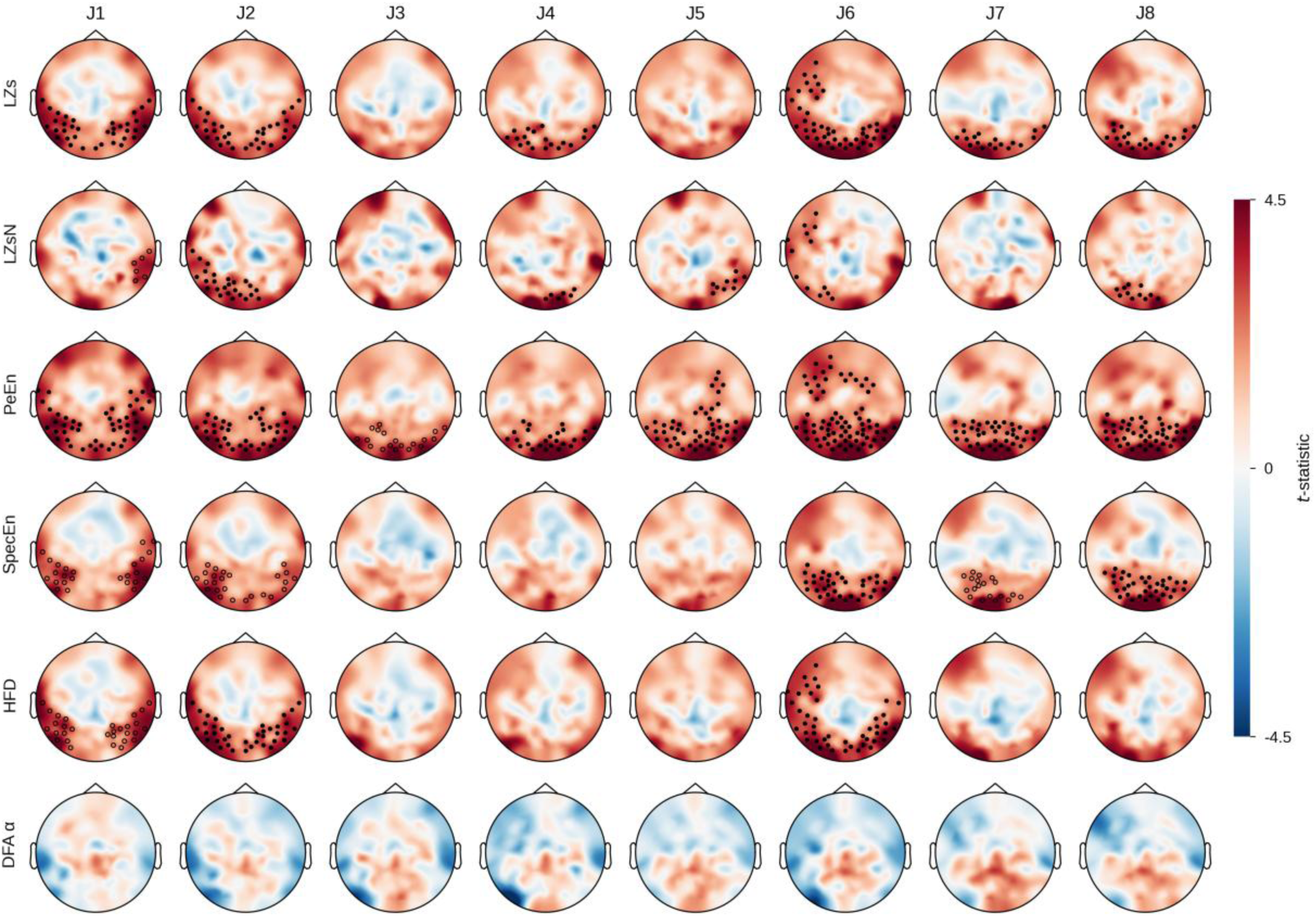
Topography of signal complexity during ACAM-J1–8. Effect of ACAM-J on global complexity. Topographies of paired t-statistics for global complexity measures across ACAM-J–control contrasts. Rows show LZsum, LZsumN, PeEn, SpecEn, HFD, and DFA, and columns show ACAM-J1 to ACAM-J8. A single global symmetric color scale was applied across all panels. Black circles mark electrodes within cluster-significant effects (cluster p < 0.025) in contrasts surviving row-wise FDR correction (p < 0.025), filled circles indicate p < 0.01 and hollow circles indicate 0.01 ≤ p < 0.025. n per state is given in Table 1. Cluster statistics are given in Supplementary Data 2.

Taken together, these findings indicate that ACAM-J states were characterized by a broad increase in neural signal complexity, particularly over posterior scalp regions, alongside reductions in spectral power and aperiodic spectral features. Precise estimates and effect sizes for global complexity measures are reported in Supplementary Data 1.

### Exploratory correlation matrices of global measures

The correlation matrix in Figure 7 is based on ACAM-J–control difference scores that were first computed separately for each ACAM-J state and then averaged across ACAM-J states. For a similar plot for each ACAM-J – control contrast, see Supplementary Figure 13. Three broad patterns emerged. First, changes in total spectral power were strongly and positively intercorrelated across frequency bands, with particularly strong associations among theta, low-beta, and high-beta power. This indicates that participants showing larger ACAM-J changes in one band also tended to show parallel changes in other bands. The strongest associations within this block were observed between theta and low-beta power.

**Figure 7.**
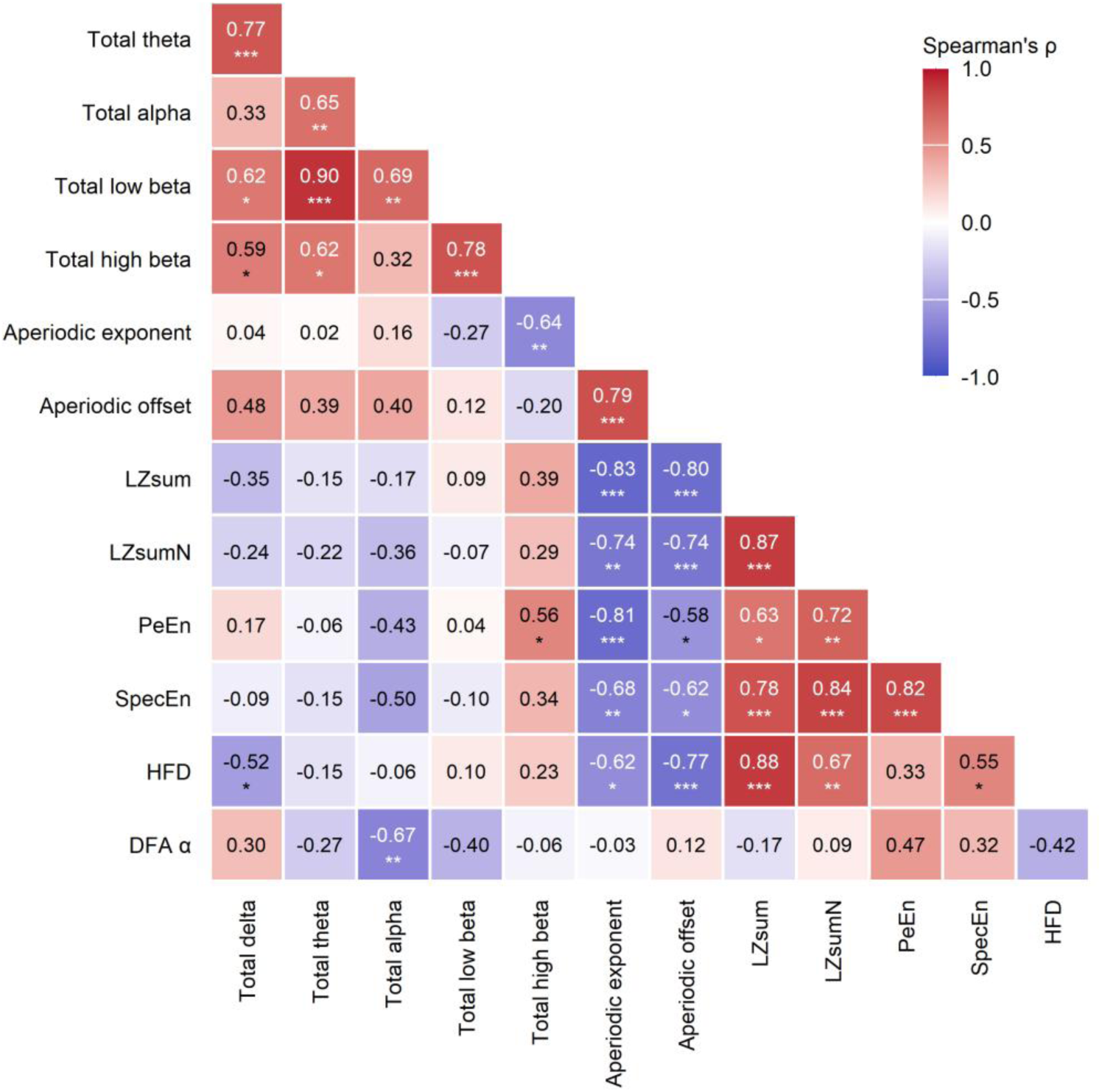
Correlation matrix showing associations among subject-level changes in electrophysiological measures for the mean ACAM-J state relative to the control condition. For each subject, values from ACAM-J1 to ACAM-J8 were averaged within each measure, and the control value was subtracted to yield a single Δ (Mean ACAM-J − Control) score per measure. Tiles represent Spearman correlations across participants between changes in global EEG measures (mean of ACAM-J1–8 minus the combined control; n = 20). Positive correlations indicate that larger increases in one measure were associated with larger increases in another, whereas negative correlations indicate opposing patterns of change. Asterisks mark correlations surviving FDR correction across all 78 pairs (*q < 0.05, **q < 0.01, ***q < 0.001). Per state matrices are shown in Supplementary Figure 13.

Second, the aperiodic measures, β and offset, showed a distinct correlational structure. As expected, aperiodic exponent and offset were strongly positively correlated with one another. Both were also strongly negatively associated with several complexity measures, particularly LZsum, LZsumN, PeEn, SpecEn, and HFD, but not DFA α. Overall, this pattern suggests that greater ACAM-J increases in complexity tended to co-occur with reductions in aperiodic exponent and offset.

Third, the complexity measures themselves formed a correlated cluster. LZsum, LZsumN, PeEn, SpecEn, and HFD were all positively associated with one another, except for the association between HFD and PeEn. This overall pattern indicates substantial convergence across these measures in capturing a shared dimension of ACAM-J–related signal change.

### Brain-phenomenology relationships

Partial Least Squares Correlation (PLSC) was used to relate the sixteen global EEG measures to the seven phenomenological items scored at the state level, across 149 observations from 20 participants. Two complementary analyses were run: a between-participant analysis using the observed values, and a within-participant analysis in which both matrices were centered within participant so that only state-to-state variation remained. Each analysis yielded exactly one significant latent variable, and the two identified different dimensions.

In the between-participant analysis, the first latent variable explained 77.6% of the brain– phenomenology covariance (permutation p = .0001; r = 0.54 between the latent scores; Figure 8a). All ten power measures contributed reliably, and in the same direction, with the largest contributions from the aperiodic theta (-9.9), alpha (-9.8), and low beta (-9.7) components, and three complexity measures contributed reliably in the opposite direction: DFA α (4.4), PeEn (3.1), and SpecEn (2.2) (Figure 8b). Four phenomenological items loaded reliably: sights (0.37), sounds (0.19), and quality of ACAM-J (0.14) at one pole, and physical sensations (−0.26) at the other (Figure 8c). Practitioners with lower spectral power and higher DFA α, permutation entropy, and spectral entropy across their session therefore reported more visual and auditory content, higher quality of ACAM-J, and less physical sensation.

**Figure 8.**
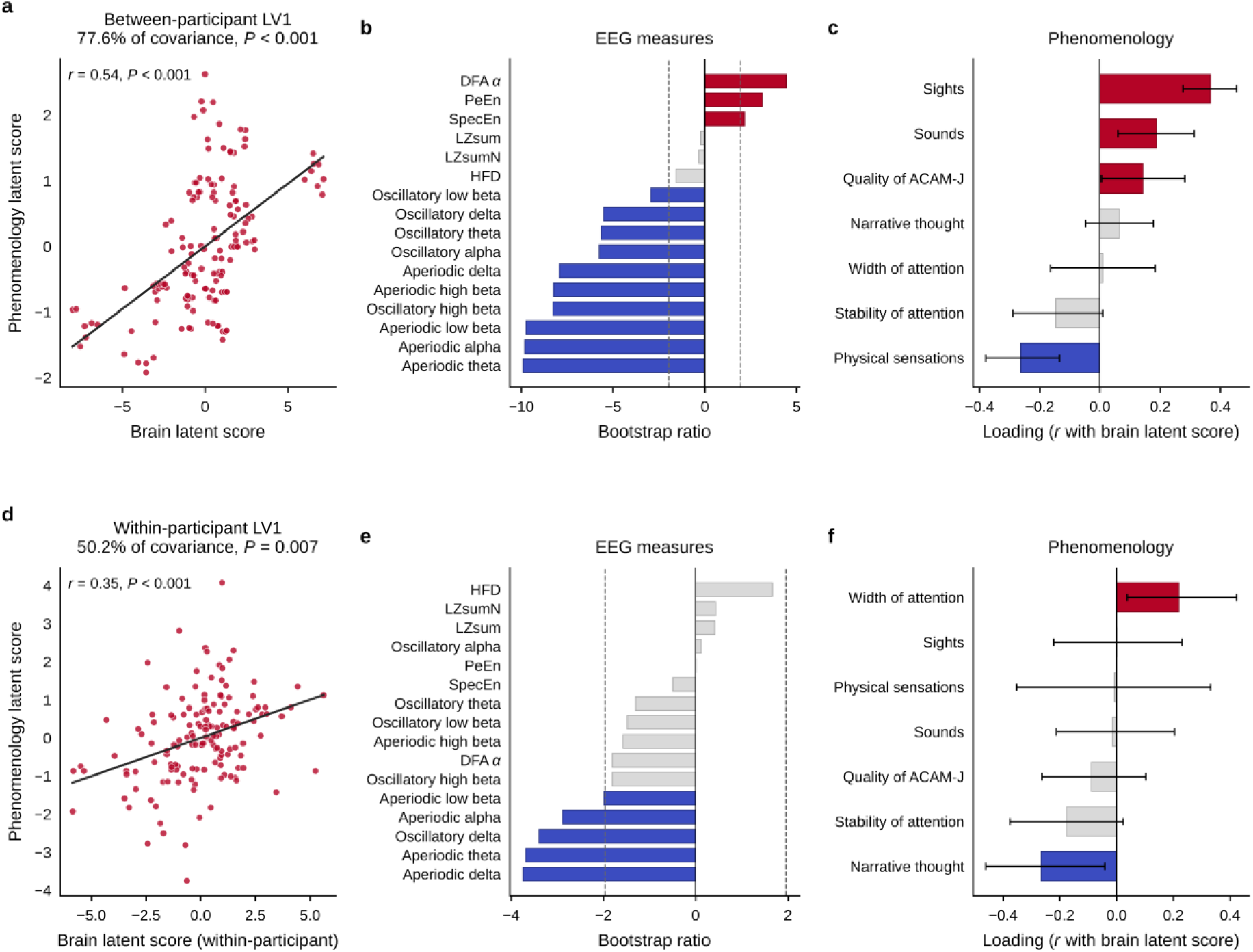
Brain–phenomenology relationships during ACAM-J. Partial least squares correlation linking sixteen global EEG measures to seven phenomenological items across the concatenated ACAM-J states (149 observations, 20 participants). **a–c)** Between-participant analysis. **d–f)** Within-participant analysis, computed after centering both matrices within each participant. Both panels show the first latent variable, the only significant component in each analysis. **a, d)** Latent phenomenology scores against latent brain scores, with the least-squares fit. **b, e)** Contributions of the EEG measures, expressed as bootstrap ratios; dashed lines mark ± 1.96 and colored bars indicate reliable contributions. **c, f)** Phenomenological loadings with 95% bootstrap confidence intervals; colored bars indicate intervals excluding zero. The sign of a latent variable is arbitrary; both are oriented so that the positive pole corresponds to lower spectral power, the direction in which ACAM-J differs from the control condition.

In the within-participant analysis, the first latent variable explained 50.2% of the covariance and was significant (p = .0072; r = 0.35; Figure 8d). Its brain side comprised a range of low-frequency power bands (aperiodic delta (−3.8), aperiodic theta (−3.7), oscillatory delta (−3.4), aperiodic alpha (−2.9), and aperiodic low beta (−2.0)) with no complexity measure contributing reliably (Figure 8e). Width of attention (0.22) and narrative thought (−0.27) loaded reliably, while the four content items and quality of ACAM-J did not (Figure 8f). Within a participant, the ACAM-J states in which low-frequency power fell furthest below that participant’s own average were therefore the states in which attention was reported as wider and narrative thought as less present.

The between-participant dimension relates a combined power and complexity profile to the balance of exteroceptive and bodily content and is expressed as stable differences between practitioners. The within-participant dimension relates low-frequency power alone to attentional scope and discursive thought and is expressed as variation across states within a practitioner.

Two further post hoc analyses were conducted (Supplementary Figure 14). A separate PLSC relating the run-level quality items to EEG measures averaged across the ACAM-J states yielded no significant latent variable (84.2% of covariance, p = .43). Correlations between the EEG measures and the ACAM-J-specific factors yielded no association surviving FDR correction within any factor. The strongest trends were between formlessness and LZsumN (ρ = 0.40), HFD (ρ = 0.36) and DFA α (ρ = −0.37), and between contentment and both permutation entropy (ρ = 0.40) and oscillatory alpha (ρ = −0.40).

## Discussion

To our knowledge, this is the first group study to include the late, formless ACAM-J states and to characterize spectral and complexity-related changes in each of the eight states individually. Three findings emerged. First, ACAM-J was associated with widespread reductions in total, oscillatory, and aperiodic spectral power across the delta, theta, alpha, and low-beta bands, with a predominantly posterior-central scalp distribution and posterior clusters of increased power in the high-beta band for several states. Second, the aperiodic component itself flattened over the posterior cortex during ACAM-J, with reductions in the aperiodic exponent across all eight ACAM-J states and a more broadly distributed reduction in aperiodic offset. Third, ACAM-J produced convergent increases in entropy-based complexity (LZsum, LZsumN, PeEn, SpecEn) over posterior scalp, more modest increases in HFD, and no significant broadband change in DFA. Across spectral measures, effects strengthened across successive ACAM-J states, paralleling the progressive deepening of absorption described phenomenologically, whereas changes in complexity appeared more dispersed across states. Together, these results place ACAM-J within the same neurophysiological neighborhood as classical psychedelics, sub-anesthetic ketamine, breathwork, and other forms of advanced meditation, and contrast with anesthesia, NREM sleep, and disorders of consciousness.

Our spectral-power findings replicate and extend the case-study results of Chowdhury et al. 2025, showing widespread decreases in oscillatory EEG power across multiple frequency bands during ACAM-J^34^. The present cohort reproduces this pattern at the group level, and across total, oscillatory, and aperiodic decompositions of the signal. A group study of ten practitioners reported the same reduction in alpha power, together with an increase in gamma power^37^. That study compared the first four ACAM-J states, pooled, with mindfulness of breathing rather than with a non-meditative control and, like an earlier EEG study of 27 jhana practitioners ^38^, neither resolved individual states nor included the late, formless states. The convergence between previous studies^34,37^ and the present results indicate that posterior alpha attenuation is a reproducible signature of jhana absorption.

The direction of these spectral changes is also consistent with the most reproducible spectral signature of classical 5-HT2A psychedelics. Posterior alpha desynchronisation has been reported across LSD ^31,39^, psilocybin ^40,41^, and DMT ^32,42,43^. Reductions in delta and theta power are also commonly observed under classical psychedelics, although less consistently than the alpha reduction ^39,40^. The posterior topography of our reductions is consistent with the role assigned to occipito-parietal cortex in the psychedelic alpha-desynchronization pattern. The high-beta posterior increases observed in ACAM-J1, ACAM-J2, ACAM-J5 and ACAM-J6 echo the gamma-power increase reported by Mago et al. 2025, although high-frequency increases over posterior scalp warrant cautious interpretation owing to the spectral overlap with electromyographic activity^37^. Several procedures were implemented to minimize EMG contamination and are discussed in the ‘Methodological Considerations’ section in the supplementary material.

The aperiodic findings reinforce this convergence. A flatter 1/f component, indexed by a reduced exponent, has been reported across the psychedelic literature. Muthukumaraswamy and Liley 2018, using IRASA on resting-state MEG, showed that LSD decreased the high-frequency aperiodic exponent across the cortex, and Mago et al. 2025 reported a flatter aperiodic slope during jhana meditation^31,37^. The flatter aperiodic slope is likely a result of decreased lower band and increased high beta band power. This study’s topographical overlap with the posterior high beta band and a flatter slope further suggests this relationship (Figure 1–2 and 4). The same posterior elevation in high-frequency power is also a plausible contributor to the entropy results, as permutation entropy, Lempel-Ziv complexity and spectral entropy are all sensitive to fast, irregular signal fluctuations, and the topography of the entropy increases in this study overlaps with that of the posterior high beta power increase. This raises the possibility that the slope flattening and the rise in complexity share a common signal-level driver in elevated posterior high-frequency activity, rather than being fully independent phenomena. A flatter aperiodic slope has been interpreted as a shift toward an excitation-dominated cortex, on a model in which steeper slopes index greater inhibition and flatter slopes relatively greater excitation ^28,30^. Medel et al. 2023 develop this E/I framework further by proposing that the 1/f slope and signal complexity are not independent properties of a brain state but two windows onto the same underlying quantity, namely the transition entropy of the cortical system. In their account, a flat 1/f spectrum approximates white noise and a high-entropy regime ^44^. Within this framework, the flattening of the aperiodic slope and the increase in entropy observed here can be interpreted as a single shift in cortical regime, with the strong negative correlation between changes in aperiodic slope and entropy across subjects (Figure 7) offering an empirical anchor for this joint interpretation.

Hohn et al. (2024) make the related point that what the slope reflects depends on the frequency range over which it fits. When the fit is taken across a broadband range that includes the oscillatory frequencies, slope and complexity track each other across different sleep stages as well as across rest and active cognitive tasks, with flatter slopes and higher complexity co-occurring in lighter, more wake-like states ^45^. Our 3-35 Hz fit falls in this broadband regime, so the co-modulation of slope, offset, and entropy reported here aligns with this pattern and supports reading ACAM-J as a shift toward greater excitatory drive of posterior cortex rather than a purely oscillatory change. The aperiodic offset is typically interpreted as an index of overall neuronal firing rate or broadband signal level ^28^, and its reduction here is consistent with the more general attenuation of total power. Read together, the simultaneous reduction in low-frequency oscillatory power, in the aperiodic exponent, and in the aperiodic offset describes a posterior cortex that is less synchronized at low frequencies, less amplified overall, and proportionally biased toward higher frequencies.

The increases in EEG signal complexity reported here also replicate and extend the case-study findings of Chowdhury et al. 2025, who reported increases in LZs during ACAM-J^34^. In the present cohort, LZs, LZsN, LZsum, LZsumN, PeEn and SpecEn all increased significantly in several ACAM-J states. The surrogate-normalized LZsumN remained significant for ACAM-J2, ACAM-J4, ACAM-J5, ACAM-J6 and ACAM-J8, indicating that the LZ increase is not solely a consequence of the changes in spectral power reported above ^46^. These increases were primarily localized to posterior scalp regions, mirroring the topography of the aperiodic flattening and consistent with the negative correlation between aperiodic features and complexity reported in our matrix above (Figure 7).

The direction and magnitude of these complexity effects place ACAM-J at the high-complexity end of the spectrum of altered states of consciousness. Anesthesia, NREM sleep and disorders of consciousness occupy the low-complexity pole, with reduced LZ complexity, reduced perturbational complexity, and reduced entropy-based measures more generally ^46–50^. At the high-complexity pole sit classical psychedelics (LSD, psilocybin, ayahuasca, DMT), sub-anesthetic ketamine, and high-ventilation breathwork, all of which increase LZ complexity and related entropy measures ^19,32,41–43,51–56^. Advanced meditation populates this same end of the spectrum ^24,27,37,57^. Our results extend this picture by characterizing the complexity profile of each ACAM-J state individually in a large group study using a wide variety of complexity measures.

The entropic brain hypothesis (EBH) proposes that neural entropy tracks the richness of conscious experience, with classical psychedelics treated as the prototype of the high-entropy regime ^12,13^. Phenomenologically, however, ACAM-J is described as a progressive unification and attenuation of sensory and cognitive content rather than as an enrichment of experience. The deepest ACAM-J states approach what has been described as a minimal phenomenal experience ^17^. If complexity tracked phenomenal richness in the strict sense intended by the EBH, complexity should decrease, not increase, as absorption deepens. Our data show the opposite pattern, and this dissociation does not stand alone. In a study of extended cessations of consciousness, a state in which experience is reported to be entirely absent for minutes at a time, increased complexity was observed in most participants^23^. Three alternative interpretations advanced in that work also apply to ACAM-J. First, the increase in complexity may track sustained, volitionally cultivated attentional engagement rather than phenomenal content. Second, elevated complexity may reflect a latent reorganization of cortical dynamics that supports the post-state clarity and equanimity that participants report. Third, complexity may track deviation from a conventional resting state rather than the richness of experience itself, given that caffeine ingestion ^18^, simple perceptual tasks ^20^, exposure to AI-generated psychedelic-like videos ^58^, and intense external sensory stimulation ^19^ have all been shown to elevate brain entropy without any plausible enrichment of conscious experience.

A recent theoretical reframing of the EBH attempts to resolve precisely this conundrum. The Complex Brain Hypothesis (CBH)^59^ proposes that brain entropy and phenomenal richness need not stand in a one-to-one relation, and that classical psychedelic experiences and minimal phenomenal experience states such as ACAM-J, extended cessation, and 5-MeO-DMT can both show elevated entropy while differing sharply in their phenomenology. The CBH distinguishes the two by the underlying inferential regime in which the brain is operating. High-content psychedelic experiences are modelled as a fine-grained, overfitting regime in which posterior beliefs are pulled far from prior beliefs. Minimal phenomenal experience states such as ACAM-J are modelled as a coarse-grained, underfitting regime in which sensory precision is attenuated by the maintained attentional set, so that posterior beliefs remain close to prior beliefs despite elevated entropy. Phenomenal richness, on this account, is indexed not by signal-level entropy, as captured by the measures used here, but by the divergence between posterior and prior beliefs, which the authors term complexity. The ACAM-J pattern reported here is consistent with the underfitting regime the CBH attributes to minimal phenomenal experience states. Direct empirical tests of the CBH require perturbational measures and computational modelling that the present design cannot support, but the framework offers a principled way to read these results without abandoning the EBH outright.

Taken together, entropy increases appear to be neither specific to psychedelics nor cleanly mapped onto phenomenal richness. The present ACAM-J results suggest a more cautious reading of the EBH framework. High entropy appears to be neither sufficient for, nor consistently aligned with, increased phenomenal richness, and the empirical link between entropy and phenomenology is best treated as one component of a multidimensional account of altered states rather than as a single biomarker.

Although ACAM-J states are thoroughly defined phenomenologically, PLSC analysis makes it possible to directly test the relationship between EEG measures and fine-grained subjective reports. Between practitioners, lower spectral power together with higher DFA α, PeEn, and SpecEn accompanied more visual and auditory content, higher rated quality of ACAM-J, and less physical sensation. Within practitioners, the states in which delta, theta, and alpha power fell furthest below a participant’s own average were those reported with wider attention and less narrative thought.

In a single practitioner, Chowdhury et al. 2025 found width of attention to be positively correlated with widespread increases in Lempel-Ziv complexity and stability of attention with increases in alpha and theta power. Our within-participant dimension also links attentional width to the EEG, although here it is carried by reduced low-frequency power rather than by complexity, and stability of attention did not load reliably in either analysis. Using ReHo, Yang et al. (2025) reported a first latent variable that separated sensory and narrative items from attentional qualities, with quality of ACAM-J grouping with width and stability of attention. The present between-participant dimension also separates sensory from bodily experience, but places quality of ACAM-J with sights and sounds rather than with the attentional items.

For the EBH, the direction of these effects matters. If signal complexity indexed phenomenal richness, it should covary positively with reported content and should do so both across individuals and across states. Between practitioners, the low-power, higher-entropy profile accompanied more exteroceptive but less bodily content, indicating a redistribution of experiential content rather than an overall enrichment. Within practitioners, complexity did not predict experience at all, and the width of attention increase was carried by low-frequency power alone. This is consistent with the argument above, in which signal-level entropy and the content of experience are only loosely coupled, and it argues against treating any single complexity measure as a proxy for phenomenal richness. It also suggests that the more tractable within-participant signal is elevated slow-wave power accompanying mind-wandering and altered attentional scope, which matches other findings in the literature ^60,61^.

Despite the convergent picture across entropy-and aperiodic-based measures, broadband DFA α showed no significant change between ACAM-J and the control condition. This null finding parallels Mago et al. 2025, who likewise reported no broadband DFA effect during jhana, but did report changes in specific bands (reduced in theta, alpha, and beta and increased in gamma). DFA requires longer continuous epochs of clean data, and the smaller DFA sub-sample may have reduced statistical sensitivity. DFA computed over shorter epochs has been shown to be sensitive to brief, relatively fast modulations of long-range scaling associated with advanced meditation events ^62^. The broadband null should therefore be interpreted with caution rather than as evidence against criticality-related changes during ACAM-J.

Several limitations should be acknowledged. The sample size (N=20 for most states), although the largest to date for the full ACAM-J sequence, restricts statistical power for low-effect-size measures, particularly for HFD and DFA. The within-subject design speaks to state effects only and cannot speak to long-term trait differences in baseline complexity following sustained practice ^24^. The cohort also has a considerable sex imbalance (19 male, 1 female).

Several directions could build on this work. Source-level MEG could localize the posterior effects to specific generators (posterior cingulate, occipital, or parietal cortex). fMRI measures of neural complexity could test whether ACAM-J modulates large-scale brain dynamics, adding cross-modal convergence.

Finally, we assessed a large battery of outcome measures. Within each measure, the eight ACAM-J versus control contrasts were FDR-corrected, but we did not impose an additional correction across the full set of measures. This reflects the design of the study, which was intended as a comprehensive characterization of ACAM-J across complementary facets of neural dynamics rather than a single confirmatory test of one hypothesis. Thus, imposing a family-wise correction across all measures would be overly conservative given that the measures are neither independent nor of equal *a priori* interest. Notably, the between-measure correlation matrix (Figure 7) shows that the effects do not behave as scattered, independent findings: they cluster into theoretically expected groupings, with the entropy-based complexity measures co-varying with one another, the aperiodic exponent and offset forming a distinct block, and the spectral bands intercorrelating tightly. This coherent structure is difficult to reconcile with chance-level findings, which would be expected to distribute randomly across measures, and instead points to a small number of underlying, convergent dimensions of ACAM-J–related signal change.

In conclusion, ACAM-J is associated with a coordinated reorganization of the EEG signal that places it at the high-complexity end of the spectrum of altered states of consciousness. The spectral fingerprint, namely posterior reductions in oscillatory and aperiodic power and a flatter 1/f, mirrors the fingerprint reported under classical psychedelics, sub-anesthetic ketamine, breathwork, and other forms of advanced meditation. The complexity fingerprint, namely convergent increases in entropy-based measures alongside more modest increases in HFD, places ACAM-J in the same neurophysiological neighborhood as the state induced by psychedelics. Our findings replicate and extend the case-study results of Chowdhury et al. 2025 at the group level and across each individual ACAM-J state, and they converge with the recent independent preprinted jhana dataset of Mago et al. 2025. Together, these results establish ACAM-J as a non-pharmacological model state with a reproducible electrophysiological signature, and provide a quantitative basis for further investigation of the neural dynamics of ACAM-J.

## Methods

### Participants

Twenty-one advanced meditators were recruited as part of a larger project to investigate the neural correlates and phenomenology of different advanced meditative states. All participants were assessed as having a robust and reliable capacity to enter ACAM-J. Most participants were also affiliated with a well-established meditation community recognized for mastery of ACAM-J. One participant was excluded (see Preprocessing), leaving 20 participants (19 males, 1 female; age 26–67 years, mean 46.0 ± 14.4) with a median of 10,450 lifetime meditation hours (IQR 4,925–23,083; range 1,120–118,225). The subjects provided informed consent, and the study received approval from the Mass General Brigham Institutional Review Board (IRB) Protocol 2019P003902 and the ethics boards at Witten/Herdecke University.

### Advanced concentration absorption meditation

The participants were instructed to follow their normal ACAM-J routine (from ACAM-J1 to ACAM-J8) with their eyes closed, in supine position, and indicate the transition to a new ACAM-J state by pressing a button.

### Control tasks

We designed two non-meditative control tasks to engage participants in routine, everyday thought processes without eliciting meditation. A resting-state condition was avoided because experienced meditators might unintentionally enter meditative states, which could complicate the interpretation of the results. This is also consistent with recent work pointing to the limitations of the classical resting-state framework^63^. Instead, participants performed a counting task and a memory task, both with closed eyes, identical to those in our previous investigations^33,34,64–68^. During the counting task, participants were asked to mentally count down from 10,000 in steps of five for 8 min without speaking out loud or moving their lips. In the memory task, they were asked to mentally revisit and narrate the events that occurred in the past 2 weeks for 8 min without speaking out loud or moving their lips. Each task was done 1-2 times for each participant depending on which meditation tasks they were completing for the broader study. The control tasks were performed under the same conditions (eyes closed, supine) and recorded before the ACAM-J session to avoid carry-over effects from ACAM-J.

### Data acquisition

EEG data were recorded as continuous MEG-EEG activity on a TRIUX system at Massachusetts General Hospital, featuring 4 EOG channels and 128 EEG channels from a customized 128-channel MEG BrainCap system. The MEG-EEG system was located in an Imedco magnetically shielded room, with a shielding factor of approximately 250,000 at 1 Hz. The sampling frequency was 1000.0 Hz, with a high-pass filter of 0.1 Hz and a low-pass filter of 330.0 Hz. Task instructions were provided verbally and displayed on a screen inside the booth. The participants had access to a button box with buttons 1-4 to navigate through instruction slides and indicate their progression through the states of ACAM-J.

### Preprocessing

EEG data was pre-processed offline in the MATLAB environment (MATLAB Version: 23.2.0.2599560 (R2023b) Update 8, The Mathworks, Massachusetts, USA) using FieldTrip ^69^ and the EEGLAB toolbox ^70^. Data was imported into EEGLAB as raw .fif files. MEG and EOG channels were removed, and transition events, indicated by button presses, were added from a stimulus channel.

Preprocessing was primarily carried out using automated and well-validated procedures to minimize experimenter bias and inconsistencies across individuals, and increase reproducibility and standardization ^71,72^. Data cleaning was performed using the RELAX_wICA_ICLabel approach using the EEGLAB toolbox RELAX ^73,74^ (RELAX 2.0). All electrode positions were standardized to a standard realistic head model template with the 10-5 system ^75^. The sampling rate was kept at 1000 Hz. Line noise was removed with an acausal 4th-order Butterworth filter (57 and 63 Hz). The data was then bandpass filtered using a linear (zero-phase) non-causal finite impulse response filter (pop_eegfiltnew) with high-pass filtering at 1 Hz for optimal ICA performance ^76,77^ and a low-pass filter at 80 Hz. Bad electrodes were first removed using PREP’s ‘findNoisyChannels’ ^78^. RELAX’s default rejection thresholds were further applied if the total channel electrode was under 10%. Then, the extreme outlier periods were marked and removed for further processing using RELAX’s default extreme outlier settings (improbable voltage value distributions (SD): 10, single channel kurtosis: 10, all channel kurtosis: 10, absolute voltage shift: 1000, MAD from median voltage shift: 25, MAD from median voltage shift in blink affected epochs: 10, log-frequency, log-power slope threshold for drift: -4). The data were re-referenced using PREP’s robust average reference ^78^. ICLabel ^79^ then identified artifactual muscle, eye, heart, line noise, and channel noise components with the thresholds 0.8, 0.8, 0.9, 0.9, 0.8, respectively. Artifactual components were reduced using wavelet-enhanced independent component analysis (wICA) ^80^, using the PICARD algorithm ^81,82^. wICA uses wavelet thresholding to identify and remove high-amplitude oscillations in ICA components, presumed to be artifacts, while preserving lower-amplitude neural activity. Finally, all cleaned components are back-projected to the scalp space, and the rejected electrodes were interpolated. For a more detailed explanation of RELAX_wICA_ICLabel, see Bailey, Biabani, et al., 2023.

All runs were checked for clusters of rejected electrodes where reasonable interpolation was not possible, and subjects/runs were excluded if deemed necessary. Runs with more than 10% of electrodes rejected underwent visual inspection of their data quality. The data was segmented into non-overlapping epochs. Noisy epoch removal was carried out with single-channel joint probability/kurtosis 5 SD, all-channels 3 SD, absolute amplitude ± 120 µV, similar to other publications using RELAX for epoching ^83,84^. Epoch length was 2-s for LZ complexity measures, permutation entropy, spectral entropy, Higuchi’s fractal dimension, 5-s for spectral measures, and 10-s for DFA. 2-s epochs have been used in previous advanced meditation studies ^34,85^ and for various studies of neural entropy in altered states of consciousness ^16,42,54^. The data was then further low-pass filtered at 35 Hz using EEGLAB’s ‘pop_eegfiltnew’ for all measures except power, where the data was kept at 1-80 Hz. Lastly, as in other studies of entropy and complexity ^46,86^, we applied a surface Laplacian montage using the spherical spline method, scalp current density (SCD) EEGLAB plugin ^87^, with a smoothing constant of λ = 1×10⁻⁵. The SCD estimates the second spatial derivative of the scalp potential from a spherical-spline interpolation across all electrodes. This attenuates spatially broad signals, including volume-conducted activity and the reference, so that each channel more closely reflects local cortical sources beneath the electrode^88^. In addition, SCD in combination with ICA, as used here, has been shown to minimize any EMG contamination in frequencies above 25 Hz ^89^.

One subject was excluded due to clusters of rejected electrodes. One ACAM-J run from a participant with two ACAM-J runs was excluded for the same reason. Individual ACAM-J states were excluded from analyses if the total number of epochs summed to < 30 seconds of clean data.

### Comparisons

The primary analysis of this investigation compares each ACAM-J with a combined control condition. The combined control condition was calculated as the unweighted average of the memory and counting conditions, assigning equal weight to each, regardless of the epoch counts of the two conditions. See Supplementary Figure 1 and Supplementary Data for comparisons between the two control conditions.

### Total, oscillatory, and aperiodic power

Total, oscillatory, and aperiodic power were estimated within the following frequency bands: delta (1-4 Hz), theta (4-8 Hz), alpha (8-13 Hz), low-beta (13-20 Hz), and high-beta (20-35 Hz). The beta range was split into low-and high-beta sub-bands to obtain a beta-range measure below 20 Hz (13-20 Hz) that is less contaminated by muscle activity, which predominantly affects frequencies above ∼20 Hz ^89,90^. The spectrum was divided into oscillatory and aperiodic components using Irregularly Resampled AutoSpectral Analysis (IRASA) ^91^.

The oscillatory spectrum was obtained by subtracting the aperiodic spectrum from the total spectrum. Small negative values can arise in some cases when the aperiodic estimate slightly exceeds the total spectrum. Therefore, any negative values were set to zero before averaging.

IRASA was implemented using FieldTrip on clean 2- and 5-s epochs, similar to Timmermann et al., 2023, to output the total spectrum and the aperiodic spectrum for each epoch and electrode with h ranging from 1.1 to 1.9 in increments of 0.05. Analyses were restricted to 1– 70 Hz, which safely covers our highest frequency of interest (35 Hz), given the maximum resampling factor of 1.9. IRASA was initially applied to 2-s epochs but was repeated on 5-s epochs to reduce residual mixing between the oscillatory and aperiodic components (Supplementary Figure 2). All spectral and aperiodic results reported in the main text are based on 5-s epochs.

### Aperiodic slope estimation

Following IRASA-based separation of aperiodic and oscillatory components, the aperiodic power spectrum was averaged across epochs in linear power for each subject, state, and electrode, and the aperiodic exponent (β) was then estimated by linear regression in log-log space (log10 power as a function of log10 frequency) ^31^. The intercept of this same regression was retained as the aperiodic offset, reflecting the overall vertical position of the fitted aperiodic spectrum in log-power space. The primary exponent was estimated over 3–35 Hz. This range was chosen to avoid fitting directly at the 1-Hz high-pass edge and because, in IRASA, the evaluated frequency range extends beyond the nominal fitting range, making low-frequency edge effects problematic ^29^. In addition, supplementary exponents were estimated over 3–13 Hz (*β* _lo_) and 13–30 Hz (*β* _hi_) to assess whether the aperiodic profile differed between the lower- and higher-frequency portions of the spectrum ^92^ (Supplementary Figure 12). For interpretability, results were reported as the positive spectral exponent (β = −slope), with larger values indicating a steeper decay of aperiodic power with frequency.

### Choice of complexity measures

Consistent with Atad et al., 2025, the present study treats neural complexity as a multidimensional construct rather than a unitary property and therefore includes measures that capture partly distinct aspects of brain dynamics.

### Lempel-Ziv Complexity

Single-channel Lempel-Ziv complexity (LZs) was computed using the LZ76 algorithm ^93^ from the EntRate toolbox ^94^. For each 2-s epoch of length *T* samples and for each channel, the time series was binarized around the median, turning points above the median into 1’s and points below into 0’s. The total number of unique patterns, the LZ76 count (C), is the output, which grows with increasing irregularity or information richness in the sequence. The LZ76 count can be normalized to the length of T and provides an entropy rate ^19,94,95^.

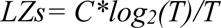

To control for changes in the spectral power profile, we additionally computed a phase-shuffled surrogate normalization of Lempel-Ziv complexity (LZsN), which is C relative to the mean of 100 phase-randomized Fourier transform surrogates of the continuous signal Surrogates were then binarized around the median, and LZ76 complexity was computed.

However, Mediano and colleagues have argued that surrogate-based normalization can be problematic in between-condition comparisons, because differences in the spectra also change the surrogate baselines. Consequently, the difference between normalized values may reflect differences between the null models themselves, potentially producing false positives rather than isolating a purely non-spectral effect ^94–96^. We therefore report both measures, to facilitate comparison with prior LZ complexity work, while basing our main inferential claim on the entropy-rate. For this study, we also report average LZs (LZsum) and LZsN (LZsumN) across all channels, similar to Lewis-Healey et al., 2024 and Timmermann et al., 2023. Note that we refrained from using a Hilbert-amplitude binarization because the instantaneous phase of broadband data is not well defined or interpretable ^19^.

### Permutation entropy

Permutation entropy (PeEn) ^97^ is a measure of signal irregularity that quantifies the unpredictability of the ordering of the signal pattern. PeEn was computed using EntropyHub ^98^ on the broadband signal (1-35 Hz) using m = 3 and τ =4 samples (4 ms at 1000 Hz), equivalent to *τ* = 1 at 250 Hz, implemented with a stride of 2 by retaining every other sample.

### Spectral entropy

Spectral entropy (SpecEn) is a measure of the dispersion of spectral power across frequencies. SpecEn was estimated using MATLAB’s spectralEntropy function ^99^ from the power spectrum in the 1–35 Hz range. Each epoch was analyzed as a single Hamming-tapered segment, and spectral entropy was computed from the normalized distribution of spectral power across frequency bins within this range.

### Higuchi fractal dimension

Higuchi fractal dimension (HFD) was used to quantify the scale-invariant temporal complexity of the EEG signal. Conceptually, HFD estimates how irregular or self-similar a time series remains when examined across progressively larger temporal intervals. The highest *k*-value, *k*_max_, was selected based on saturation analysis. For a subset of runs, we computed the mean HFD over a sweep of *k*_max_ values and chose the smallest value in the flattening region of the curve (see Supplementary Figure 15), balancing stability with computational cost. Based on this procedure, *k*_max_ = 200 was used for all analyses. After selecting K_max_=200, HFD was computed using the Higuchi_FD function ^100^.

### Broadband detrended fluctuation analysis

Brain criticality was assessed using broadband detrended fluctuation analysis on epoched data, similar to other studies ^101,102^. We quantified scale-free long-range temporal correlations (LRTC), i.e., temporal memory, in the EEG time series by estimating the DFA exponent, α ^103^. The DFA computation was performed using a modified version of DFluC (Colombo et al., 2016) that supported custom window sizes. In short, broadband DFA (1-35 Hz) was estimated by splitting the epochs into non-overlapping windows with log-spaced lengths from 100 samples to 1000 samples. Within each window, the second-order polynomial trend was removed, and the fluctuation quantified as the root mean square of the residuals pooled across all windows. This was repeated for all 50 window scales. The power-law relationship between fluctuations and window sizes is then estimated via linear regression on a log-log plot, with the slope being equal to the DFA exponent α. Window sizes for broadband DFA were chosen based on guidance from previous research ^104^, setting the minimum window sizes as n_min_=f_s_/f_max_ and maximum window sizes as n_max_=f_s_/f_min_. With f_s_=1000 Hz and a 1-35 Hz band yielding n_min_=28.6 samples (29 ms) and n_max_=1000 samples (1.0 s). Previously, small-scale curvature from possible filter artifacts has been observed ^104,105^. We investigated our choice of window sizes by visual inspection of log-log plots, assessing linearity in the data for the given box sizes using the first epoch for all ACAM-J1-8 and control runs across various electrodes, finding 100 samples as the minimum most suitable (see Supplementary Figure 16). We chose to analyze 10-second epochs, as this provides 10 boxes at the largest scale and therefore a reasonable number of windows available for averaging, as recommended^106^.

### Statistics

Cluster-based permutation tests ^107^ were carried out with FieldTrip by computing a two-sided dependent-samples t-statistic at every channel (depsamplesT) with a cluster-forming threshold of α=0.05 and with cluster-level α = 0.025 (two-sided, corresponding to an overall FWER of 0.05). Cluster significance was assessed with a max-sum statistic using all possible sign-flip permutations of the subject-wise condition labels (i.e., all possible re-assignments of ACAM-J vs control per subject). Electrode neighbors were determined using FieldTrip’s triangulation method, and clusters were required to contain at least two neighboring electrodes. Multiple contrasts (8 ACAM-J conditions) were further controlled using Benjamini–Hochberg false discovery rate (FDR) applied separately within each frequency band (delta, theta, alpha, low beta, high beta) and IRASA spectral component (total, oscillatory, aperiodic), within each complexity measure. Cluster statistics for all contrasts (cluster P, FDR q, cluster mass, size, peak t and electrode lists) are provided in Supplementary Data 2.

Differences in global complexity and power were analyzed using a linear mixed-effects model (LMM) with state as a fixed effect and a subject-specific random intercept to account for repeated measures. Combined control was computed for each participant first, and the model had nine levels (combined control plus ACAM-J1–8). Models were fit using restricted maximum likelihood in lme4^108^ with degrees of freedom and p-values obtained via Satterthwaite’s approximation (lmerTest). Planned contrasts compared each meditation state (ACAM-J1–8) to the combined control condition using estimated marginal means (emmeans). The statistical significance of the eight planned contrasts was controlled using FDR. Memory vs counting was tested in a separate model. We further computed standardized effect sizes for each planned contrast as Cohen’s *d* (emmeans::eff_size), defined as the estimated mean difference from the LMM divided by the model residual standard deviation (σ).

For spectral and entropy measures, runs belonging to the same subject and condition were pooled across epochs, and channel-wise values were calculated for each subject and state for cluster-based measures and averaged across all channels for global measures analyzed with LMMs. Full model estimates, 95% CIs, Satterthwaite df, uncorrected and FDR-corrected P values, Cohen’s d and omnibus F tests are provided in Supplementary Data 1.

### Between-metric correlation analysis

For each ACAM-J state, subject-level delta scores were computed for all global metrics as the difference between the ACAM-J and control conditions. Pairwise Spearman rank-order correlations were then calculated across subjects between all delta measures, producing one correlation matrix per ACAM-J state. To account for multiple testing, p-values for all unique pairwise correlations within each matrix were adjusted using FDR.

### Phenomenology

Immediately after each recording session, participants completed a structured retrospective questionnaire, developed in collaboration with advanced meditators and meditation instructors, on which they scored, from 0 to 10, how strongly a range of experiential features had been present. The items differ in the temporal resolution at which they are reported, which determines how each enters the analyses below. Three items were scored for every ACAM-J separately: *stability of attention*, ranging from poor (0) to excellent (10) stability; *width of attention*, from narrow and point-like (0) to wide and panoramic (10); and *quality of ACAM-J*, how closely the experience corresponded to the canonical description of that particular absorption. Four content items: *sights*, *sounds*, *physical sensations,* and *narrative thought*, were scored twice per session rather than per ACAM-J. Once covering the early ACAM-J states (ACAM-J1-4) and once covering the later states (ACAM-J5-8), yielding an early and a late value for each. Five items are tied to particular ACAM-J states and were scored only there: *bliss/joy* and *automatic movements* (ACAM-J1-2), *contentment* (ACAM-J3), *equanimity* (ACAM-J4), and *formlessness* (ACAM-J5-8). Four remaining items refer to the session as a whole: *overall quality*, *sense of unity*, *sense of self,* and *luminosity*. Two participants completed two ACAM-J sessions. Where both sessions survived EEG quality control, scores were averaged across them. Where only one survived, the scores from that session alone were used.

### Brain-phenomenology analyses

Relationships between the global EEG measures and phenomenology were examined with partial least squares correlation (PLSC), a multivariate method that extracts latent variables (LVs) capturing patterns of covariance between two sets of variables ^109–111^.

Observations were structured at the state level, one row per participant per ACAM-J state. The brain matrix comprised sixteen global measures: LZsum, LZsumN, HFD, DFA, SpecEn, and PeEn, together with aperiodic and oscillatory power in the delta, theta, alpha, low beta, and high beta bands. Total power was not included alongside its own two components, since the three parameterizations are not independent. The phenomenology matrix comprised the seven items scored at the level of individual or aggregate states: quality of ACAM-J, stability of attention, width of attention, and sights, sounds, physical sensations, and narrative thought. For the four content items, the early score was assigned to ACAM-J1-4 and the late score to ACAM-J5-8. Observations with missing EEG values were removed, leaving 149 observations from 20 participants.

Two analyses were run on these data. In the between-participant analysis, the matrices were used as observed. Because a participant’s global EEG values vary far less across the ACAM-J sequence than they do between participants, this analysis is dominated by stable differences between practitioners. In the within-participant analysis, both matrices were first demeaned within participants, removing each participant’s own mean and leaving their deviations across states, so that any recovered covariance must arise from state-to-state variation inside individuals.

Both analyses were carried out with the pyls package (v0.0.2; ^112^). Statistical significance of each latent variable was assessed with 10,000 permutations, with permuted solutions Procrustes-rotated onto the original decomposition before their singular values were evaluated ^111^. Permutations were unrestricted in the between-participant analysis and constrained to occur within participant in the within-participant analysis, so that in each case the null preserves the structure the design is not testing. The Kaiser criterion was additionally applied, retaining only components explaining more variance than the average across components. Latent variables were interpreted only when both criteria were met. Reliability of the weights was estimated with 10,000 bootstrap iterations, resampling observations in the between-participant analysis and participants in the within-participant analysis. Contributions of individual EEG measures are reported as bootstrap ratios, with bootstrap ratio > 1.96 considered reliable, and phenomenological loadings with 95% bootstrap confidence intervals. Pearson correlations were computed between the latent brain and phenomenology scores.

The four run-level items (overall quality, sense of unity, sense of self, luminosity) could not enter these analyses, since a single score repeated across eight states carries no within-participant variance and would violate the independence assumed by the resampling procedures. They were instead examined in a separate PLSC in which the EEG measures were averaged across the eight ACAM-J states, so that each participant contributed one observation (n = 20).

Because the ACAM-J specific factors are scored only for the states in which they are defining features, they were analysed separately, each as its own family of tests. For each factor, both the score and the EEG measures were averaged across the relevant states (bliss/joy across ACAM-J1-2, contentment at ACAM-J3, equanimity at ACAM-J4 and formlessness across ACAM-J5-8) so that each participant contributed one observation. Spearman rank-order correlations were computed between each EEG measure and each factor, consistent with the between-metric correlation analysis reported above and appropriate to the ordinal scores. P-values were adjusted using FDR within each factor.

### Stability analysis

As a supplementary check of the chosen minimum epoch thresholds, we examined the stability of global total, oscillatory, and aperiodic power, LZsum, LZsumN, PeEn, SpecEn, HFD, and DFA as a function of the number of clean epochs included. Quantitative analyses using intraclass correlation and a non-overlapping split-half procedure indicated that 15 2-s epochs for complexity measures, six 5-s epochs for IRASA, and three 10-s epochs for DFA provided sufficiently stable estimates, except for DFA, LZsumN, and the theta-band contrasts (Supplementary Figures 17–21). A qualitative inspection, similar to van Lutterveld ^36^, likewise showed averages plateauing at these epoch counts (Supplementary Figure 22). These counts were therefore used as the minimum for a run to be included in the DFA (3 epochs), power (6 epochs), and complexity (15 epochs) analyses, and most participants far exceeded them (DFA: 19.5 ± 8.8; power: 39.0 ± 17.7; complexity: 99.8 ± 44.7 epochs). See ‘Extended methods’ in the supplementary material for a detailed description of the procedures.

## Supporting information

Supplementary Data 1 - Linear mixed-effects model statistics

Supplementary Data 2 - Cluster-based permutation test statistics

## Supplementary Data

**Supplementary Data 1. Linear mixed-effects model statistics for global EEG measures.** For each global power and complexity measure: estimates, standard errors, 95% confidence intervals, Satterthwaite degrees of freedom, *t* values, uncorrected and Benjamini–Hochberg FDR-corrected *P* values, and Cohen’s *d* with 95% confidence intervals. These cover each ACAM-J state versus the combined control and counting versus memory, together with omnibus *F* tests of state.

**Supplementary Data 2. Cluster-based permutation test statistics for topographic analyses.** For each contrast (ACAM-J1–8 versus the combined control, and memory versus counting) across power, aperiodic and complexity measures: *n*, number of permutations, minimum cluster *P*, FDR *q* across the eight ACAM-J states, and for each cluster its direction, size, cluster statistic, peak *t*, peak electrode, approximate location and member electrodes. Analysis parameters are also included.

## Data Availability

Data may be requested and is subject to the Massachusetts General Hospital Institutional Review Board’s guidelines and approval.

## Code Availability

All custom code used for preprocessing, spectral and complexity analyses, statistics, PLSC and more is available at: https://github.com/peterskov-andersen/acam-j_power_complexity

## Author Contributions

Conceptualization: MDS, RvL

Methodology: PMS-A, RvL, MDS, SP, TC, AC-J

Investigation: RP, WFZY, PMS-A

Formal analysis: PMS-A

Visualization: PMS-A

Supervision: RvL, MDS

Writing – original draft: PMS-A

Writing – review & editing: PMS-A, RvL, MDS, WFZY, TC, SP, RP, AC-J

All authors reviewed and approved the final manuscript.

## Acknowledgement and funding

Dr. Sacchet and the Meditation Research Program are supported by the Dimension Giving Fund, Tan Teo Charitable Foundation, and additional individuals.

