## Supplementary Information for "Spectral and complexity-based signatures of advanced concentration absorption meditation: A high-density EEG study of jhana"

### Supplementary material

### Extended methods

#### Participants

Twenty-one advanced meditators were recruited as part of a larger project to investigate the neural correlates and phenomenology of different advanced meditative states. All participants were assessed as having a robust and reliable capacity to enter ACAM-J. Most participants were also affiliated with a well-established meditation community recognized for mastery of ACAM-J. The participant group consisted of 20 males and 1 female, with an age range spanning 28-67 (mean = 45.95 ± 14.00). Participants had a median of 10,450 lifetime meditation hours (IQR 4,600–23,170). The subjects provided informed consent, and the study received approval from the Mass General Brigham Institutional Review Board (IRB) Protocol 2019P3902 and the ethics boards at Witten/Herdecke University.

#### Preprocessing

EEG data was pre-processed offline in the MATLAB environment (MATLAB Version: 23.2.0.2599560 (R2023b) Update 8, The Mathworks, Massachusetts, USA) using the fieldtrip [8] and the EEGLAB toolbox [9]. Data was imported into EEGLAB as raw .fif files. MEG and EOG channels were removed, and transition events, indicated by button presses, were added from a stimulus channel.

Preprocessing was primarily carried out using automated and well-validated procedures to minimize experimenter bias and inconsistencies across individuals, and increase reproducibility and standardisation [10,11]. Data cleaning was performed using the RELAX_wICA_ICLabel approach using the EEGLAB toolbox RELAX [12,13] (RELAX 2.0). All electrode positions were standardized to a standard realistic head model template with the 10-5 system [14]. The sampling rate was kept at 1000 Hz. Line noise was removed with an acausal 4th-order Butterworth filter (57 and 63 Hz). The data was then bandpass filtered using a linear (zero-phase) non-causal finite impulse response filter (pop_eegfiltnew) with high-pass filtering at 1 Hz for optimal ICA performance [15,16] and a low-pass filter at 80 Hz. Bad electrodes were first removed using PREP's 'findNoisyChannels' [17]. RELAX's default rejection thresholds were further applied if the total channel electrode was under 10%. Then, the extreme outlier periods were marked and removed for further processing using RELAX's default extreme outlier settings (improbable voltage value distributions (SD): 10, single channel kurtosis: 10, all channel kurtosis: 10, absolute voltage shift: 1000, MAD from median voltage shift: 25, MAD from median voltage shift in blink affected epochs: 10, log-frequency, log-power slope threshold for drift: -4). The data were re-referenced using PREP's robust average reference [17]. ICLabel [18] then identified artifactual muscle, eye, heart, line noise, and channel noise components with the thresholds 0.8, 0.8, 0.9, 0.9, 0.8, respectively. Artifactual components were reduced using wavelet-enhanced independent component analysis (wICA) [19], using the PICARD algorithm [20,21]. wICA uses wavelet thresholding to identify and remove high-amplitude oscillations in ICA components, presumed to be artifacts, while preserving lower-amplitude neural activity. Finally, all cleaned components are back-projected to the scalp space, and the rejected electrodes were interpolated. For a more detailed explanation of RELAX_wICA_ICLabel, see Bailey, Biabani, et al., 2023.

All runs were checked for clusters of rejected electrodes where reasonable interpolation was not possible, and subjects/runs were excluded if deemed necessary. Runs with more than 10% of electrodes rejected underwent visual inspection of their data quality. The data was segmented into non-overlapping epochs. Noisy epoch removal was carried out with single-channel joint probability/kurtosis 5 SD, all-channels 3 SD, absolute amplitude ± 120 µV, similar to other publications using RELAX for epoching [22,23]. Epoch length was 2-s for LZ complexity measures, permutation entropy, spectral entropy, Higuchi's fractal dimension, 5-s for spectral measures, and 10-s for DFA. 2-s epochs have been used in previous advanced meditation studies [1,24] and for various studies of neural entropy in altered states of consciousness [25-27]. The data was then further low-pass filtered at 35 Hz using EEGLAB's 'pop_eegfiltnew' for all measures except power, where the data was kept at 1-80 Hz. Lastly, as in other studies of entropy and complexity [28,29], we applied a surface Laplacian montage using the spherical spline method, scalp current density (SCD) EEGLAB plugin [30], with a smoothing constant of $\lambda = 1\times10^{-5}$. The SCD estimates the second spatial derivative of the scalp potential from a spherical-spline interpolation across all electrodes. This attenuates spatially broad signals, including volume-conducted activity and the reference, so that each channel more closely reflects local cortical sources beneath the electrode[31]. In addition, SCD in combination with ICA, as used here, has been shown to minimize any EMG contamination in frequencies above 25 Hz [32].

One subject was excluded due to clusters of rejected electrodes. One ACAM-J run from a participant with two ACAM-J runs was excluded for the same reason. Individual ACAM-J states were excluded from analyses if the total number of epochs summed to < 30 seconds of clean data.

### Comparisons

The primary analysis of this investigation compares each ACAM-J with a combined control condition. The combined control condition was calculated as the unweighted average of the memory and counting conditions, assigning equal weight to each, regardless of the epoch counts of the two conditions. See Supplementary Figure 1 and Supplementary Data 1 for comparisons between the two control conditions

#### Total, oscillatory, and aperiodic power

The EEG power spectrum can be described as the sum of narrowband oscillatory components and a broadband aperiodic component. In the power spectral density (PSD) frequency domain, the oscillatory component appears as distinct peaks, such as the alpha peak. On the other hand, the PSD of the aperiodic component follows a continuous 1/f power-law distribution, $P(f) = 1/f^{\beta}$, where $\beta$ denotes the spectral slope in log–log space [33]. As the aperiodic component exhibits self-similarity across temporal scales, it is sometimes also referred to as *fractal* or *scale-free* activity [34]. The aperiodic component was for a long time referred to as “1/f noise” but it has recently gained more traction as it is being related to neurological and psychiatric disorders [35] and shows sensitivity to pharmacological interventions [36-38], suggesting that it reflects broader aspects of brain function. While oscillatory peaks are typically interpreted as reflecting transient synchrony among neural populations, the generative mechanism of aperiodic activity remains incompletely understood. The most common interpretation is as a marker of the balance between synaptic excitation and inhibition (Donoghue, 2025; Gao et al., 2017), but other mechanisms, such as dendritic filtering [39], have also been proposed. Given the potential role of 1/f-like activity in cognition and the fact that oscillatory peaks ride on top of this broadband background, we sought to separate oscillatory and aperiodic activity when quantifying ACAM-J effects on the PSD.

Total, oscillatory, and aperiodic power were estimated within the following frequency bands: delta (1-4 Hz), theta (4-8 Hz), alpha (8-13 Hz), low-beta (13-20 Hz), and high-beta (20-35 Hz). The beta range was split into low- and high-beta sub-bands to obtain a beta-range measure below 20 Hz (13-20 Hz) that is less contaminated by muscle activity, which predominantly affects frequencies above ~20 Hz [32,40]. The spectrum was divided into oscillatory and aperiodic components using Irregularly Resampled AutoSpectral Analysis (IRASA) [41].

IRASA separates oscillatory from aperiodic activity by exploiting the fact that narrowband rhythms are tied to specific frequencies, whereas the aperiodic 1/f component is approximately scale-invariant. For each epoch, the time series was irregularly resampled using a set of non-integer factors *h* and their reciprocals *1/h*. Resampling by *h* and *1/h* shifts the frequency of any oscillatory peaks while leaving the aperiodic component approximately invariant in the log-log power-frequency space. For each resampling factor, IRASA computes a power spectrum, and the geometric mean of the h and *1/h* spectra is taken. Because the oscillatory peaks move in frequency across resampling factors while the aperiodic background does not, this geometric mean cancels out oscillatory contributions and yields an estimate of the aperiodic spectrum. The oscillatory spectrum was obtained by subtracting the aperiodic spectrum from the total spectrum. Small negative values can arise in some cases when the aperiodic estimate slightly exceeds the total spectrum. Therefore, any negative values were set to zero before averaging, since negative residuals do not have a straightforward physical interpretation as power and are typically treated as artifacts of imperfect separation, similar to previous work [42,43].

IRASA was implemented using FieldTrip on clean 2- and 5-s epochs**,** similar to Timmermann et al., 2023, to output the total spectrum and the aperiodic spectrum for each epoch and electrode with h ranging from 1.1 to 1.9 in increments of 0.05. Analyses were restricted to 1–70 Hz, which safely covers our highest frequency of interest (35 Hz), given the maximum resampling factor of 1.9.

### Aperiodic slope estimation

The slope of aperiodic activity has been studied across various altered states [36,37,44,45]. The aperiodic slope was estimated to summarize the relative distribution of aperiodic power across lower and higher frequencies, that is, whether the spectrum became steeper or flatter across a given range. Following IRASA-based separation of aperiodic and oscillatory components, the aperiodic power spectrum was averaged across epochs in linear power for each subject, state, and electrode, and the aperiodic exponent (β) was then estimated by linear regression in log-log space (log10 power as a function of log10 frequency) [36]. The intercept of this same regression was retained as the aperiodic offset, reflecting the overall vertical position of the fitted aperiodic spectrum in log-power space. The primary exponent was estimated over 3–35 Hz. This range was chosen to avoid fitting directly at the 1-Hz high-pass edge and because, in IRASA, the evaluated frequency range extends beyond the nominal fitting range, making low-frequency edge effects problematic [46]. In addition, supplementary exponents were estimated over 3–13 Hz ($\beta_{lo}$) and 13–30 Hz ($\beta_{hi}$) to assess whether the aperiodic profile differed between the lower- and higher-frequency portions of the spectrum [47]. For interpretability, results were reported as the positive spectral exponent (β = −slope), with larger values indicating a steeper decay of aperiodic power with frequency.

#### Choice of complexity measures

Consistent with Atad et al., 2025, the present study treats neural complexity as a multidimensional construct rather than a unitary property and therefore includes measures that capture partly distinct aspects of brain dynamics. PeEn, SpecEn, and LZs all quantify entropy-related features of the signal but emphasize different properties: PeEn is sensitive to the unpredictability of ordinal patterns, SpecEn reflects the distribution of power across the frequency spectrum, and LZs index the diversity or compressibility of temporal patterns. In contrast, HFD and DFA assess a different facet of complexity by characterizing scale-free temporal structure in terms of self-similarity, and long-range temporal correlations. Considered together, these measures provide a more comprehensive characterization of meditation-related changes in neural dynamics, as they allow differentiation between changes in signal irregularity and fractal temporal structure.

#### Lempel-Ziv Complexity

Lempel-Ziv complexity was computed to compare brain activity during ACAM-J with other altered states of consciousness, including anaesthesia [29,48-50], psychedelics [26-28,38,49,51-58], breathwork [59], sleep [44,50,60], meditation [55,61] and our case study of ACAM-J [1]. Single-channel Lempel-Ziv complexity (LZs) was computed using the LZ76 algorithm [62] from the EntRate toolbox [63]. For each 2-s epoch of length $T$ samples and for each channel, the time series was binarized around the median, turning points above the median into 1's and points below into 0's. The binary string is fed to the incremental parsing LZ76 algorithm that quantifies how often the sequence produces a new pattern. The total number of unique patterns, the LZ76 count (C), is the output, which grows with increasing irregularity or information richness in the sequence. The LZ76 count can be normalized to the length of T and provides an entropy rate [52,63,64].

$$LZs = C * log_2(T)/T$$

Schartner et al. 2015 introduced phase-shuffled surrogate normalisation (LZsN) to control for changes in the power spectra to be the driver of complexity change and have since been reported in various studies [29,38,49,54,57]. To control for changes in the spectral power profile, we additionally computed a phase-shuffled surrogate normalisation of Lempel-Ziv complexity (LZsN), which is C relative to the mean of 100 phase-randomized Fourier transform surrogates of the continuous signal [29]. For each epoch, we took the FFT of the signal, preserved the power spectrum but shuffled the phases of the Fourier coefficients (uniformly distributed in [0,2π] with appropriate conjugate symmetry), followed by an inverse FFT back to the time domain, yielding surrogates that share the original power spectrum but with randomized temporal structures. Surrogates were then binarized around the median, and LZ76 complexity was computed. LZsN is then defined as the relative value to the mean across 100 such phase-shuffled surrogates.

$$LZsN = C/C_{phase\_mean}$$

Toker et al., 2022 found this to be the best way of normalisation to counter effects from spectral changes. However, Mediano and colleagues have argued that surrogate-based normalization can be problematic in between-condition comparisons, because differences in the spectra also change the surrogate baselines. Consequently, the difference between normalized values may reflect differences between the null models themselves, potentially producing false positives rather than isolating a purely non-spectral effect [63-65]. We therefore report both measures, to facilitate comparison with prior LZ complexity work, while basing our main inferential claim on the entropy-rate. For this study we also report average LZs (LZsum) and LZsN (LZsumN) across all channels, similar to Lewis-Healey et al., 2024 and Timmermann et al., 2023. Note we refrained from using a Hilbert-amplitude binarization because the instantaneous phase of broadband data is not well defined or interpretable [52].

#### Permutation entropy

Permutation entropy (PeEn) [66] is an entropy estimator increasingly being used to study electrophysiological signals and differentiate between altered and minimal states of consciousness [51,55,67-70]. One of PeEn's advantages is its resilience to low signal-to-noise ratios [66]. PeEn is a measure of signal irregularity that quantifies the unpredictability of the ordering of the signal pattern. First, the EEG timeseries is converted into a sequence of ordinal patterns by transforming the signal into vectors of $m$ points, separated by a delay $\tau$. This results in $m!$ number of ordinal patterns. The values of the $m$ points are ranked within each vector, and the occurrence of each $m!$ ordinal pattern is counted. For example, if the 3 points sub-segment is strictly decreasing, it will be coded as [3, 2, 1]; if it strictly increases it will be coded as [1, 2, 3]; if it decreases in the first 2 points but the last one increases above of them it will be coded as [2, 1, 3]. The relative frequency of ordinal patterns forms the probability distribution, and PeEn is computed as the Shannon entropy [71] of this distribution. Shannon entropy (and PeEn) will be high for a homogeneously distributed set, and low for a heterogeneously distributed set. The value is then normalized by $log(m!)$ to lie between 0 and 1, where higher values indicate a more diverse set of ordinal patterns (higher entropy) and lower values indicate a more repetitive signal structure. PeEn was computed using EntropyHub [72] on the broadband signal (1-35 Hz) using m = 3 and τ =4 samples (4 ms at 1000 Hz), equivalent to $\tau = 1$ at 250 Hz, implemented with a stride of 2 by retaining every other sample.

#### Spectral entropy

Spectral entropy (SpecEn) is a measure of the dispersion of spectral power across frequencies. Conceptually, it reflects the Shannon entropy of the normalized power spectral density, with higher values indicating a more homogeneous distribution of power and lower values indicating a spectrum dominated by fewer frequencies. This makes spectral entropy a useful summary measure of spectral diversity and irregularity and has been used in electrophysiological studies of altered states of consciousness [45,73-78]. SpecEn was estimated using MATLAB's spectralEntropy function [79] from the power spectrum in the 1–35 Hz range. Each epoch was analyzed as a single Hamming-tapered segment, and spectral entropy was computed from the normalized distribution of spectral power across frequency bins within this range.

#### Higuchi fractal dimension

Higuchi fractal dimension (HFD) was used to quantify the scale-invariant temporal complexity of the EEG signal. It captures how signal complexity is organized across the temporal dimension, in contrast to measures that characterize the signal's complexity via uniformness (PeEn and SpecEn) or compression (LZs). HFD, similarly to other complexity measures described here, has been used to study the complexity of the electrophysiology in various altered states of consciousness, including propofol [80,81], LSD [82], and experienced meditators [45,83]. Conceptually, HFD estimates how irregular or self-similar a time series remains when examined across progressively larger temporal intervals. For a discrete signal $x(1), x(2), \dots, x(N)$, where $x$ are samples in the timeseries, Higuchi's algorithm constructs, for each integer $k = 1, \dots, K_{\max}$, a set of $k$ subseries

$$X_m^{(k)} = \{x(m), x(m+k), x(m+2k), \dots\}, m = 1, \dots, k.$$

Where $m$ is the offset for the subseries. For a given scale $k$, the signal is represented by $k$ subseries, each defined by a different starting offset $m$. A normalized curve length $L_m(k)$ is computed for each subseries from the point-to-point absolute changes, and these values are then averaged to give the mean curve length at scale $k$:

$$L(k) = \frac{1}{k}\sum_{m=1}^{k} L_m\,(k)$$

For fractal signals, $L(k)$ decreases with increasing time scale, and this relationship is approximately linear in log-log space. HFD is therefore estimated as the slope of the linear regression of $\log L(k)$on $\log(1/k)$. Higher HFD values indicate a more irregular, complex, and scale-free temporal profile, whereas lower values indicate a smoother and less complex signal. In practice, for one-dimensional time series, HFD typically lies between 1 and 2.

The highest $k$-value, $k_{\mathrm{max}}$, was selected based on saturation analysis. For a subset of runs, we computed the mean HFD over a sweep of $k_{\mathrm{max}}$ values and chose the smallest value in the flattening region of the curve (see Supplementary Figure 15), balancing stability with computational cost. Based on this procedure, $k_{\mathrm{max}} = 200$ was used for all analyses. After selecting Kmax=200, HFD was computed using the Higuchi_FD function [84].

### Broadband detrended fluctuation analysis

Brain criticality was assessed using broadband detrended fluctuation analysis on epoched data, similar to other studies [85,86]. We quantified scale-free long-range temporal correlations (LRTC), i.e., temporal memory, in the EEG time series by estimating the DFA exponent, α [87], which is similar to the Hurst exponent. α is estimated as the slope of the linear relation between root mean square fluctuations across different timescales or window sizes of the time series. For interpretation, if $0 < \alpha < 0.5$, the signal is anticorrelated with memory; if $\alpha = 0.5$, it is indistinguishable from white noise; if $0.5 < \alpha < 1$, the signal exhibits positive correlation and has a memory. LRTC becomes stronger as α nears 1. If $1 < \alpha < 2$, the process is non-stationary [88]. The DFA computation was performed using a modified version of DFluC (Colombo et al., 2016) that supported custom window sizes. In short, broadband DFA (1-35 Hz) was estimated by splitting the epochs into non-overlapping windows with log-spaced lengths from 100 samples to 1000 samples. Within each window, the second-order polynomial trend was removed, and the fluctuation quantified as the root mean square of the residuals pooled across all windows. This was repeated for all 50 window scales. The power-law relationship between fluctuations and window sizes is then estimated via linear regression on a log-log plot, with the slope being equal to the DFA exponent α. Window sizes for broadband DFA were chosen based on guidance from previous research [89], setting the minimum window sizes as $n_{min}=f_s/f_{max}$ and maximum window sizes as $n_{max}=f_s/f_{min}$. With $f_s$=1000 Hz and a 1-35 Hz band yielding $n_{min}$=28.6 samples (29 ms) and $n_{max}$=1000 samples (1.0 s). Previously, small-scale curvature from possible filter artifacts has been observed [89,90]. We investigated our choice of window sizes by visual inspection of log-log plots, assessing linearity in the data for the given box sizes using the first epoch for all ACAM-J1-8 and control runs across various electrodes, finding 100 samples as the minimum most suitable (see Supplementary Figure 16). This confirmed our window size choices. We chose to analyse 10-second epochs, as this provides 10 boxes at the largest scale and therefore a reasonable number of windows available for averaging, as recommended in Hardstone et al., 2012.

#### Brain-phenomenology analyses

Relationships between the global EEG measures and phenomenology were examined with partial least squares correlation (PLSC), a multivariate method that extracts latent variables (LVs) capturing patterns of covariance between two sets of variables [93-95]. PLSC was preferred over mass-univariate correlation because the EEG measures are strongly intercorrelated and because the analysis targets overarching brain-experience patterns rather than individual variable pairs, while avoiding the multiple-comparison problem.

#### Stability analysis

As a supplementary check of the chosen minimum epoch threshold, we examined the stability of global total, oscillatory, and aperiodic power, LZsum, LZsumN, PeEn, SpecEn, HFD, and DFA as a function of the number of clean epochs included. For each measure and condition we estimated single-epoch reliability with a two-way random-effects, absolute-agreement, single-measurement intraclass correlation, ICC(2,1), which models epoch position (temporal order) as an occasion factor and is the form recommended for repeated-measures data [97], on the participant x epoch matrix of single-epoch values (unequal epoch counts handled with the standard mean-group-size correction; 95% confidence intervals by the F-distribution method [98]), and projected the reliability of a $K$-epoch average with the Spearman–Brown formula, $R_K = K \cdot \mathrm{ICC1} / (1 + (K-1) \cdot \mathrm{ICC1})$. This quantifies the test–retest reliability of the epoch-averaged measure, how reproducibly it is estimated as the number of epochs increases, computed separately within each condition. Reliability was interpreted with the benchmarks: poor (<0.50), moderate (0.50-0.75), good (0.75-0.90), excellent (>0.90) [99].

As an additional analysis, and because the main analyses rest on the within-participant jhana-minus-control difference, we estimated the reliability of that contrast as a function of the number of epochs per condition, using a non-overlapping split-half procedure. For each participant we pooled the ACAM-J epochs and the combined control (memory + counting) epochs, drew $K$ epochs of each into two disjoint halves, formed the contrast $\Delta$ = mean (ACAM-J) − mean(control) from each half, and computed the correlation of the two half-estimates across participants (60 resamples averaged). In plain terms, each participant's epochs are repeatedly split into two independent halves, the effect is estimated separately from each half, and reliability is how well the two estimates agree, improving as more epochs enter each half. This split-half logic is the classical basis of the Spearman-Brown projection and the standard way to establish how many trials or epochs a neural measure requires [100,101].

For the qualitative stability analyses, LZsum, LZsumN, PeEn, SpecEn, and HFD, only runs with at least 15 clean epochs were included. For each epoch count from 1 to 20, clean epochs were randomly sampled without replacement within that count, and a global metric was computed by averaging across the selected epochs and channels. If multiple runs were available for a subject within a condition, run-level values were averaged to yield a single subject-level estimate. Group-level plots display the mean, standard deviation, and individual subject values across epoch counts, whereas subject-level plots show the same procedure within each subject relative to that subject's full-data estimate. For each condition, the dashed line indicates the estimate obtained using all available clean epochs from eligible runs and thus serves as a full-data reference against which subsampled estimates at each epoch count were compared. This analysis was used as a qualitative assessment similar to previous investigations [102] of whether estimates appeared to stabilize at the 15-epoch inclusion threshold used in the main analyses. An analogous stability analysis was performed for IRASA-derived spectral power measures and DFA. Because these analyses were based on longer epochs, clean epochs were subsampled from 1 to 10 epochs using 5-s segments for IRASA and 10-s segments for DFA. These analyses were used to qualitatively assess whether the minimum inclusion thresholds of 6 epochs for IRASA and 3 epochs for DFA were sufficient for stable estimation.

### Supplementary Data

**Supplementary Data 1. Linear mixed-effects model statistics for global EEG measures.** For each global power and complexity measure: estimates, standard errors, 95% confidence intervals, Satterthwaite degrees of freedom, $t$ values, uncorrected and Benjamini–Hochberg FDR-corrected $P$ values, and Cohen's $d$ with 95% confidence intervals. These cover each

ACAM-J state versus the combined control and counting versus memory, together with omnibus $F$ tests of state.

**Supplementary Data 2. Cluster-based permutation test statistics for topographic analyses.** For each contrast (ACAM-J1–8 versus the combined control, and memory versus counting) across power, aperiodic and complexity measures: $n$, number of permutations, minimum cluster $P$, FDR $q$ across the eight ACAM-J states, and for each cluster its direction, size, cluster statistic, peak $t$, peak electrode, approximate location and member electrodes. Analysis parameters are also included.

### Power topography of memory vs counting

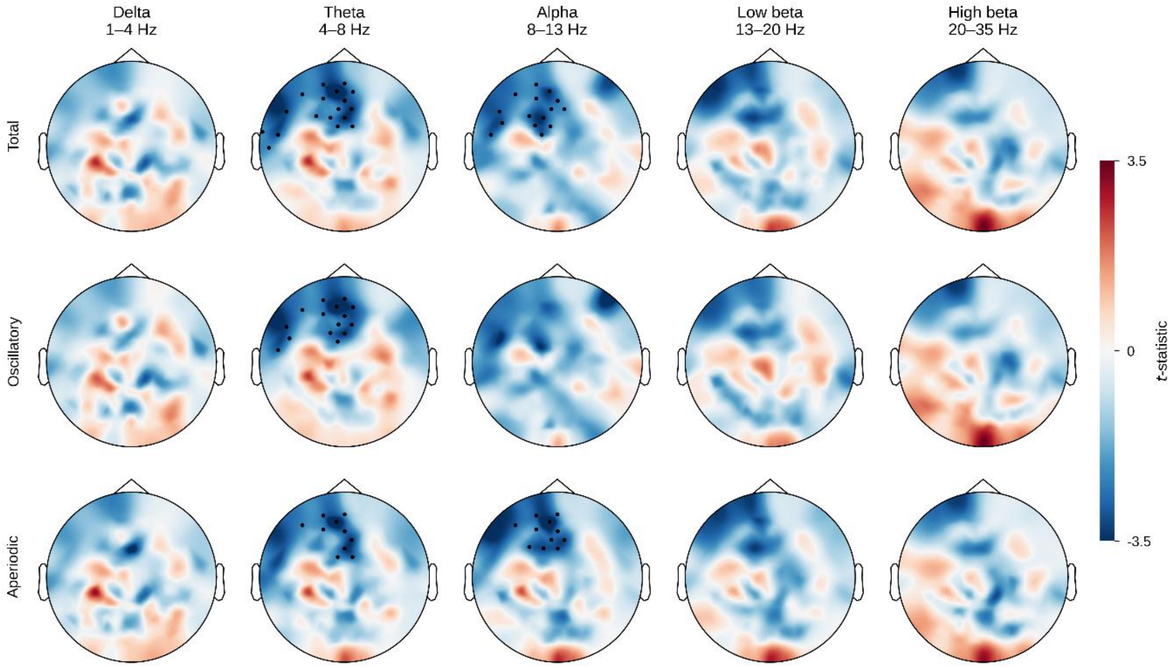

***Supplementary Figure 1. Topography of memory versus counting.*** *Paired t-statistics for memory minus counting in the total, oscillatory and aperiodic IRASA components (rows) across frequency bands (columns). Red indicates higher power during memory. Markers show electrodes in clusters with P < 0.025 (two-sided cluster-based permutation test as in Fig. 1). No FDR correction was applied because there is one contrast per band. n = 20. Cluster statistics are given in Supplementary Data 2.*

### IRASA separation plots

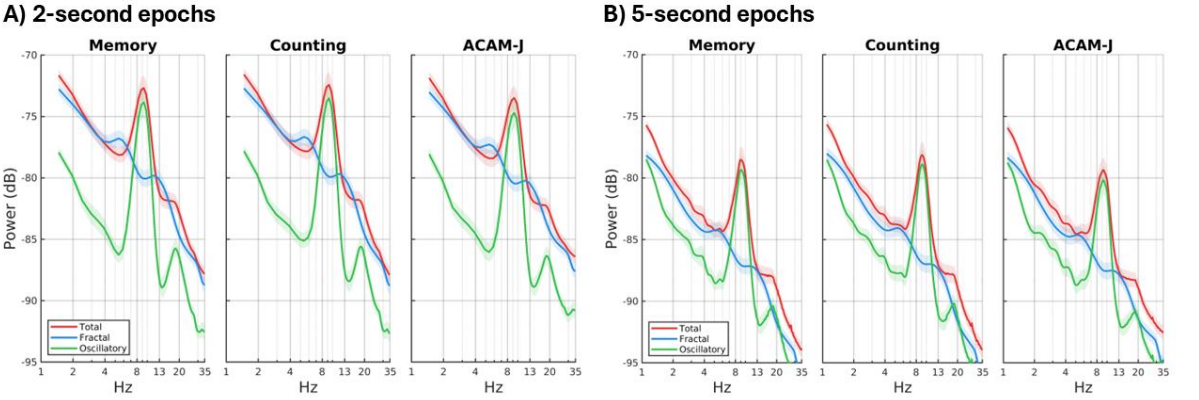

***Supplementary Figure 2.*** *Comparison of IRASA-derived spectral decomposition across epoch lengths. (A) Power spectra computed from 2-second epochs and (B) power spectra computed from 5-second epochs for the Memory, Counting, and ACAM-J conditions. Red lines show the total power spectrum, blue lines show the fractal (aperiodic) component, and green lines show the oscillatory component after IRASA-based separation. Spectra were averaged across channels and participants and are plotted in dB over 1–35 Hz on a logarithmic frequency axis. Shaded areas indicate the standard error of the mean (SEM). The ACAM-J panel represents the pooled all-ACAM-J composite rather than a simple equal-weight average of the eight ACAM-J states*

### Spectral total, oscillatory, and aperiodic power using 2-s epochs

#### Total spectral power (2-s epoch)

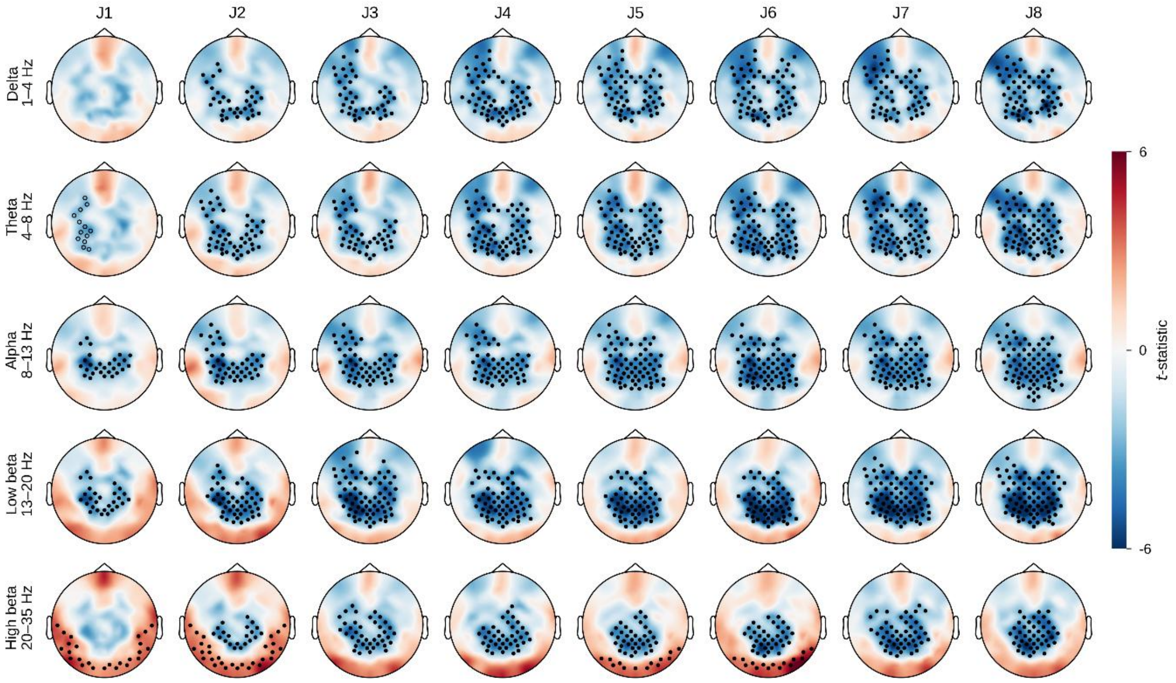

***Supplementary Figure 3. Topography of total spectral power from 2-s epochs.*** *Channel-wise paired t-statistics for each ACAM-J state minus the combined control condition (mean of memory and counting), for the total IRASA component estimated from 2-s epochs. Rows show frequency bands and columns show ACAM-J1 to ACAM-J8. Red indicates higher and*

*blue lower power during ACAM-J; the colour scale is shared across all maps. Markers show electrodes in significant clusters from a two-sided cluster-based permutation test (cluster-forming threshold P < 0.05; cluster-level P < 0.025; exact null distribution from all sign-flip permutations). Markers are drawn only for contrasts that also survived FDR correction across the eight ACAM-J states within each band (q < 0.025). Filled markers show q < 0.01 and open markers show 0.01 ≤ q < 0.025. n = 19, 20, 20, 20, 20, 19, 18 and 17 for ACAM-J1–8.*

### Oscillatory spectral power (2-s epoch)

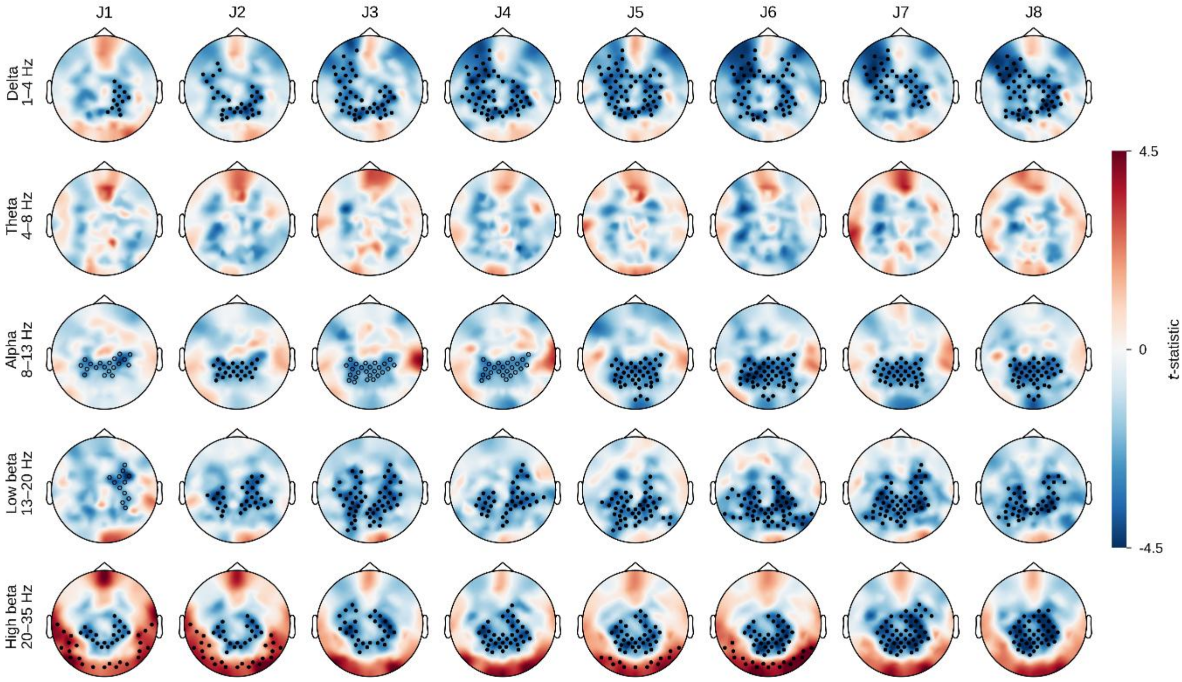

***Supplementary Figure 4. Topography of oscillatory spectral power from 2-s epochs.** As in Supplementary Figure 9, for the oscillatory IRASA component.*

### Aperiodic spectral power (2-s epoch)

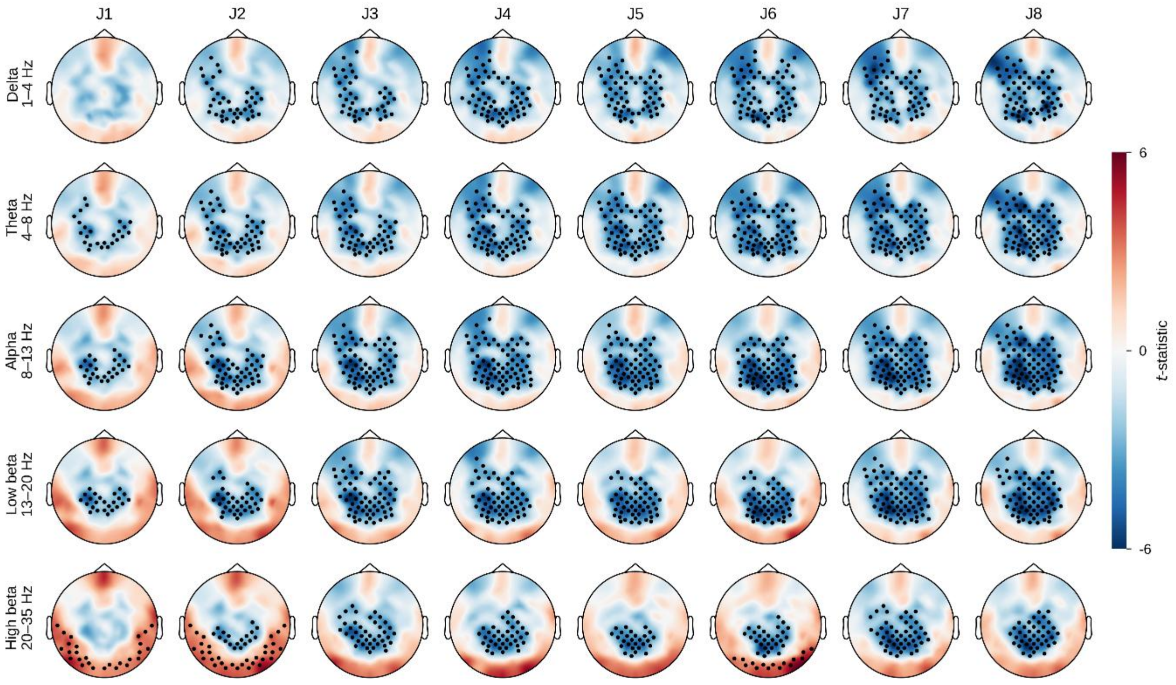

***Supplementary Figure 5. Topography of aperiodic spectral power from 2-s epochs.*** *As in Supplementary Figure 9, for the aperiodic IRASA component.*

### Total spectral power (5-s epoch)

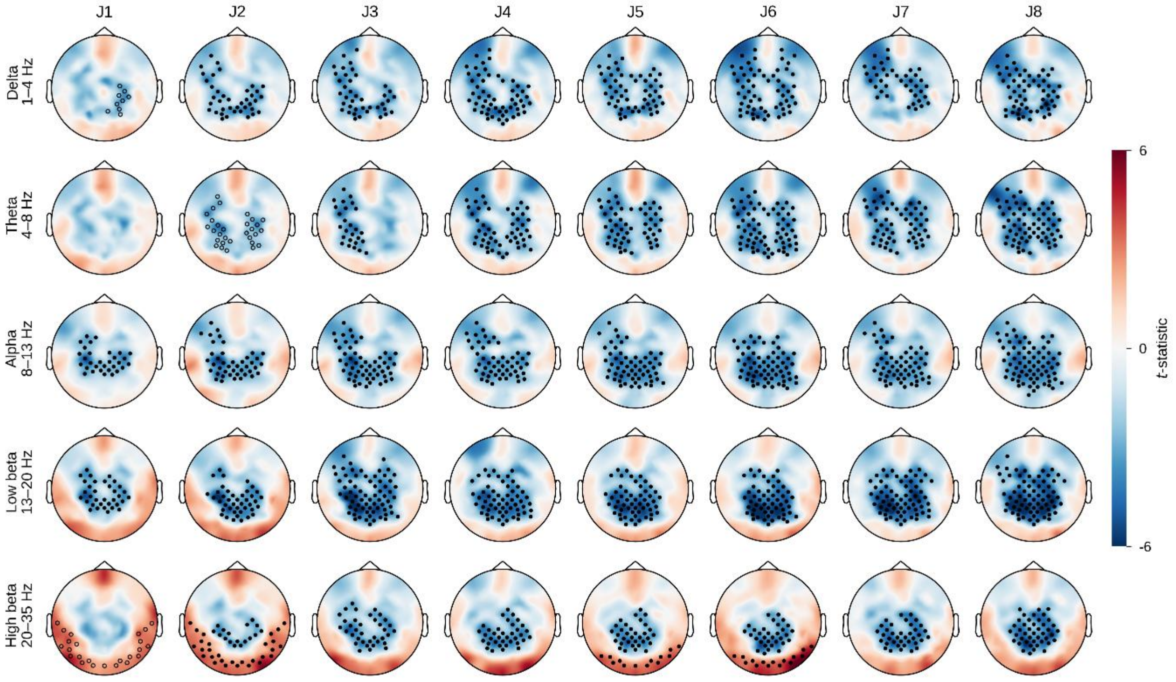

***Supplementary Figure 6.*** *Topography of total spectral power during ACAM-J1–8. Scalp maps show channel-wise paired t-statistics for each ACAM-J state minus the combined control condition (mean of memory and counting), for the total IRASA component estimated from 5-s epochs. Rows show frequency bands and columns show ACAM-J1 to ACAM-J8. Red indicates higher and blue lower power during ACAM-J; the colour scale is shared across all maps. Markers show electrodes in significant clusters from a two-sided cluster-based permutation test (cluster-forming threshold $P < 0.05$; cluster-level $P < 0.025$; exact null distribution from all sign-flip permutations). Markers are drawn only for contrasts that also survived Benjamini–Hochberg FDR correction across the eight ACAM-J states within each band ($q < 0.025$). Filled markers show $q < 0.01$ and open markers show $0.01 \leq q < 0.025$. n per state is given in Table 1. Cluster statistics are given in Supplementary Data 2.*

### Full spectrum activity (5-s epochs)

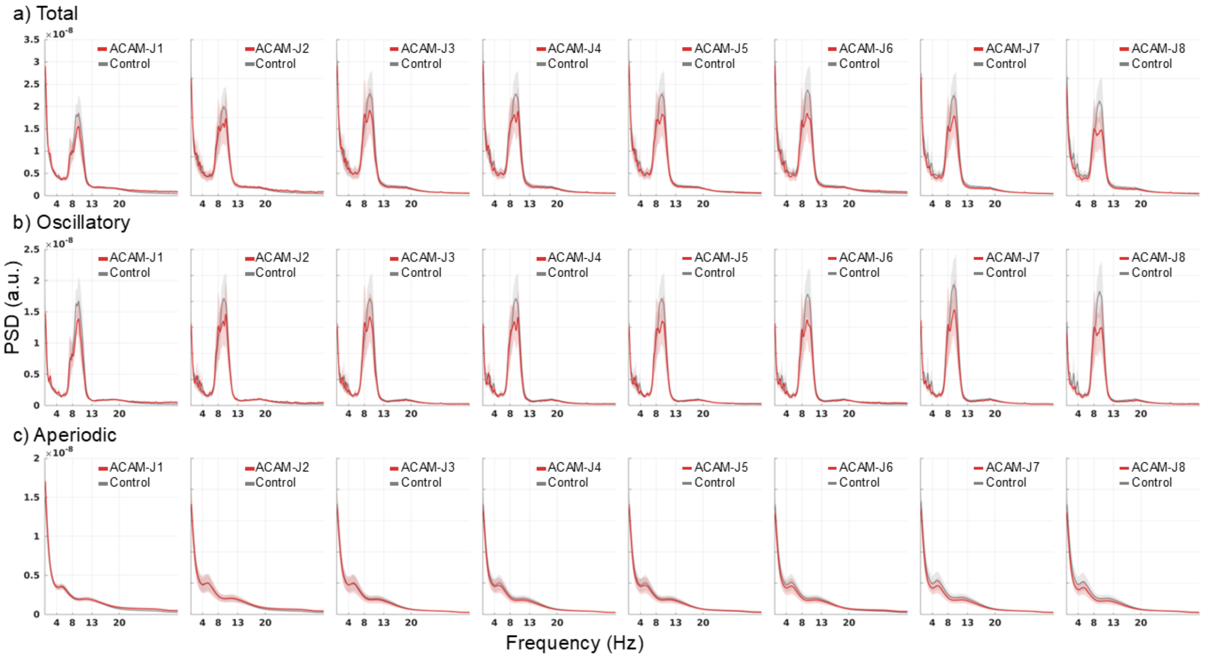

***Supplementary Figure** 7. **Spectral activity across power categories and ACAM-J**. Group-average IRASA power spectra (1-35 Hz) for each jhana state compared with the combined control condition. Columns show ACAM-J1 to ACAM-J8. Rows show the three IRASA components: **a**, total power, **b**, oscillatory power, and **c**, aperiodic power. Spectra were averaged across channels within subject and then across subjects, shaded bands indicate SEM. n per state is given in Table 1.*

### Post-hoc EMG confound analyses

After the ACAM-J scan, participants were asked to rate the following question for ACAM-J1 and ACAM-J2 using a 1–10 scale: "Automatic movements include twitching, rocking, swaying, unintentional movements, or switches in the position of the body (such as the head spontaneously moving upwards or downwards)"

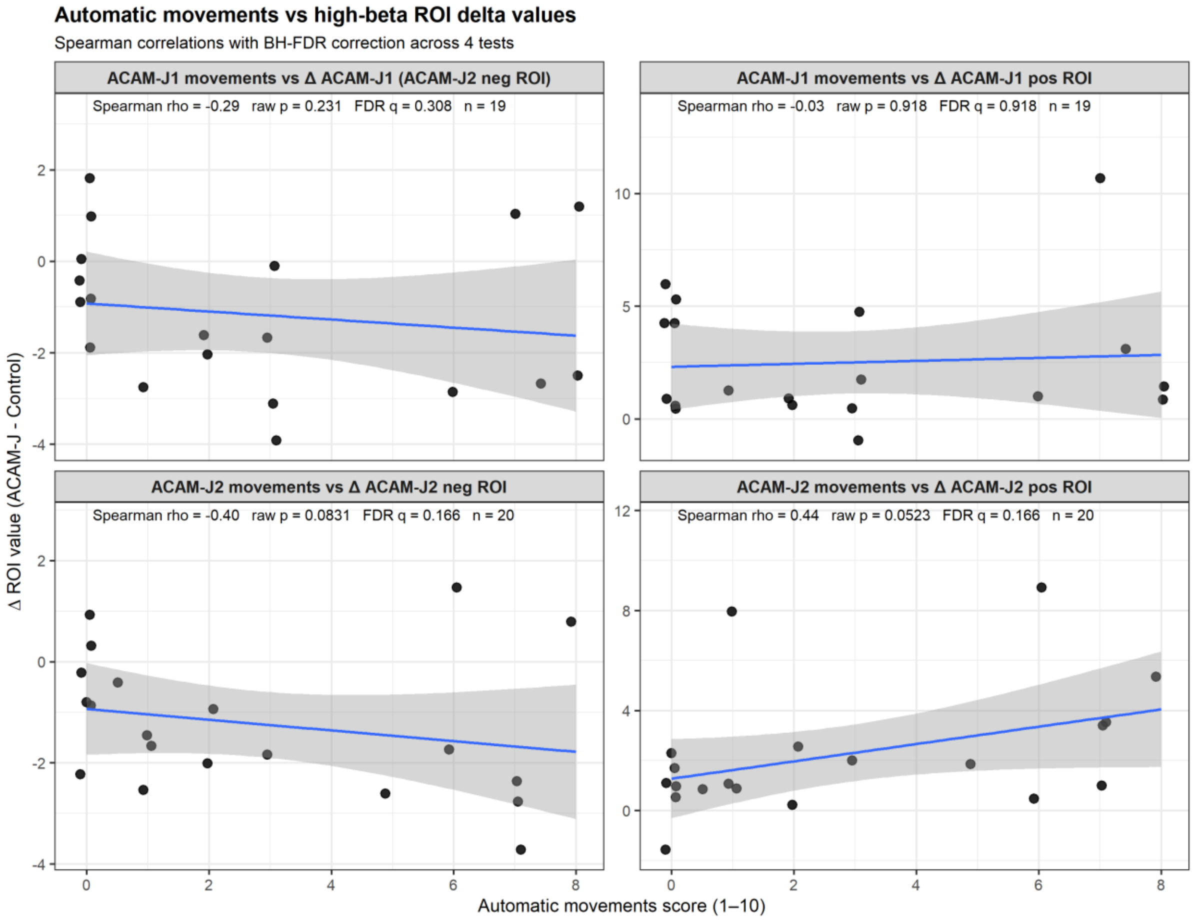

***Supplementary Figure 8.*** *Scatterplots showing associations between automatic movement ratings and control-relative changes in high-beta ROI values for ACAM-J1 and ACAM-J2. ROI change scores were calculated as Δ = ACAM-J − Control, using high-beta power derived from 2-s epochs to match the exact epochs used for the entropy measures. Panels show correlations between automatic movement ratings and the relevant positive and negative high-beta ROIs for ACAM-J1 and ACAM-J2. Points represent individual subjects, and lines indicate least-squares fits with shaded 95% confidence intervals for visualization. Associations were tested using Spearman correlations, with FDR correction applied across the four tests.*

### Global total power

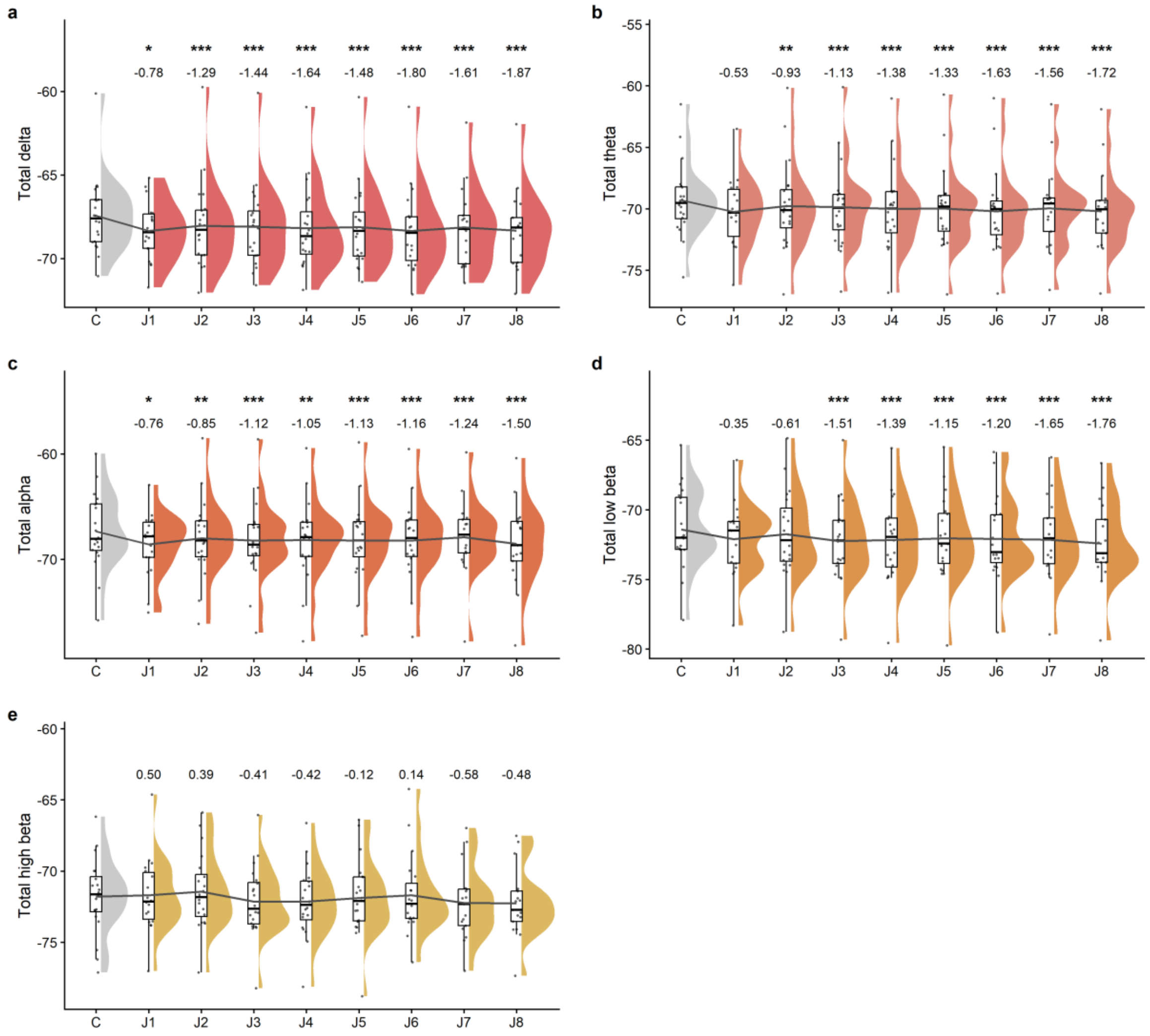

***Supplementary Figure 9.*** *Raincloud plots of total delta, theta, alpha, low beta, and high beta power (5-s epoch) across the combined control condition and ACAM-J1–ACAM-J8. Points represent individual subjects; half-violin densities, boxplots, and the grey mean line summarize the distribution within each state. Asterisks indicate BH-FDR-corrected contrasts versus the combined control condition (* p<0.05, ** p<0.01, *** p<0.001), and d values denote model-based effect sizes from the linear mixed-effects model. n per state is given in Table 1. Model statistics are given in Supplementary Data 1.*

### Global oscillatory power

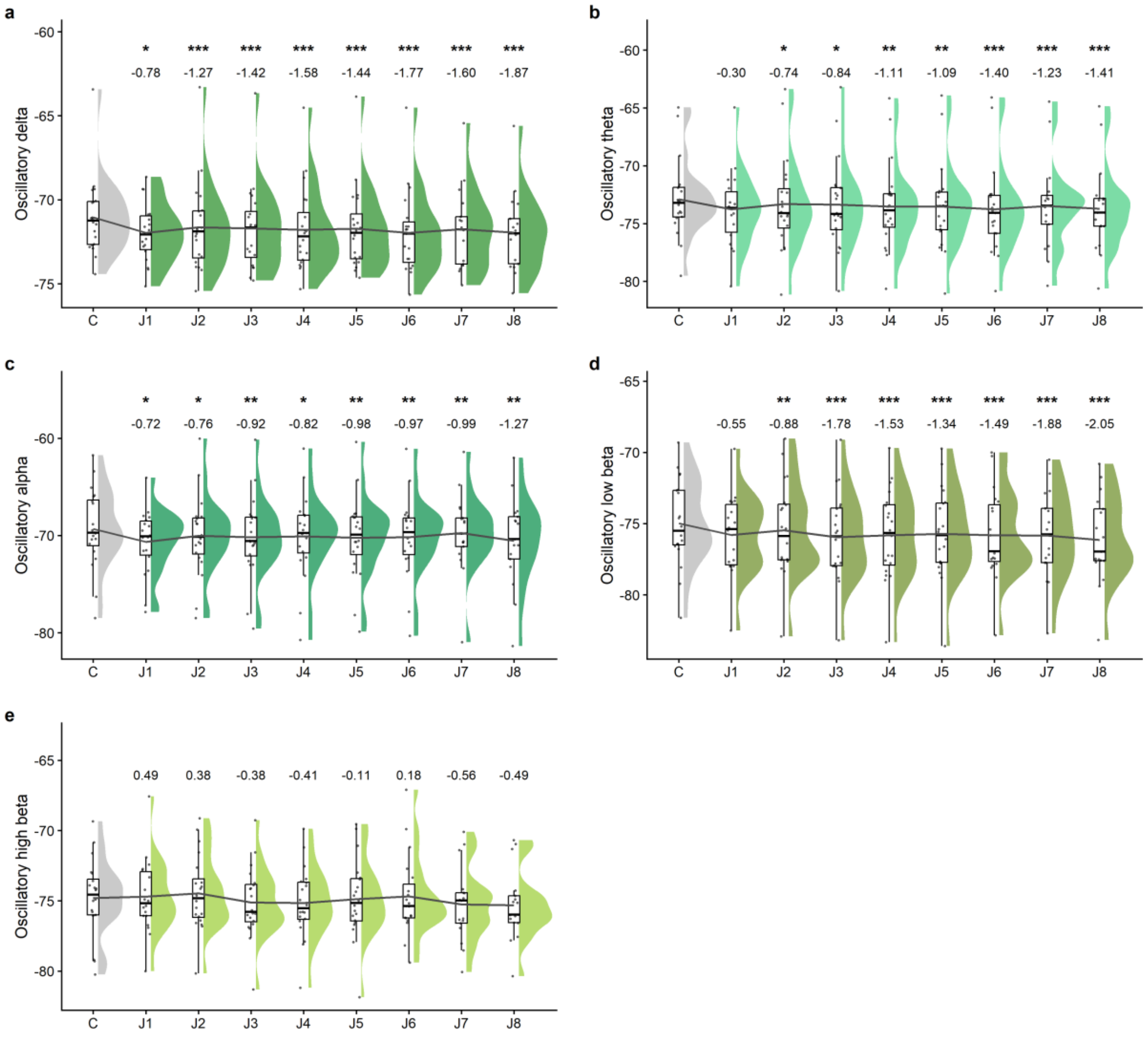

***Supplementary Figure 10.*** *Raincloud plots of oscillatory delta, theta, alpha, low beta, and high beta power (5-s epoch) across the combined control condition and ACAM-J1–ACAM-J8. Points represent individual subjects; half-violin densities, boxplots, and the grey mean line summarize the distribution within each state. Asterisks indicate BH-FDR-corrected contrasts versus the combined control condition (* p<0.05, ** p<0.01, *** p<0.001), and d values denote model-based effect sizes from the linear mixed-effects model. n per state is given in Table 1. Model statistics are given in Supplementary Data 1.*

### Global aperiodic power

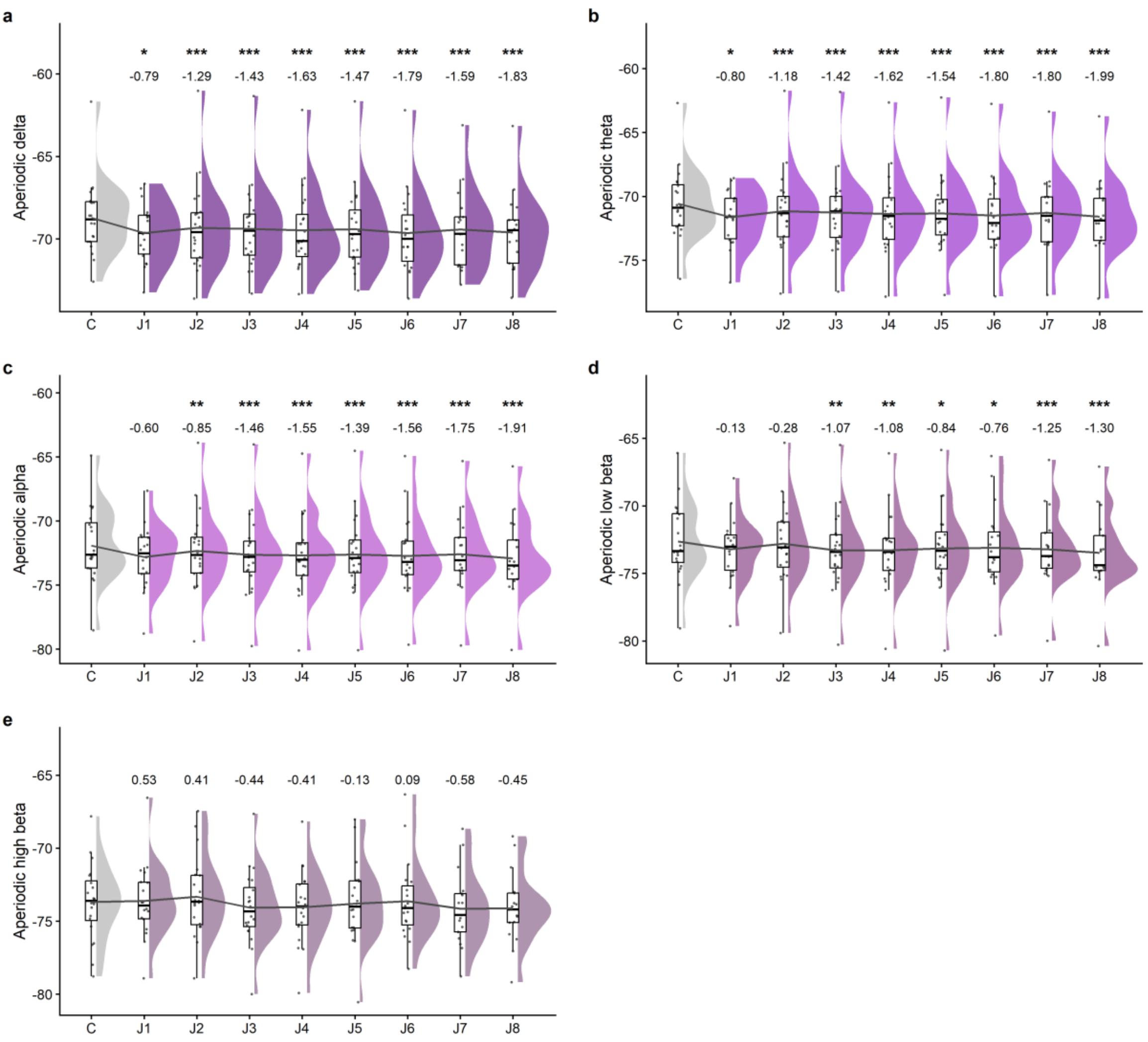

***Supplementary Figure 11.*** *Raincloud plots of aperiodic delta, theta, alpha, low beta, and high beta power (5-s epoch) across the combined control condition and ACAM-J1–ACAM-J8. Points represent individual subjects; half-violin densities, boxplots, and the grey mean line summarize the distribution within each state. Asterisks indicate BH-FDR-corrected contrasts versus the combined control condition (* p<0.05, ** p<0.01, *** p<0.001), and d values denote model-based effect sizes from the linear mixed-effects model. n per state is given in Table 1. Model statistics are given in Supplementary Data 1.*

### Extended exploration of aperiodic measures (5s epoch)

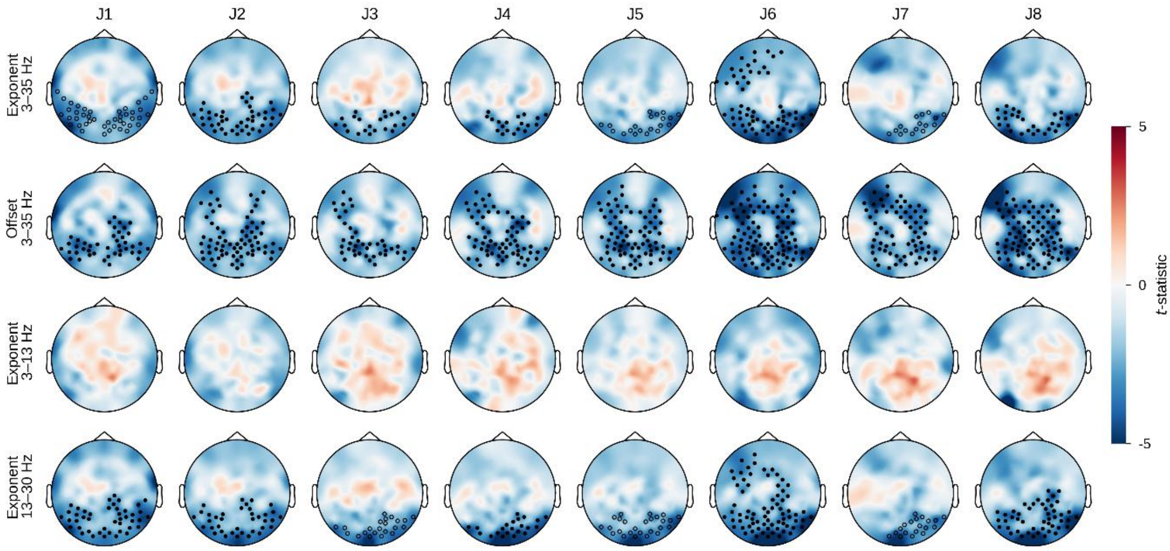

***Supplementary Figure 12. Topography of the aperiodic exponent and offset, including band-limited exponents.*** *Paired t-statistics for each ACAM-J state minus the combined control condition for the aperiodic exponent (3–35 Hz), the offset (3–35 Hz), and the exponent fitted over 3–13 Hz and over 13–30 Hz (rows), across ACAM-J1 to ACAM-J8 (columns), estimated from 5-s epochs. Negative values indicate a flatter spectrum (lower exponent) or lower broadband aperiodic power (lower offset) during ACAM-J. Markers as in Supplementary Figure 9, with FDR correction across the eight ACAM-J states within each measure. n per state is given in Table 1 of the main text.*

### ACAM-J specific between-measure correlation

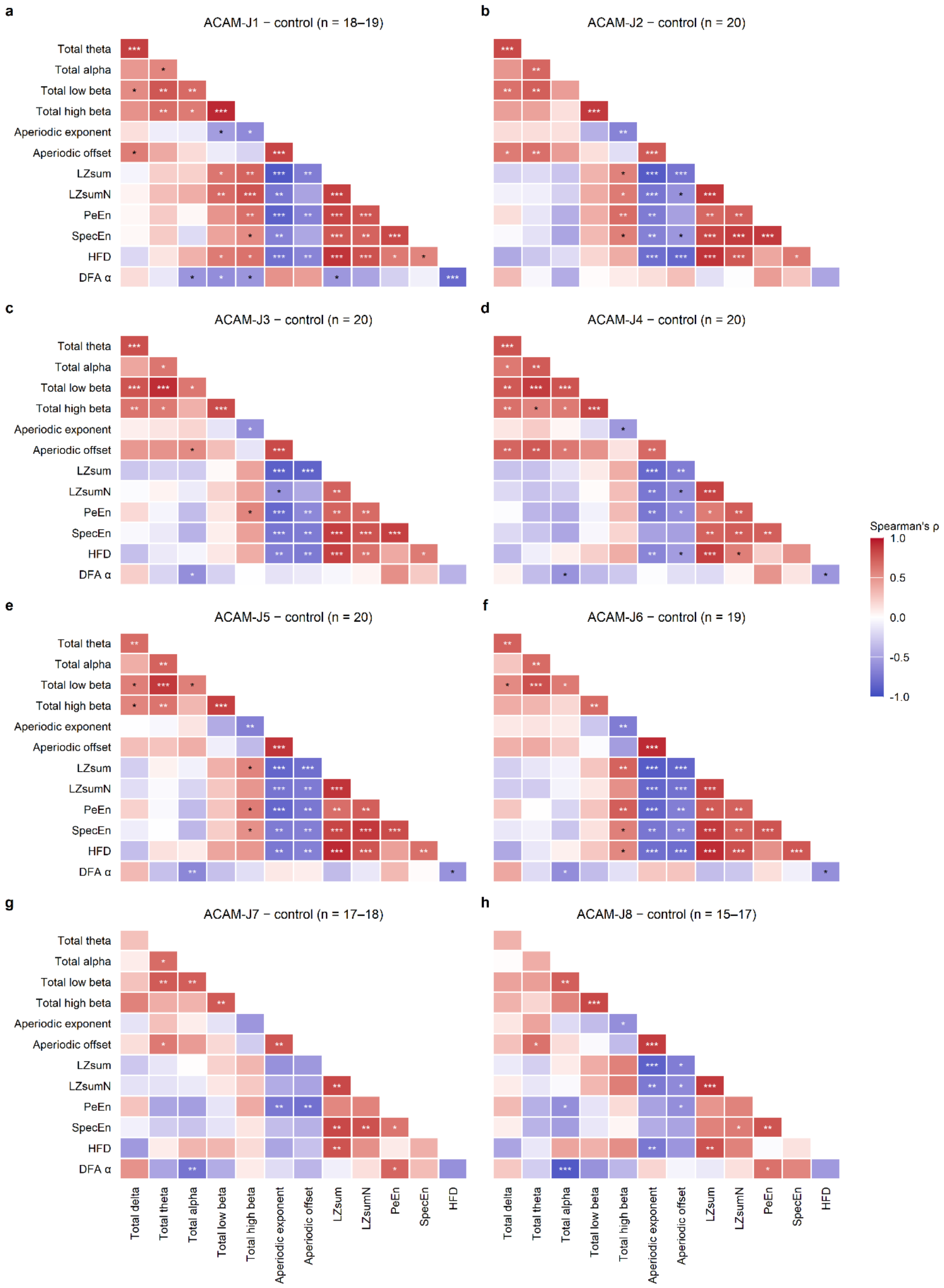

***Supplementary Figure 13. Between-measure correlations of ACAM-J-induced change for each ACAM-J state.*** *a–h, Spearman correlations across participants between changes in global EEG measures for ACAM-J1 (a) to ACAM-J8 (h). Each change is the ACAM-J state minus the combined control condition (mean of memory and counting), computed separately for each participant and measure. Power measures and the aperiodic exponent and offset were estimated from 5-s epochs, LZsum, LZsumN, PeEn, SpecEn and HFD from 2-s epochs, and DFA α from 10-s epochs. Red indicates positive and blue negative correlations; the colour scale is shared across panels. Asterisks mark correlations surviving Benjamini–Hochberg FDR correction across the 78 unique pairs within each matrix (*q < 0.05, **q < 0.01, ***q < 0.001). Correlations were computed on all participants with data for both measures, so n varies slightly between pairs (range given in each panel title). The matrix for the mean across ACAM-J states is shown in Fig. 7.*

### ACAM-J–specific factors and run-level quality

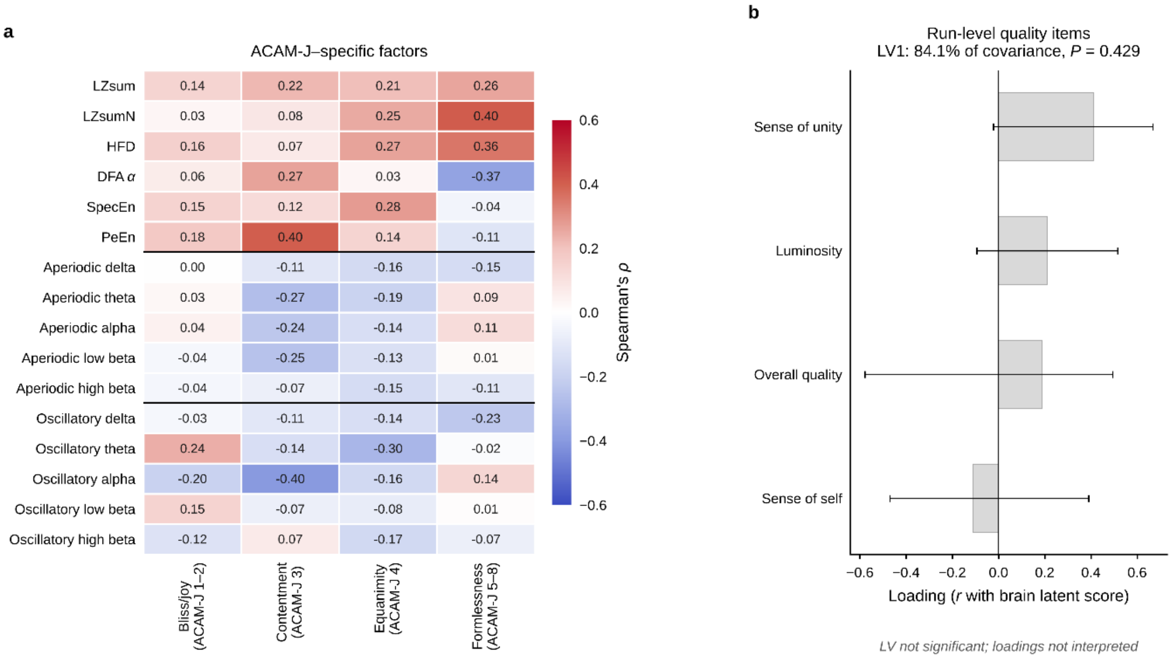

***Supplementary Figure 14. ACAM-J–specific factors and run-level quality. a)*** *Spearman rank-order correlations between the sixteen EEG measures and the four ACAM-J–specific factors. Both the scores and the EEG measures were averaged across the states for which each factor is scored, so that each participant contributed one observation (n = 20). Cell values give the correlation coefficient. No correlation survived FDR correction. Horizontal rules separate the complexity, aperiodic, and oscillatory blocks.* ***b)*** *Phenomenological loadings with 95% bootstrap confidence intervals for the first latent variable of a separate PLSC relating the run-level quality items to EEG measures averaged across the eight ACAM-J states (n = 20). This latent variable was not significant.*

### HFD saturation plot examples

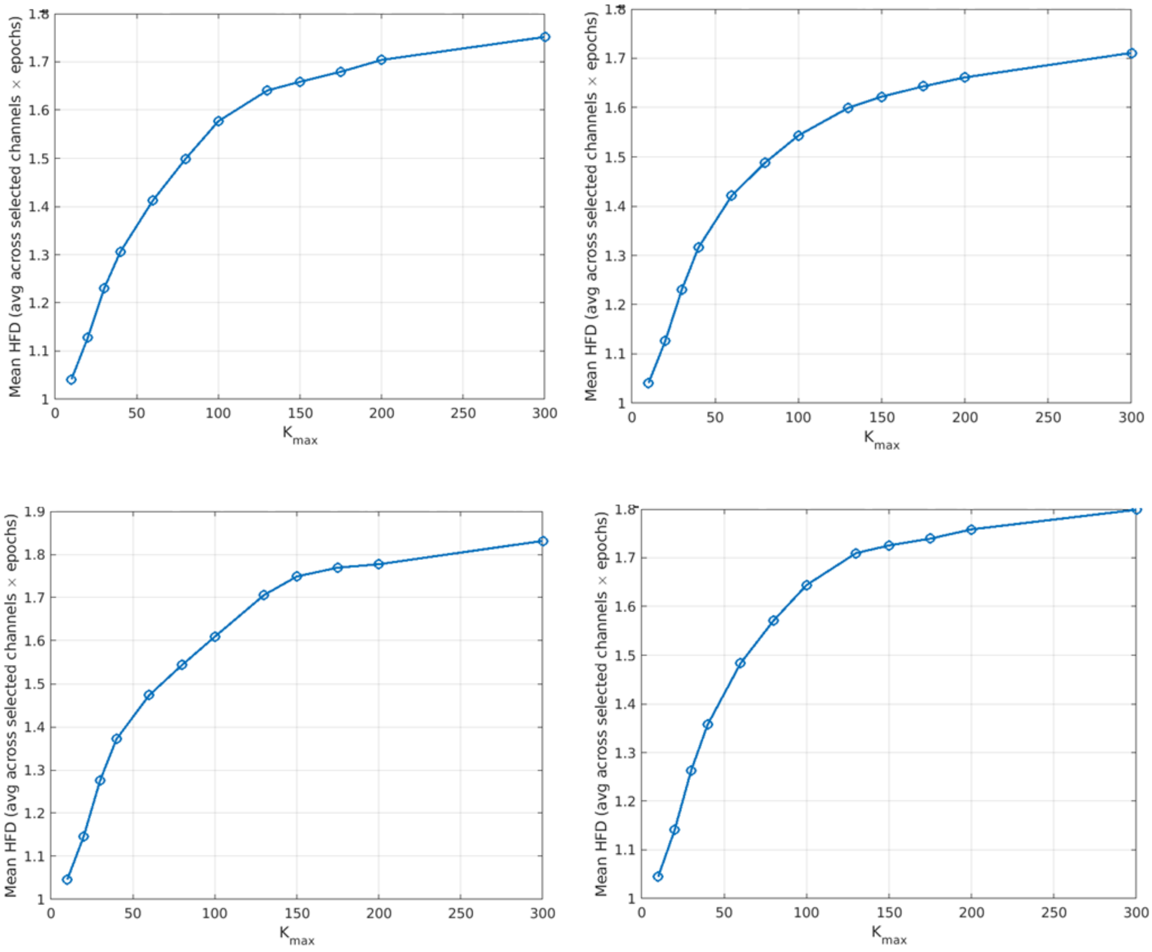

***Supplementary Figure 15****. Run-wise saturation curves for Higuchi fractal dimension (HFD) across K_max values. Each panel shows the mean HFD estimated for a single epoched EEG run as a function of K_max. For each run, HFD was computed for randomly selected epoch–channel pairs (up to 20 epochs and 32 channels), and values were then averaged to obtain one mean estimate per K_max. This analysis was used as a diagnostic assessment of the stability of the HFD estimate across increasing K_max values and to identify the range in which the estimate approached a plateau. Tested K_max values were 10, 20, 30, 40, 60, 80, 100, 130, 150, 175, 200, and 300.*

### DFA box size

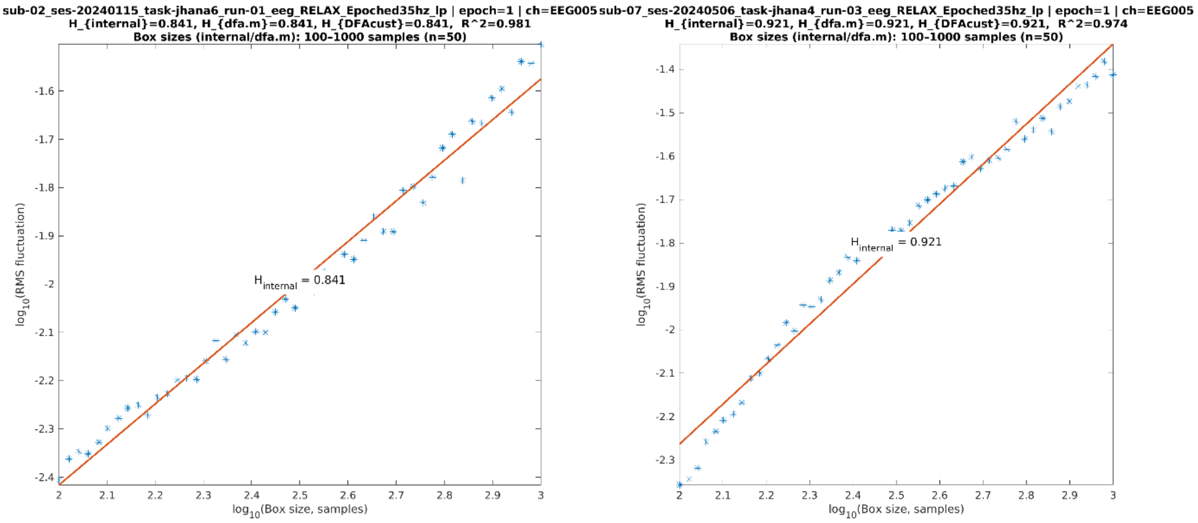

***Supplementary Figure 16.*** *Examples of log-log plots, assessing linearity in the data for the given box sizes for DFA estimation.*

### Quantitative epoch-stability analyses

#### Methods

For every measure, we quantified how its reliability grows with the number of epochs averaged and confirmed that the epoch thresholds used for inclusion (6 epochs for spectral power, 3 for detrended fluctuation analysis, 15 for the complexity measures) yield reliable estimates. Two complementary reliabilities were computed, matched to the two kinds of inference drawn in the paper: a between-subject reliability, relevant to the planned correlations of the neural measures with phenomenology, and a within-subject reliability of the ACAM-J vs combined-control contrast, relevant to the linear mixed model and cluster-based permutation analyses. Unless stated, analyses used the global value per epoch (mean across the 128 channels). Computations were performed in Python (NumPy, SciPy, h5py).

##### Epoch-stability reliability (ICC(2,1))

For each measure and condition we estimated single-epoch reliability with a two-way random-effects, absolute-agreement, single-measurement intraclass correlation, ICC(2,1), which models epoch position (temporal order) as an occasion factor and is the form recommended for repeated-measures data [97], on the participant x epoch matrix of single-epoch values (unequal epoch counts handled with the standard mean-group-size correction; 95% confidence intervals by the F-distribution method [98]), and projected the reliability of a $K$-epoch average with the Spearman–Brown formula, $R_K = K \cdot \text{ICC1} / (1 + (K-1) \cdot \text{ICC1})$. This quantifies the test–retest reliability of the epoch-averaged measure — how reproducibly it is estimated as the number of epochs increases — computed separately within each condition. Reliability was interpreted with the benchmarks: poor (<0.50), moderate (0.50–0.75), good (0.75–0.90), excellent (>0.90) [99])

##### Additional analysis: reliability of the ACAM-J vs combined-control contrast

As an additional analysis, and because the main analyses rest on the within-participant jhana-minus-control difference, we estimated the reliability of that contrast as a function of the number of epochs per condition, using a non-overlapping split-half procedure. For each participant we pooled the ACAM-J epochs and the combined control (memory + counting) epochs, drew $K$ epochs of each into two disjoint halves, formed the contrast $\Delta$ = mean (ACAM-J) − mean(control) from each half, and computed the correlation of the two half-estimates across participants (60 resamples averaged). In plain terms, each participant's epochs are repeatedly split into two independent halves, the effect is estimated separately from each half, and reliability is how well the two estimates agree, improving as more epochs enter each half. This split-half logic is the classical basis of the Spearman-Brown projection and the standard way to establish how many trials or epochs a neural measure requires [100,101].

#### Results

##### Epoch-stability reliability

At the inclusion thresholds, the epoch-averaged estimate was reliable for spectral power and for most complexity measures. Power at $K = 6$ was excellent in the theta, alpha, low-beta and high-beta bands ($R$ = 0.94–0.99) across total, aperiodic and oscillatory components, and good for delta (lowest $R$ = 0.82; Supplementary Figure 17-18). Complexity at $K = 15$ was excellent for spectral entropy ($R$ = 0.90–0.96), LZsum (0.95–0.98), permutation entropy (0.97–0.98) and Higuchi fractal dimension (0.96–0.98), and DFA at $K = 3$ was good-to-excellent (0.89–0.93; Supplementary Figures 19 and 20). The phase-normalised LZsumN was the single exception. Its single-epoch reliability was very low (ICC1 ≈ 0.08–0.14) so that even at 15 epochs $R$ reached only ≈ 0.55–0.71. Across all 231 measure × condition cells, 95% met the good and 93% the excellent criterion, with the sub-threshold cells confined to LZsumN.

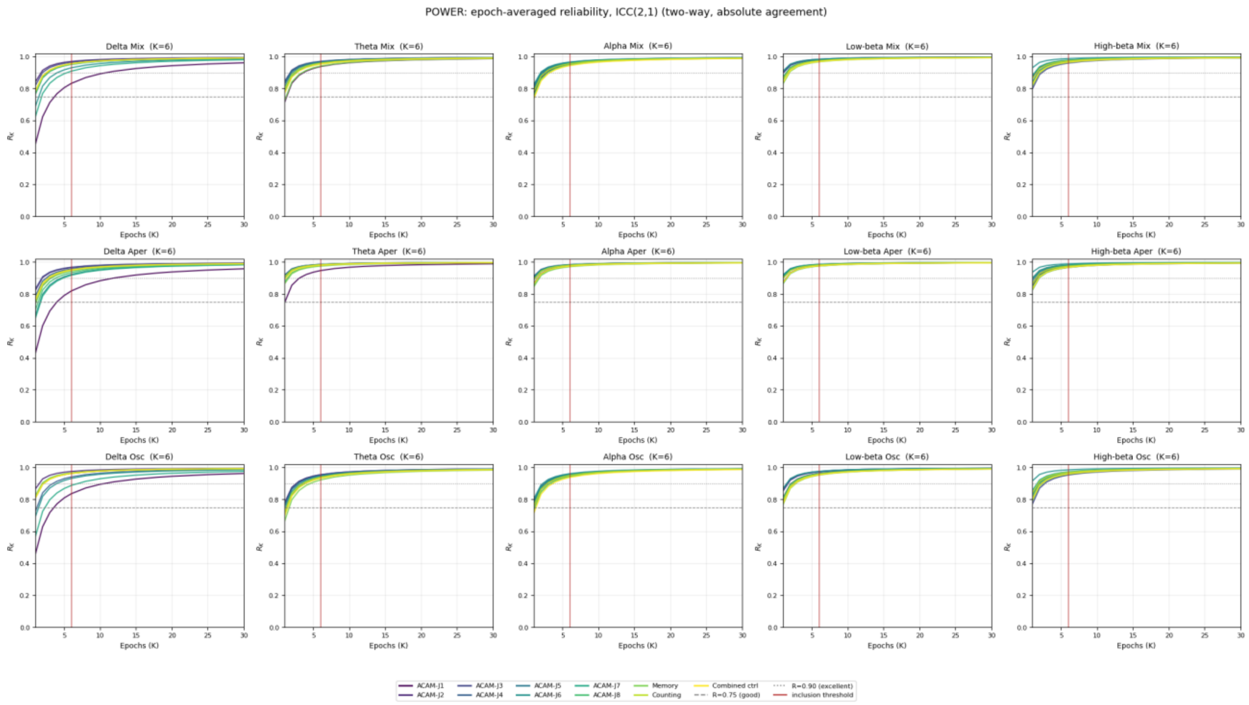

**Supplementary Figure 17.** Between-subject reliability of spectral power versus number of epochs averaged. Reliability RK (Spearman–Brown projection of the single-epoch ICC(2,1)) for K = 1–30, for IRASA total, aperiodic and oscillatory power in five bands; each coloured line is one of the 11 conditions (ACAM-J1-8, memory, counting, combined control). Dashed and dotted lines mark 0.75 (good) and 0.90 (excellent), the vertical line marks the six-epoch inclusion threshold.

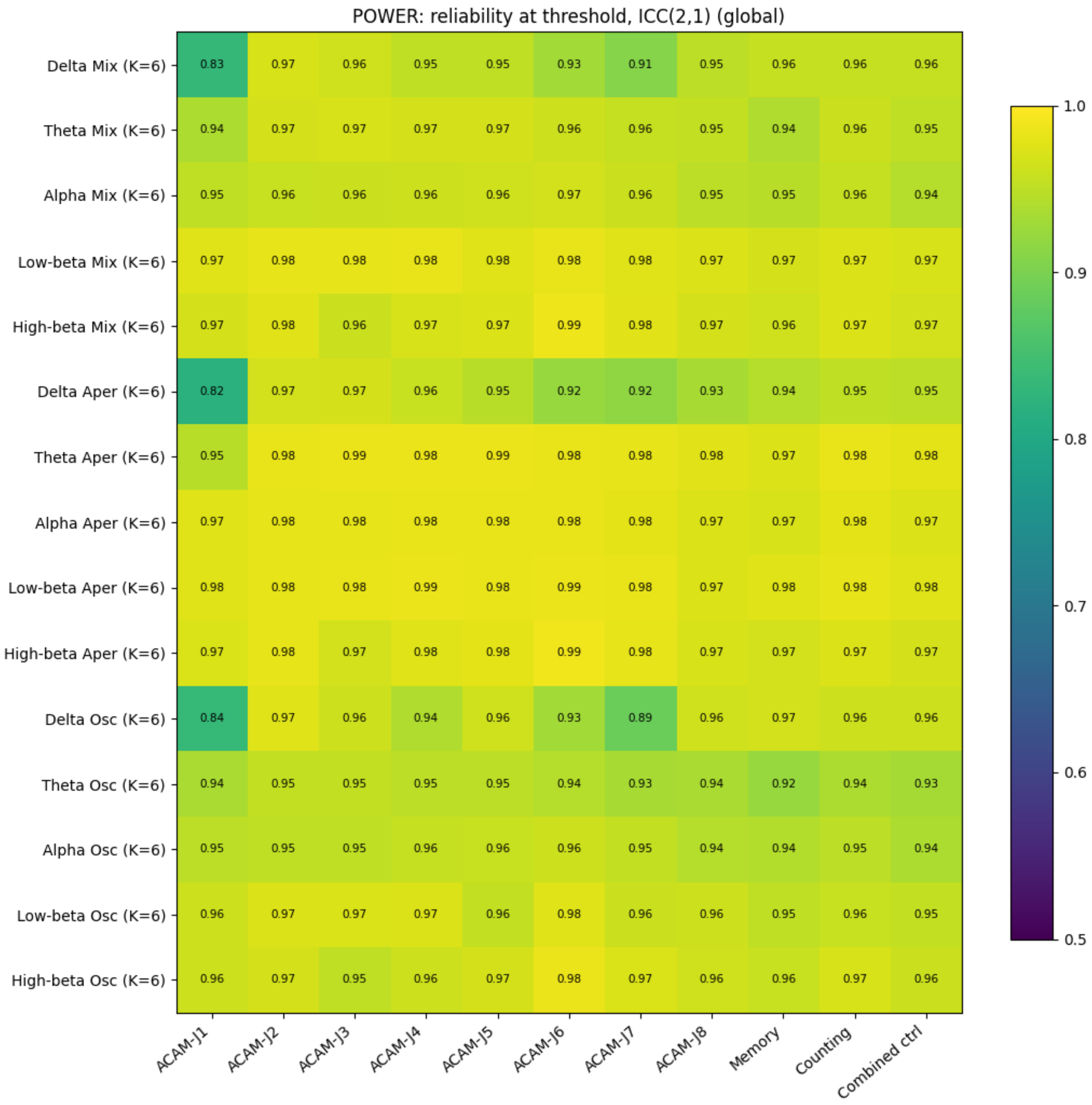

**Supplementary Figure 18.** Between-subject reliability of spectral power at the six-epoch threshold, for every power measure × condition (global 128-channel-mean value, annotated per cell). Reliability was ≥ 0.82 for every power estimate and ≥ 0.94 for the theta, alpha, low-beta and high-beta bands. n = 17–20 participants per cell.

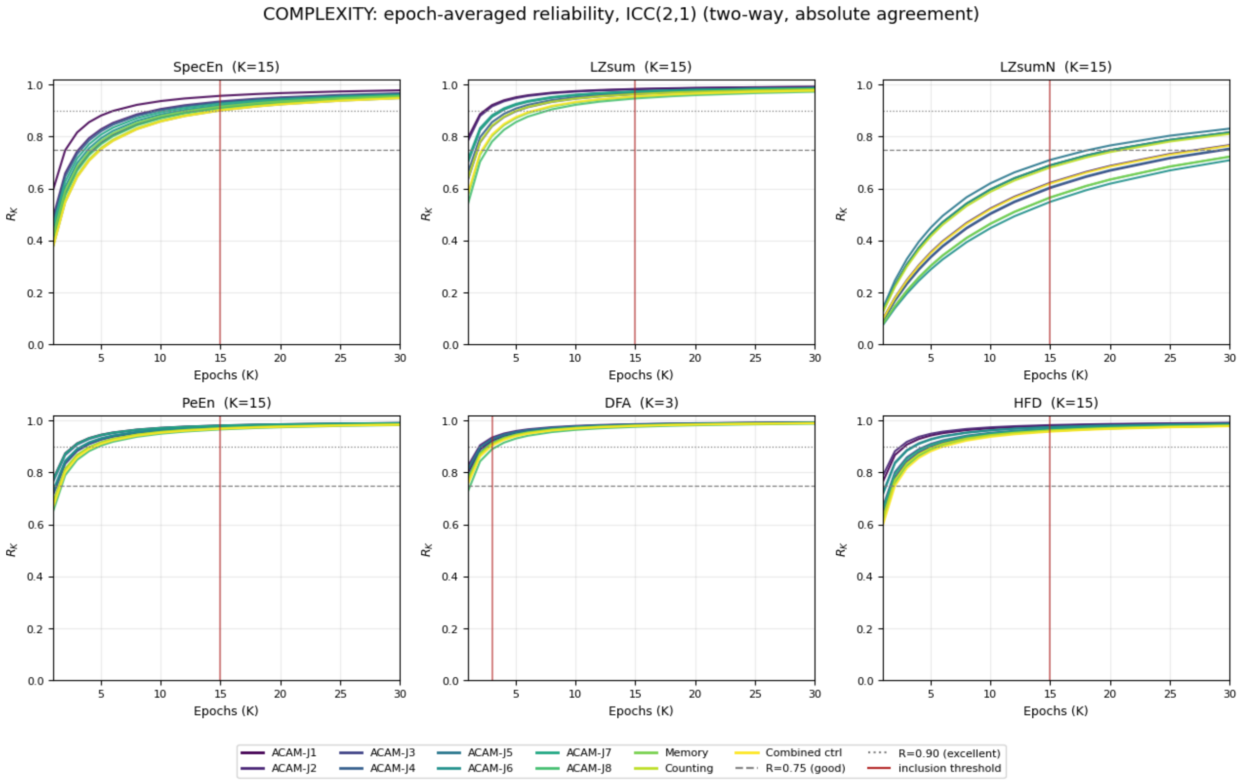

**Supplementary Figure 19.** Between-subject reliability of the complexity measures versus number of epochs averaged: spectral entropy (SpecEn), Lempel–Ziv complexity (LZsum), phase-normalised Lempel-Ziv complexity (LZsumN), permutation entropy (PeEn), detrended fluctuation analysis (DFA) and Higuchi fractal dimension (HFD). Dashed and dotted lines mark 0.75 (good) and 0.90 (excellent), the vertical line marks the inclusion threshold (threshold = 15 epochs for all measures except DFA, 3).

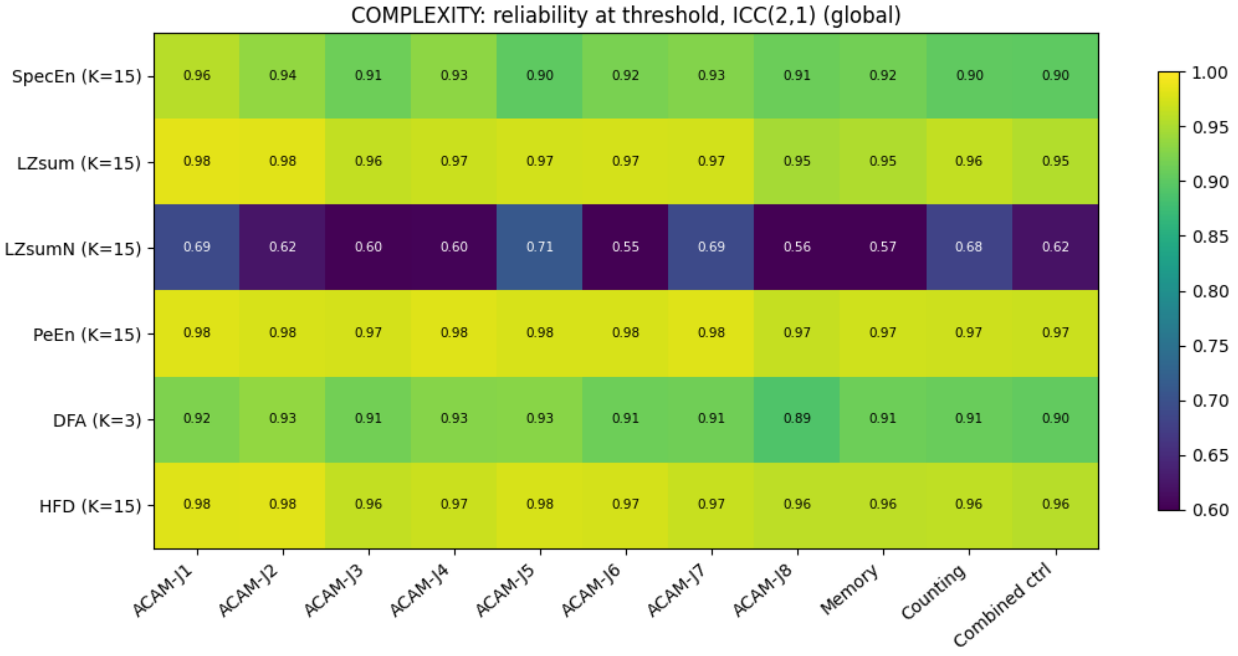

**Supplementary Figure 20.** Between-subject reliability of the complexity measures at their inclusion thresholds, for every measure × condition. All measures were good-to-excellent except LZsumN, whose phase-surrogate normalisation leaves little reliable between-participant variance (R ≈ 0.55–0.71), the others reached R ≥ 0.89. n = 15–20 participants per cell.

##### Contrast reliability

For the ACAM-J vs combined-control contrast, the complexity measures entered into the analyses were reliably estimated at 15 epochs per condition (LZsum 0.85, permutation entropy 0.84, spectral entropy 0.80, Higuchi fractal dimension 0.80, Supplementary Figure 21). DFA at three epochs was marginal (0.49; about eight epochs per condition were needed to reach 0.75), and LZsumN was unreliable (0.27). Among power components the contrast was most reliable in the high-beta band (0.74–0.76 at six epochs); lower-frequency and oscillatory contrasts were less reliably resolved at the individual level at six epochs (theta lowest, 0.17–0.28).

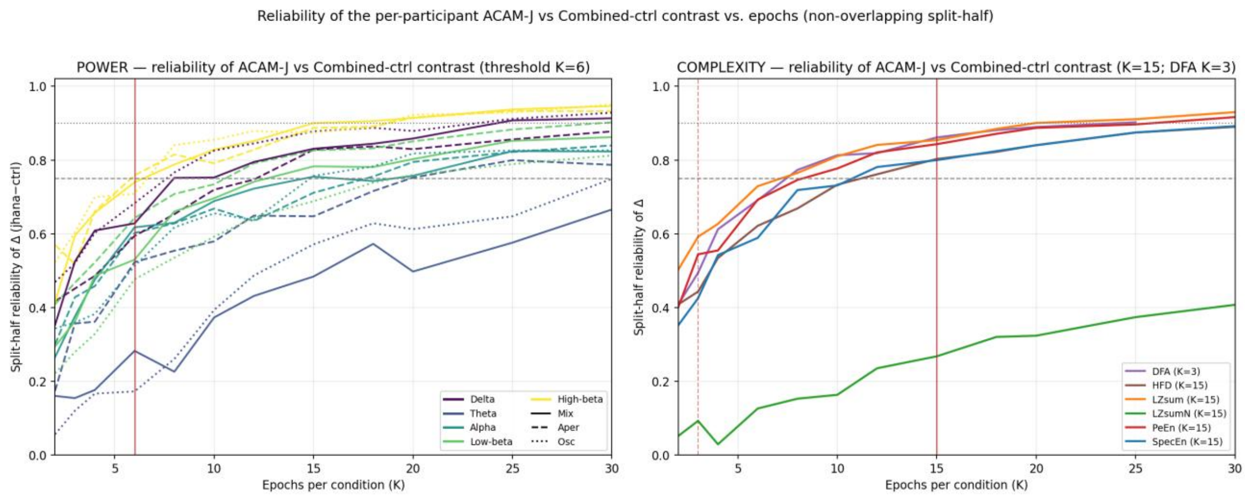

**Supplementary Figure 21.** Within-subject reliability of the ACAM-J vs combined-control contrast versus number of epochs averaged per condition. For each participant Δ = mean (ACAM-J) – mean (combined control) was estimated from two non-overlapping halves of K epochs per condition (channel-averaged), reliability is the correlation of the two half-estimates across participants (60 resamples averaged). (Left) IRASA power (colour = band, line style = total/aperiodic/oscillatory), (Right) complexity measures. Dashed and dotted lines mark 0.75 and 0.90, vertical lines mark the inclusion thresholds (6 power, 15 complexity, 3 DFA). n = up to 20 participants per point.

### Discussion

Across the measures examined, the phase-normalised Lempel-Ziv complexity (LZsumN) was an outlier. Whereas the un-normalised complexity (LZsum) and every other measure reached good-to-excellent reliability at their inclusion thresholds, LZsumN stayed below the “good” criterion at both the between- and within-participant level, even after averaging 15 epochs. This dissociation is informative rather than an artifact, and it follows directly from what the normalization does to the signal.

Lempel-Ziv complexity of broadband EEG is largely determined by the signal’s power spectrum [44,103]. Phase-normalization divides the raw complexity by the mean complexity of phase-randomized surrogates of the same epoch, surrogates that preserve the epoch's power spectrum, specifically in order to strip out that spectral contribution. By construction it therefore discards the component that dominates the raw measure, which is also the component that carries most of its reliable variance, leaving a small residual dominated by noise.

Two implications follow. Surrogate normalisation of Lempel-Ziv complexity entails a genuine trade-off. It controls for spectral confounds but at a substantial cost in reliability. Where isolating complexity from concurrent spectral change is not essential, the un-normalised entropy-rate complexity (LZsum) is considerably more reliable, where the normalised measure is retained, it requires many more epochs than its un-normalised counterpart to be considered reliable and should be interpreted accordingly. To our knowledge this reliability cost has not previously been quantified for Lempel-Ziv complexity, but it is the predictable consequence of the redundancy between complexity and the power spectrum together with the classical unreliability of residualised scores.

**Interpretation for the present study**

With these exceptions noted, the measures entered into the main analyses are reliably estimated at the epoch thresholds used, so the reported ACAM-J vs control effects and the between-subject correlations with phenomenology for spectral power and for SpecEn, LZsum, PeEn and HFD rest on stable per-participant estimates. Two cautions apply. First, LZsumN is unreliable, so a small or null effect for it may reflect low reliability rather than a true absence of a complexity change, and its correlations with phenomenology will be attenuated. Conversely, significant effects should be interpreted with similar caution. Second, DFA at three epochs and the lower-frequency and oscillatory power contrasts are less reliably resolved, so effects and correlations involving these warrant more caution than those for the high-reliability measures. However, 3, 5, and 15 epochs were the minimum numbers of epochs for a given run to be included in the DFA, power, and complexity analyses, respectively. The vast majority of subjects far exceeded the minimum inclusion criteria (DFA: $19.5 \pm 8.8$, power: $39.0 \pm 17.7$, complexity: $99.8 \pm 44.7$).

### Methodological considerations

Despite considerable strengths, including the first group sample spanning all eight ACAM-J states using high-density EEG with a rigorous, standardized preprocessing pipeline, a broad multi-measure battery spanning spectral and complexity domains, and a within-subject design that resolves each of the eight absorption states individually, we note some limitations of the current study. One consideration in any EEG study is the possibility of electromyographic (EMG) contamination, which can mimic genuine cortical changes, particularly in the high-beta band, and can therefore bias broadband complexity measures. Several precautions were taken to mitigate this risk. ICA was used to remove muscle-related components. Strict epoch rejection was applied. A Laplacian montage was used to reduce volume-conduction-driven smearing of distant EMG sources. Spectral analyses were restricted to frequencies at or below 35 Hz, considerably lower than most literature exploring broadband complexity [1,26,27,38,51,55-57,67,83,104,105]. In addition, participants meditated supine, which removes the postural muscle activity associated with supporting the head in an upright position and so possibly reduces a major contributor to EMG in upright EEG paradigms [106]. As an additional control analysis, correlations were computed between the average total-power change in posterior clusters related to increased high-beta power, as well as in centrally located clusters showing decreased power, and self-reported automatic movement during ACAM-J1 and ACAM-J2 (where these phenomenological reports were available). These correlations were not significant, suggesting that automatic movement did not influence the current results (see supplementary figure 12). Despite these precautions, we cannot entirely exclude the possibility of residual EMG as a contributor to the posterior high-beta increases and, consequently, to the broadband entropy estimates. The replication of increased high-frequency band power and aperiodic-flattening pattern in the independent jhana dataset of Mago et al. (2026) somewhat alleviates this concern, as it demonstrates consistency of these effects across cohorts.

Further, the progressive strengthening of all three families of power changes across successive ACAM-J states might in principle be explained by slow drift. This is unlikely, as a 1 Hz high-pass filter was applied during preprocessing, which removes drift in the frequency range that would otherwise affect broadband power. More importantly, the phenomenology of ACAM-J is itself described as a monotonically deepening unification of mind and object, with each successive state entailing a more pronounced departure from ordinary consciousness. The across-state gradient in the EEG measures reported here thus tracks an underlying phenomenological gradient.

### Qualitative epoch stability analysis

**Supplementary Figures 22.** Stability of group-average IRASA total, aperiodic, oscillatory, (delta, theta, alpha, low beta, high beta) power estimates as a function of the number of sampled epochs. Separate panels show the memory condition, counting condition, and ACAM-J1 to ACAM-J8. For each state, global linear power was recomputed after randomly sampling k clean 5-s epochs (k = 1–10) from the pooled subject-level epoch set, including only subject-state combinations with at least 10 available epochs. The solid line shows the mean across eligible subjects, and the dashed horizontal line indicates the corresponding full-data group estimate obtained when all available clean epochs were included. This analysis was performed separately for the total, aperiodic, and oscillatory IRASA components and for each canonical frequency band.

Stability of group-average global LZsum, LZsumN, PeEn, SpecEn, HFD across states as a function of the number of sampled epochs. Separate panels show the memory condition, counting condition, and ACAM-J1 to ACAM-J8. For each state, the complexity measure was recomputed after randomly sampling $k$ clean 2-s epochs from each eligible run ($k$ = 1–20; minimum 20 epochs per run). Where multiple runs were available for a subject within a state, run-level estimates were averaged to obtain a single subject-level value. The solid line shows the mean across subjects, and the dashed horizontal line marks the full-data group estimate derived from all available clean epochs in the eligible runs.

Stability of group-average global detrended fluctuation analysis (DFA) estimates across states as a function of the number of sampled epochs. Panels show the memory condition, counting condition, and ACAM-J1 to ACAM-J8. For each state, global DFA was recomputed after randomly sampling $k$ clean 10-s epochs ($k$ = 1–10) from the pooled subject-level epoch set, including only subject-state combinations with at least 10 available epochs. The solid line shows the mean across eligible subjects, and the dashed horizontal line marks the full-data group estimate obtained when all available clean epochs were included. The figure therefore illustrates how the group-average DFA estimate approaches the full-data reference as more epochs are incorporated.

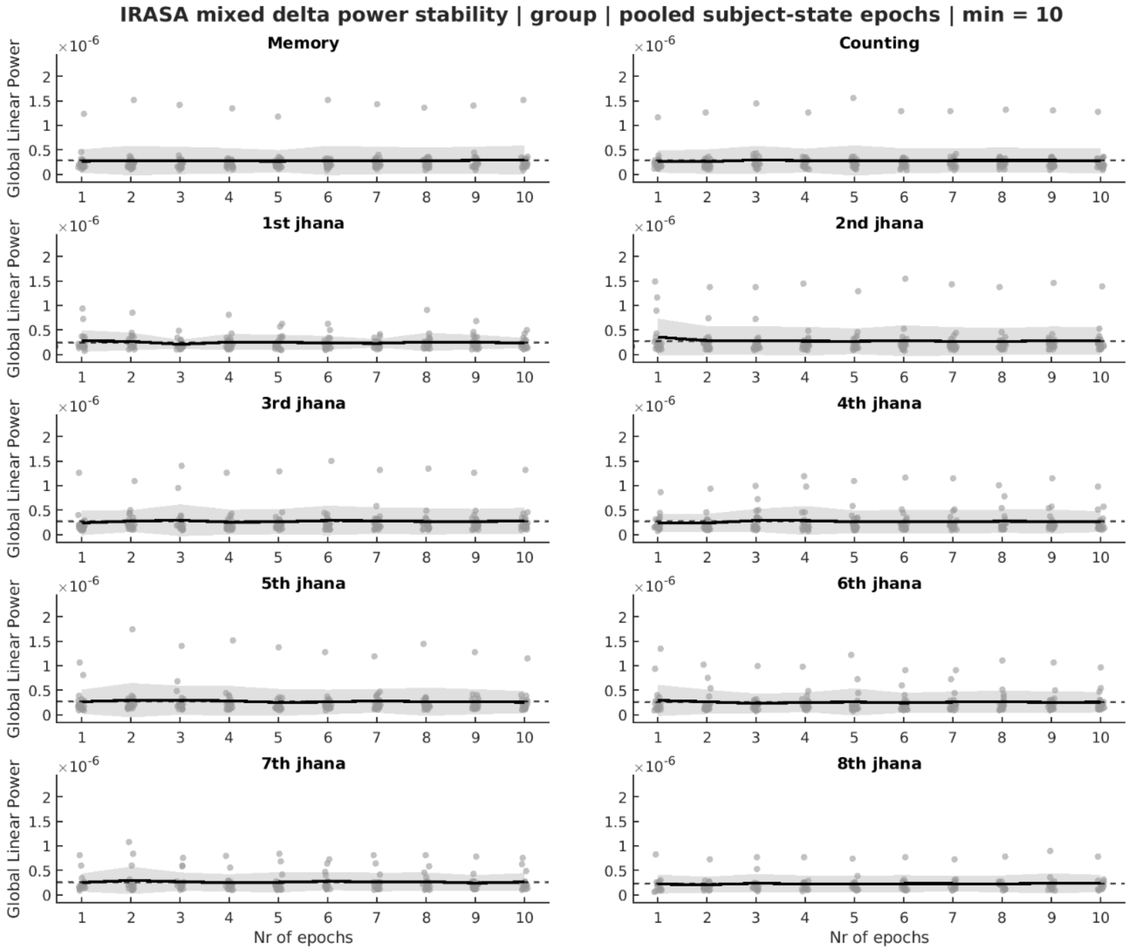
IRASA mixed delta power stability | group | pooled subject-state epochs | min = 10
Memory
Counting
1st jhana
2nd jhana
3rd jhana
4th jhana
5th jhana
6th jhana
7th jhana
8th jhana
Global Linear Power
Nr of epochs
×10^-6

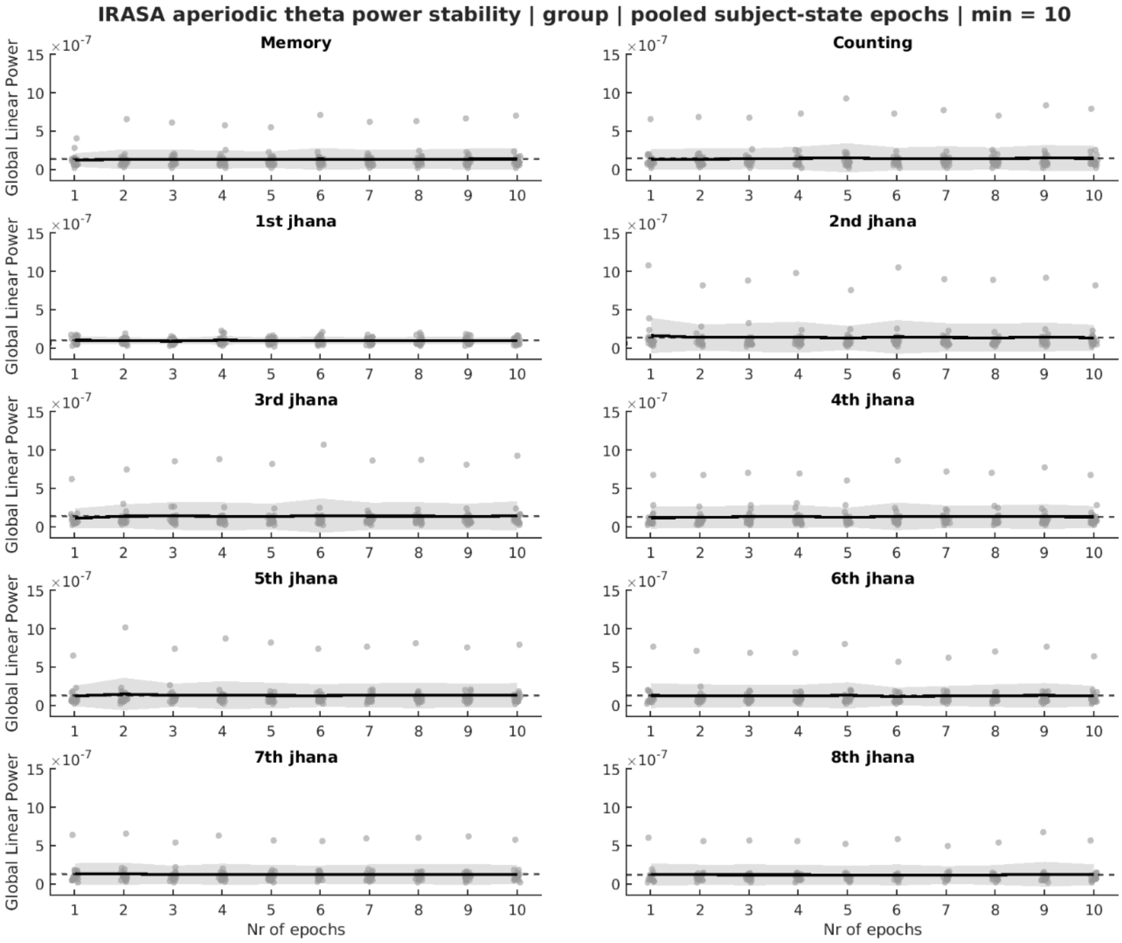
IRASA aperiodic theta power stability | group | pooled subject-state epochs | min = 10
Memory
Counting
1st jhana
2nd jhana
3rd jhana
4th jhana
5th jhana
6th jhana
7th jhana
8th jhana
Global Linear Power
×10^-7
Nr of epochs

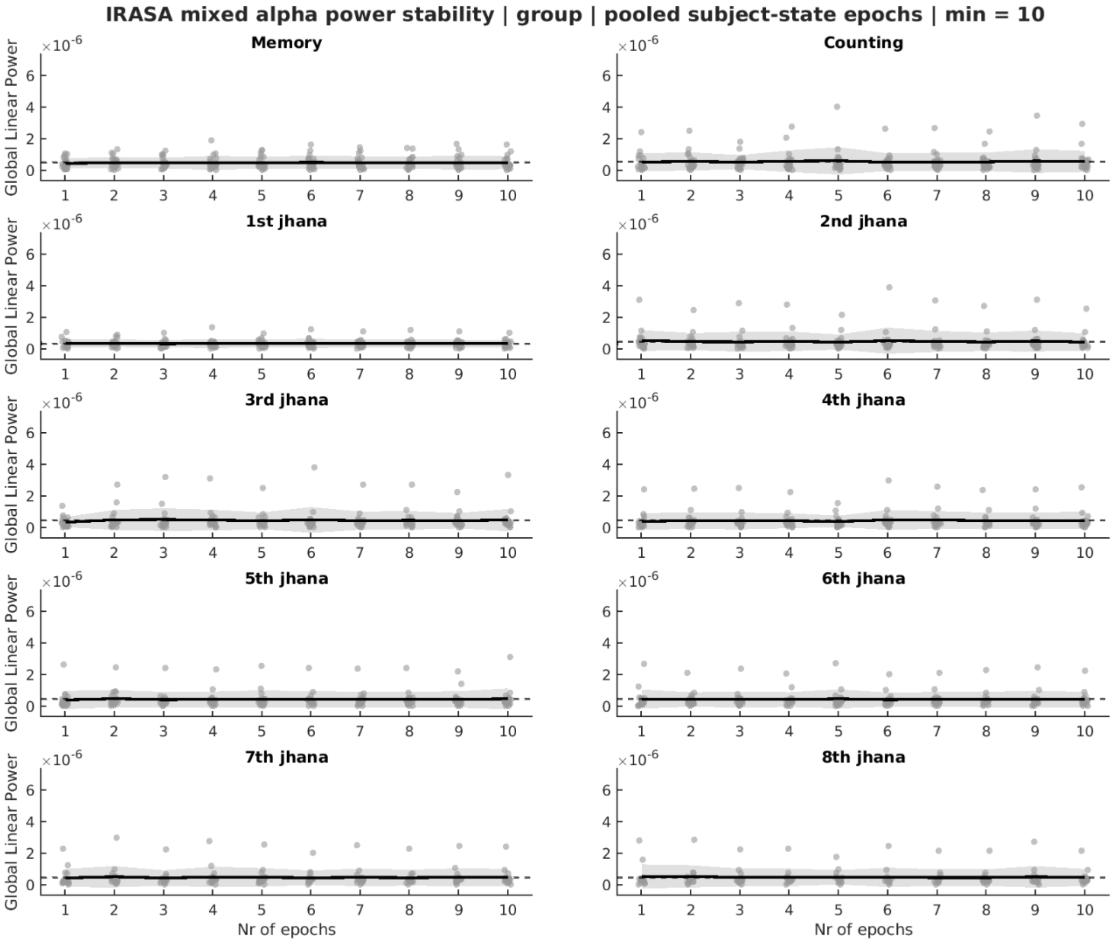
IRASA mixed alpha power stability | group | pooled subject-state epochs | min = 10
Memory
Counting
1st jhana
2nd jhana
3rd jhana
4th jhana
5th jhana
6th jhana
7th jhana
8th jhana
Global Linear Power
×10^-6
Nr of epochs

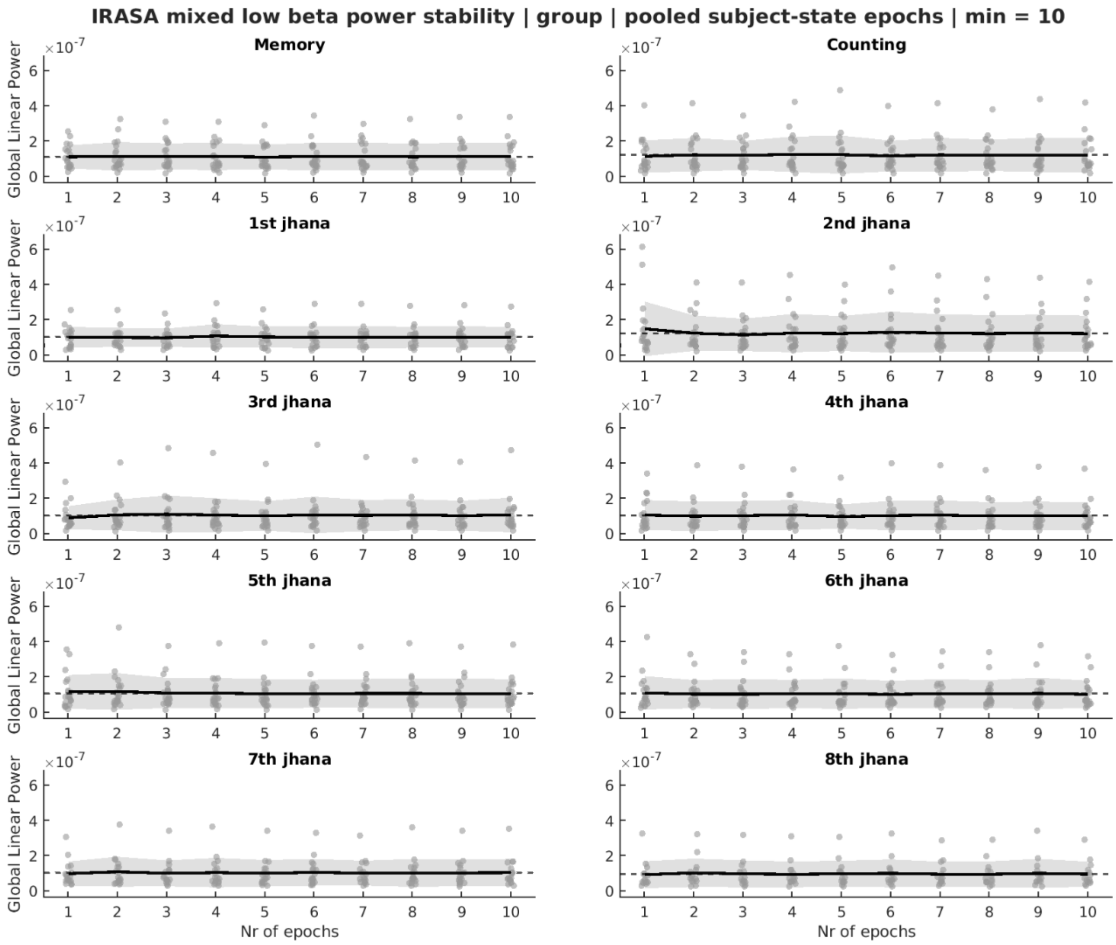
IRASA mixed low beta power stability | group | pooled subject-state epochs | min = 10
Memory
Counting
1st jhana
2nd jhana
3rd jhana
4th jhana
5th jhana
6th jhana
7th jhana
8th jhana
Global Linear Power
×10^-7
Nr of epochs

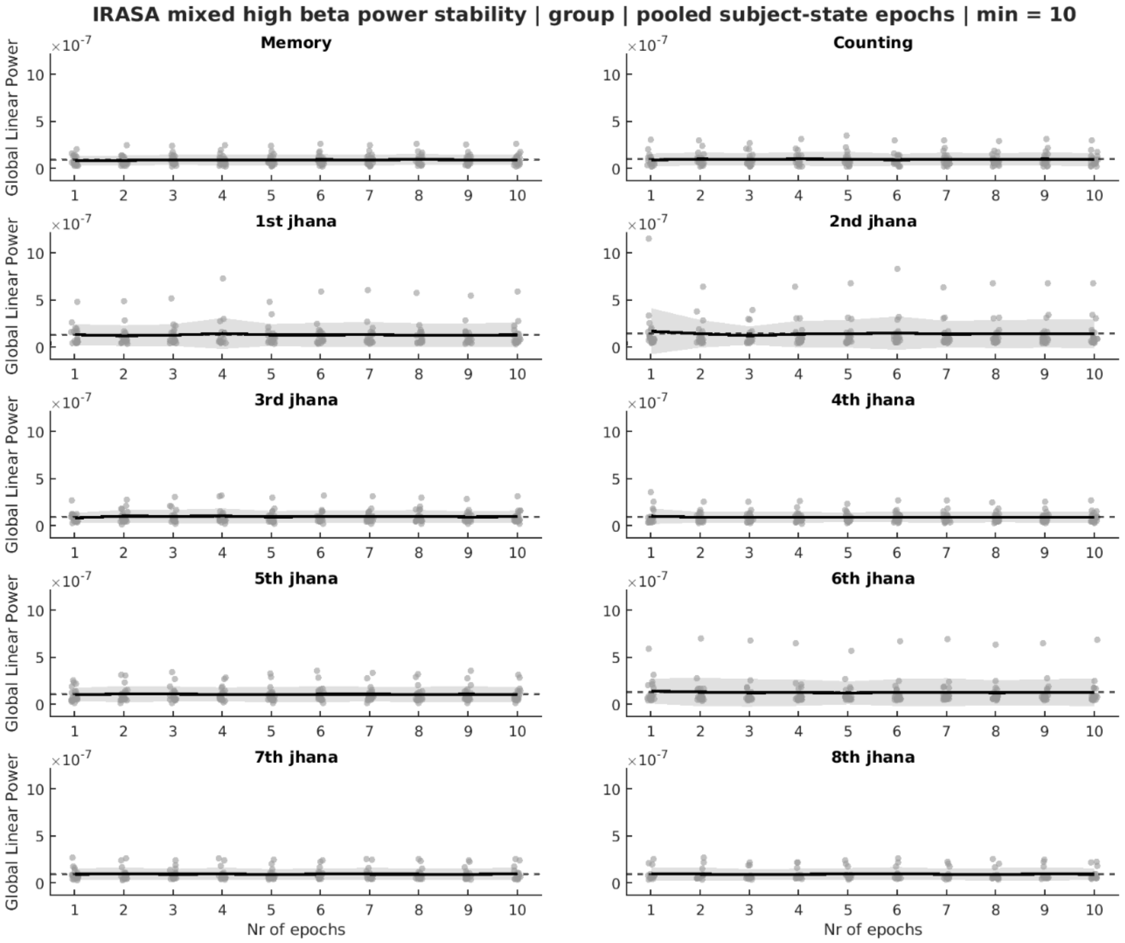

IRASA mixed high beta power stability | group | pooled subject-state epochs | min = 10
Memory
Counting
1st jhana
2nd jhana
3rd jhana
4th jhana
5th jhana
6th jhana
7th jhana
8th jhana
Global Linear Power
×10^-7
Nr of epochs

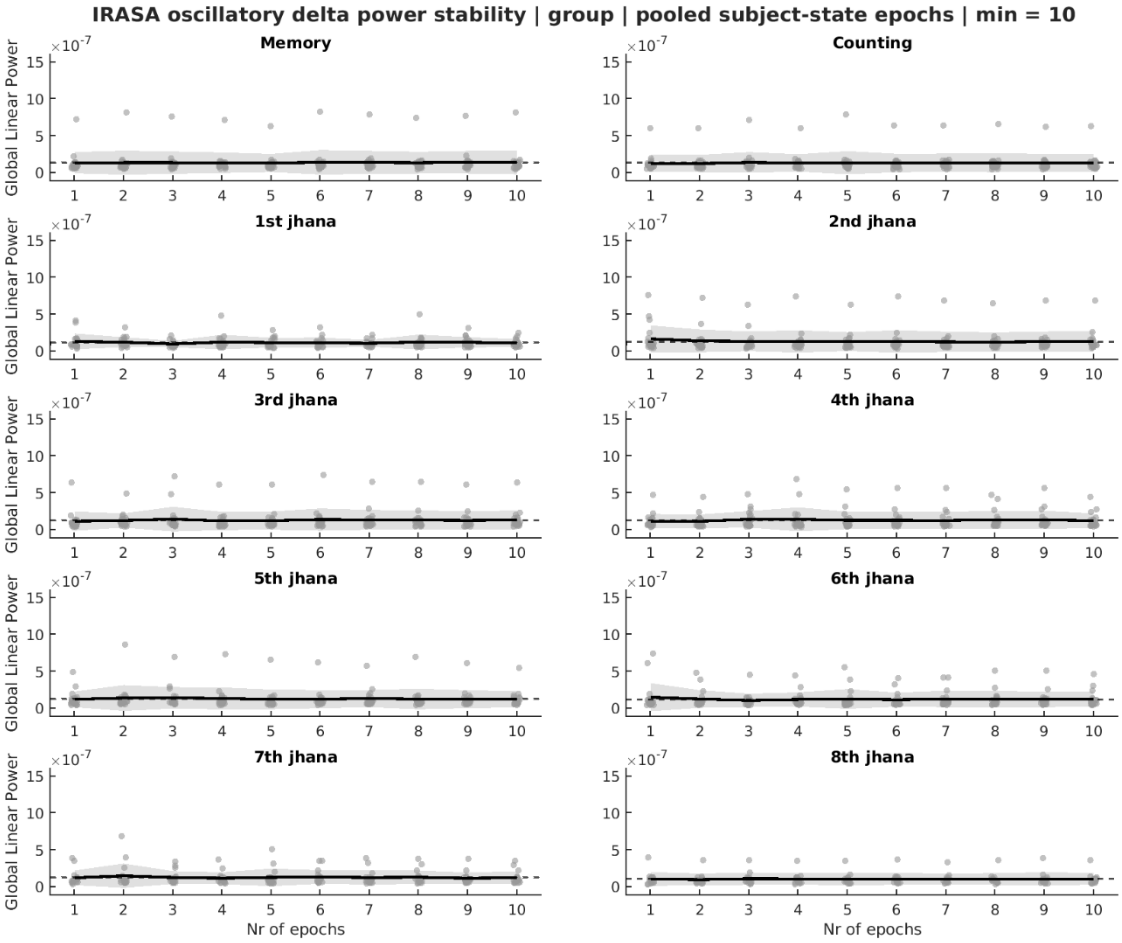
IRASA oscillatory delta power stability | group | pooled subject-state epochs | min = 10
Memory
Counting
1st jhana
2nd jhana
3rd jhana
4th jhana
5th jhana
6th jhana
7th jhana
8th jhana
Global Linear Power
×10^-7
Nr of epochs

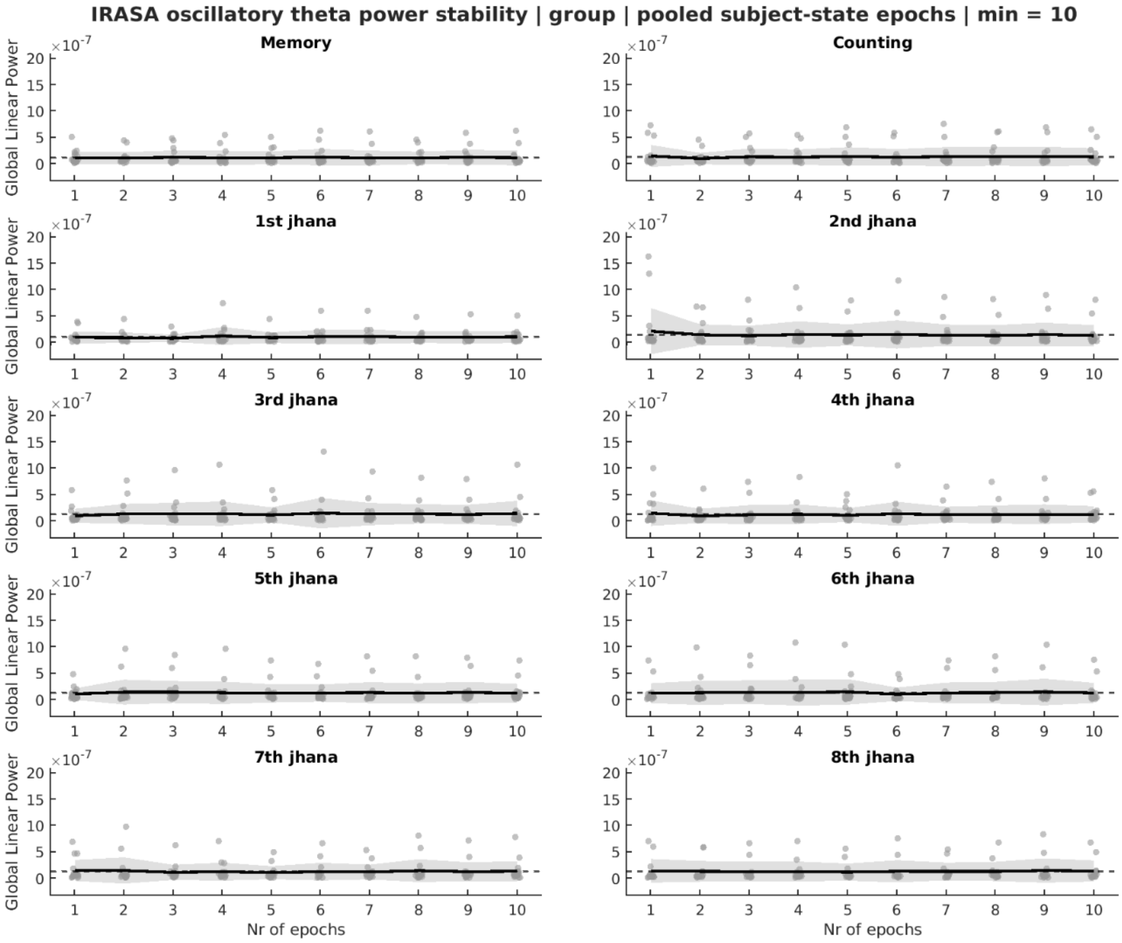
IRASA oscillatory theta power stability | group | pooled subject-state epochs | min = 10
Memory
Counting
1st jhana
2nd jhana
3rd jhana
4th jhana
5th jhana
6th jhana
7th jhana
8th jhana
Global Linear Power
×10^-7
Nr of epochs

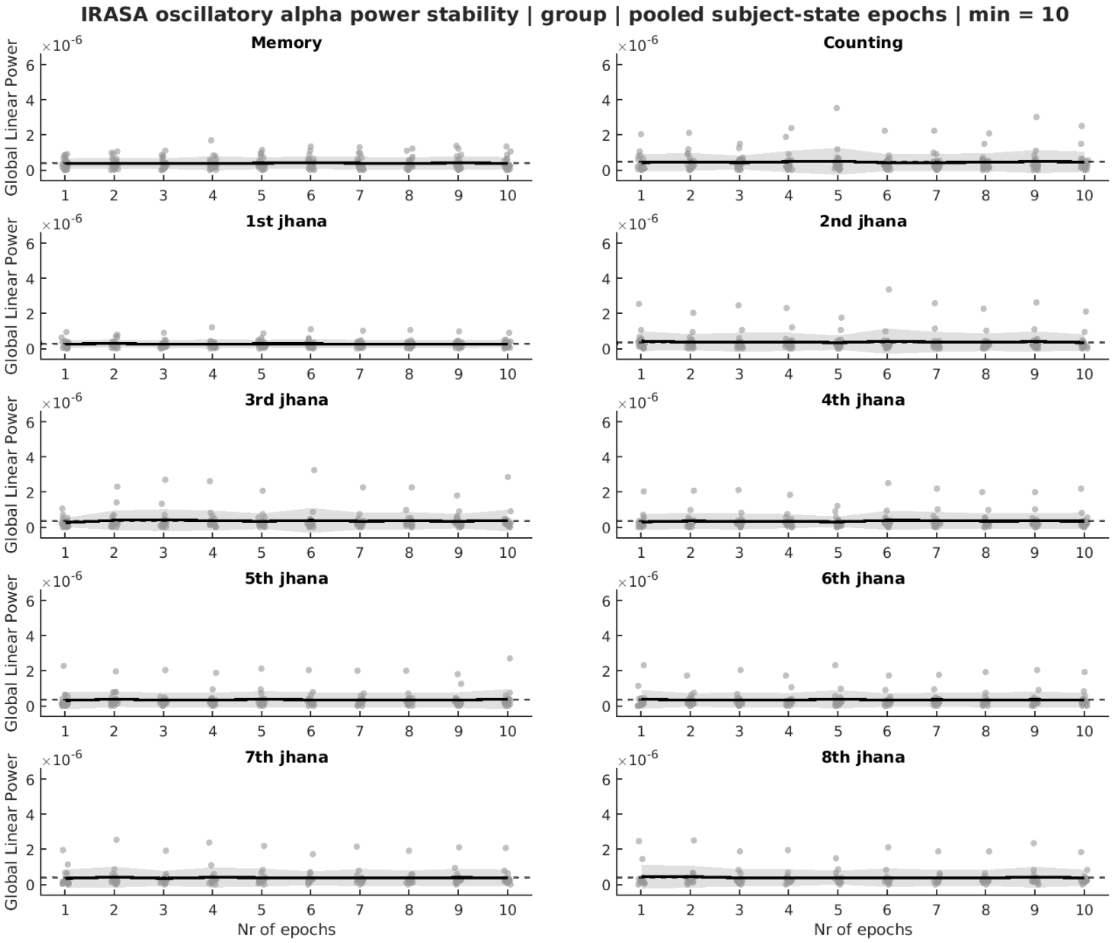
IRASA oscillatory alpha power stability | group | pooled subject-state epochs | min = 10
Memory
Counting
1st jhana
2nd jhana
3rd jhana
4th jhana
5th jhana
6th jhana
7th jhana
8th jhana
Global Linear Power
×10^-6
Nr of epochs

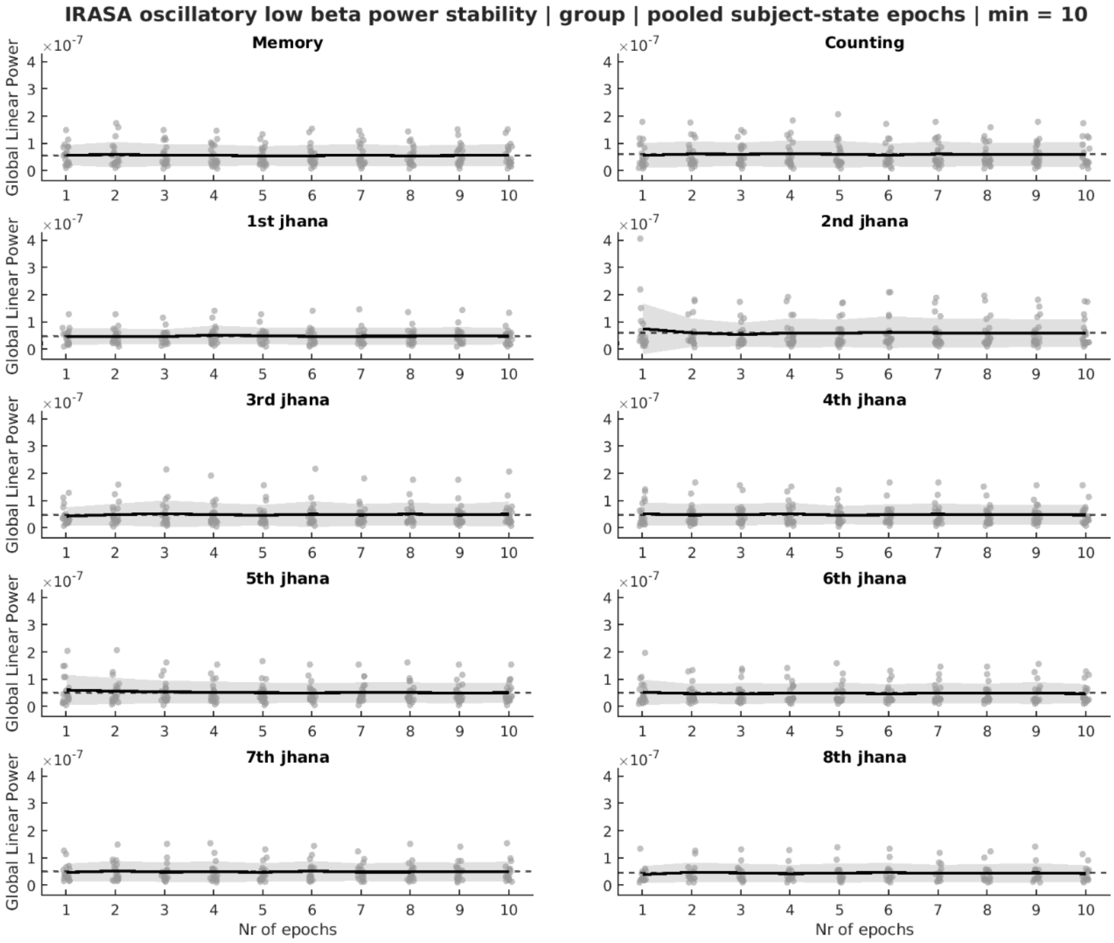

IRASA oscillatory low beta power stability | group | pooled subject-state epochs | min = 10
Memory
Counting
1st jhana
2nd jhana
3rd jhana
4th jhana
5th jhana
6th jhana
7th jhana
8th jhana
Global Linear Power
×10^-7
Nr of epochs

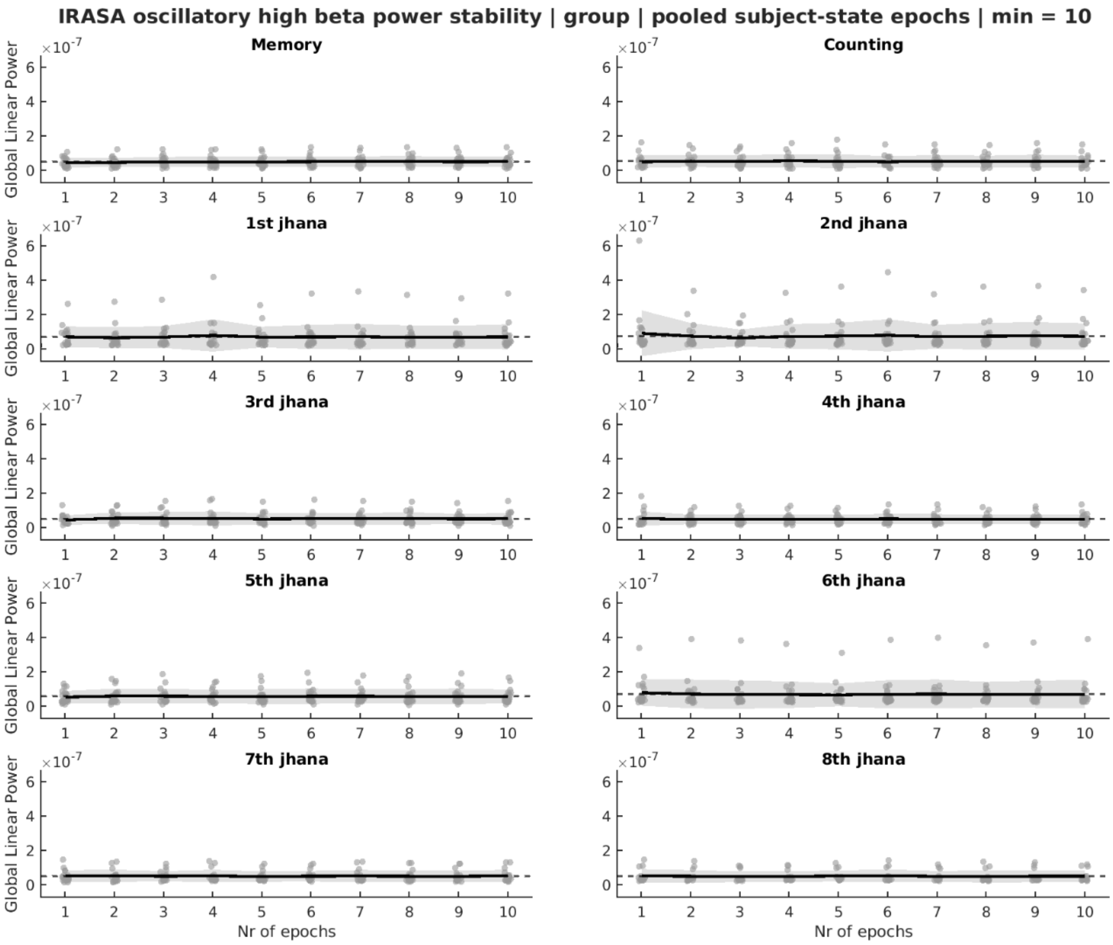
IRASA oscillatory high beta power stability | group | pooled subject-state epochs | min = 10
Memory
Counting
1st jhana
2nd jhana
3rd jhana
4th jhana
5th jhana
6th jhana
7th jhana
8th jhana
Global Linear Power
$\times 10^{-7}$
Nr of epochs

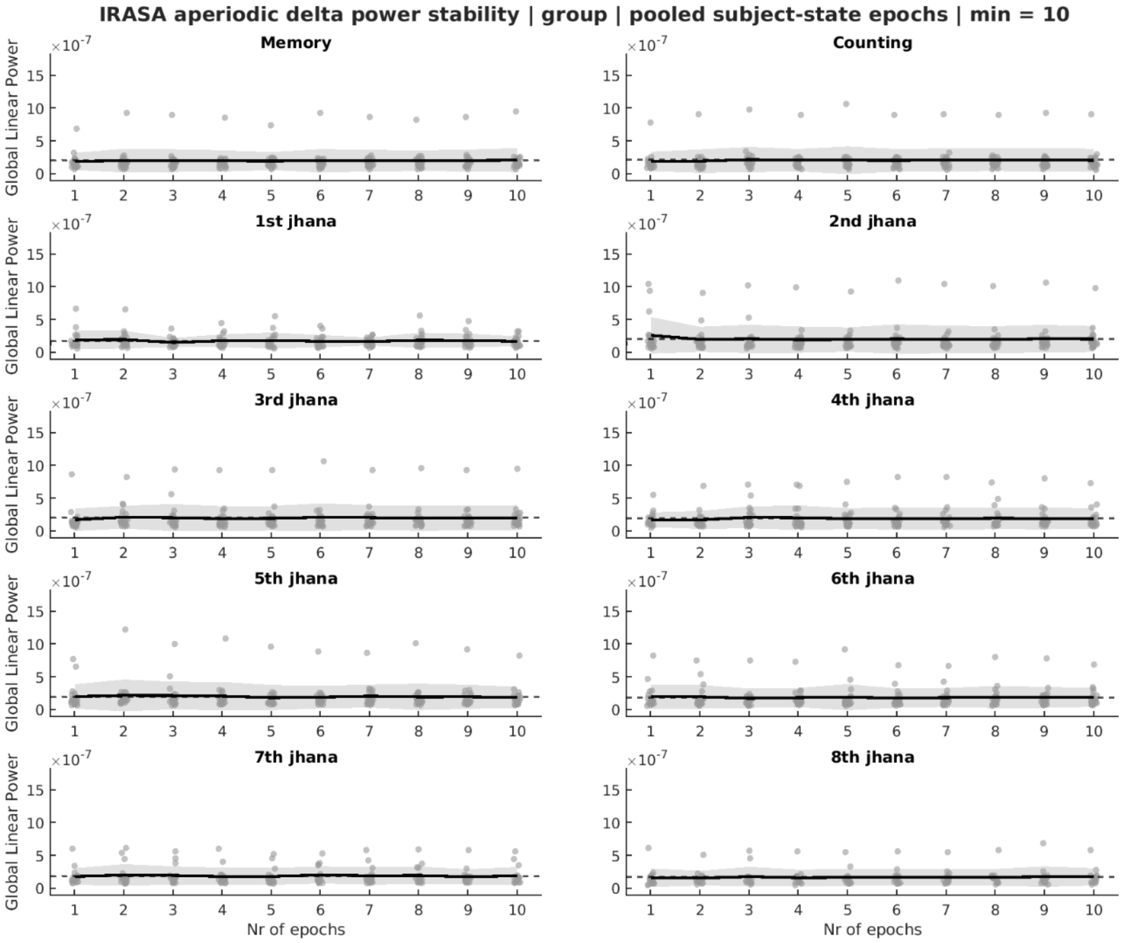

IRASA aperiodic delta power stability | group | pooled subject-state epochs | min = 10
Memory
Counting
1st jhana
2nd jhana
3rd jhana
4th jhana
5th jhana
6th jhana
7th jhana
8th jhana
Global Linear Power
Nr of epochs

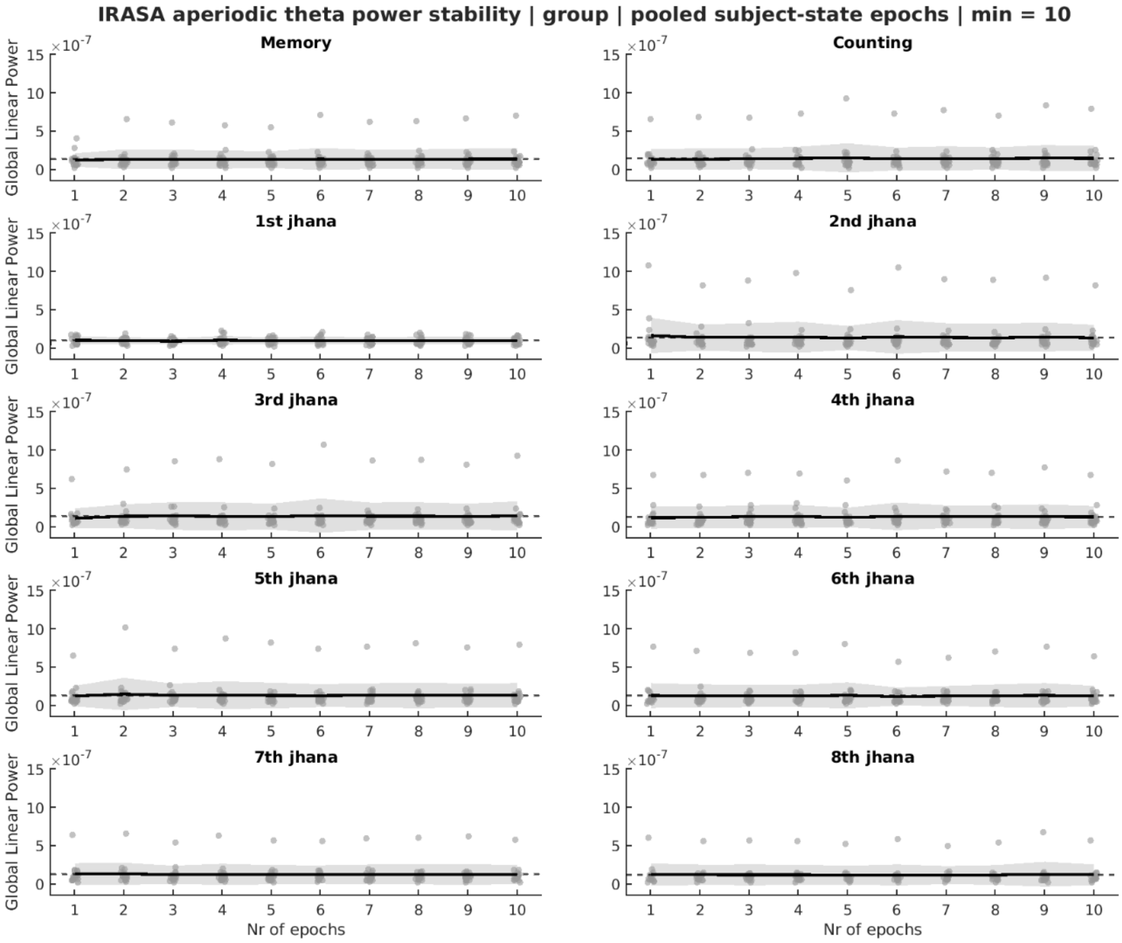

IRASA aperiodic theta power stability | group | pooled subject-state epochs | min = 10
Memory
Counting
1st jhana
2nd jhana
3rd jhana
4th jhana
5th jhana
6th jhana
7th jhana
8th jhana
Global Linear Power
×10^-7
Nr of epochs

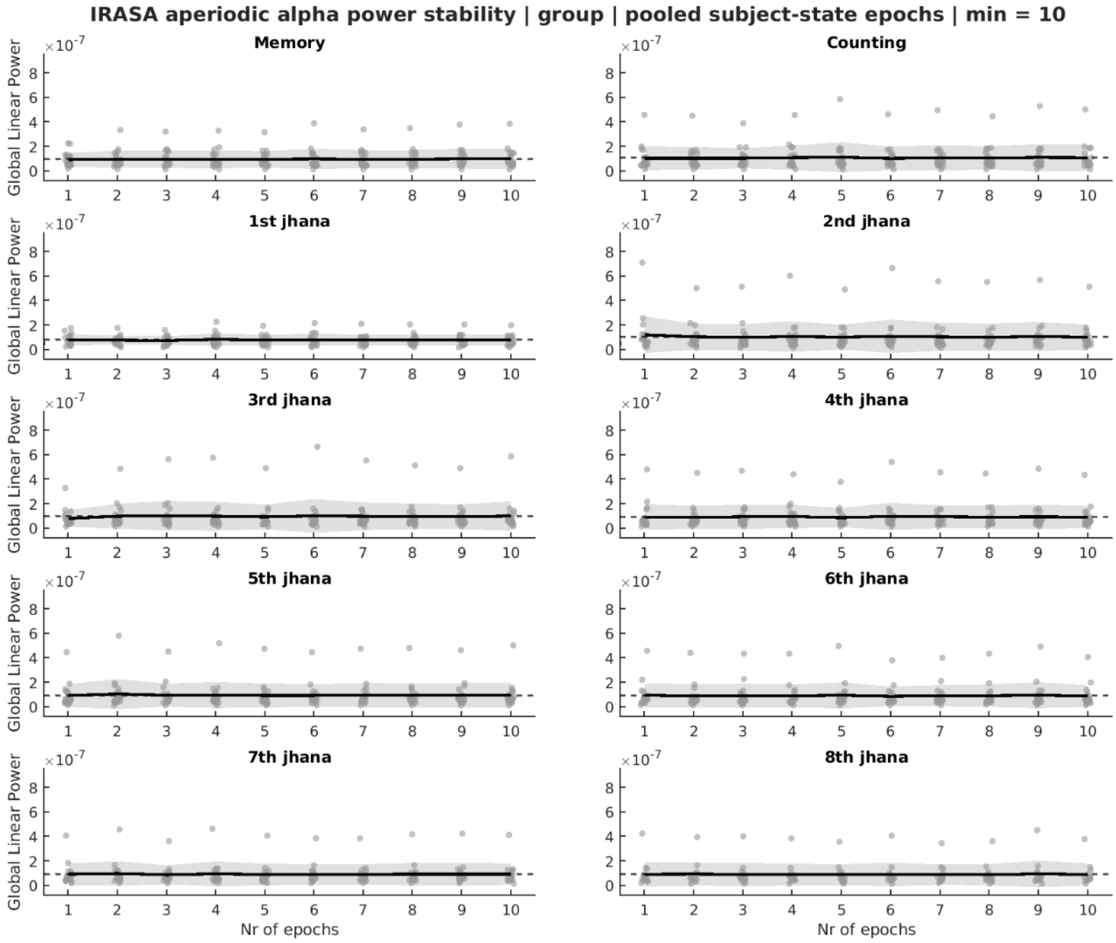
IRASA aperiodic alpha power stability | group | pooled subject-state epochs | min = 10
Memory
Counting
1st jhana
2nd jhana
3rd jhana
4th jhana
5th jhana
6th jhana
7th jhana
8th jhana
Global Linear Power
×10^-7
Nr of epochs

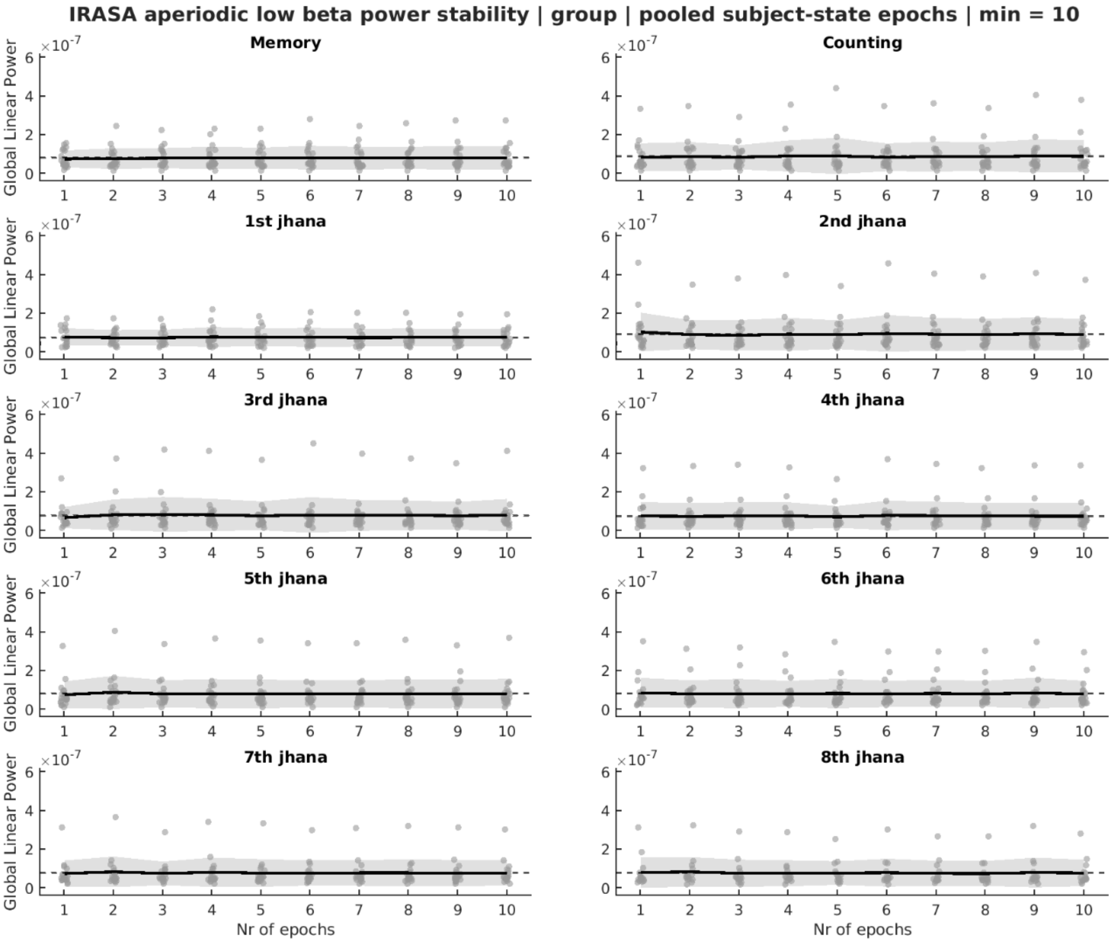
IRASA aperiodic low beta power stability | group | pooled subject-state epochs | min = 10
Memory
Counting
1st jhana
2nd jhana
3rd jhana
4th jhana
5th jhana
6th jhana
7th jhana
8th jhana
Global Linear Power
×10^-7
Nr of epochs

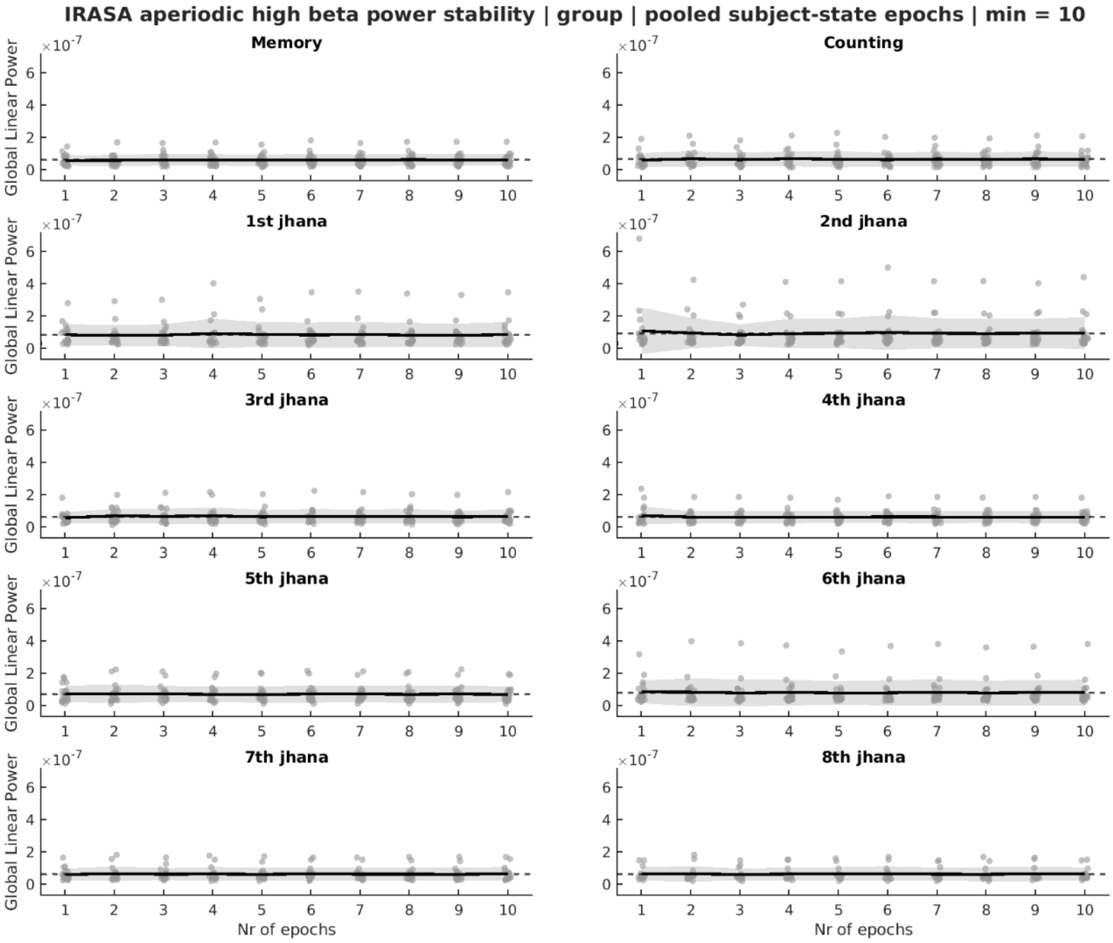

IRASA aperiodic high beta power stability | group | pooled subject-state epochs | min = 10
Memory
Counting
1st jhana
2nd jhana
3rd jhana
4th jhana
5th jhana
6th jhana
7th jhana
8th jhana
Global Linear Power
×10^-7
Nr of epochs

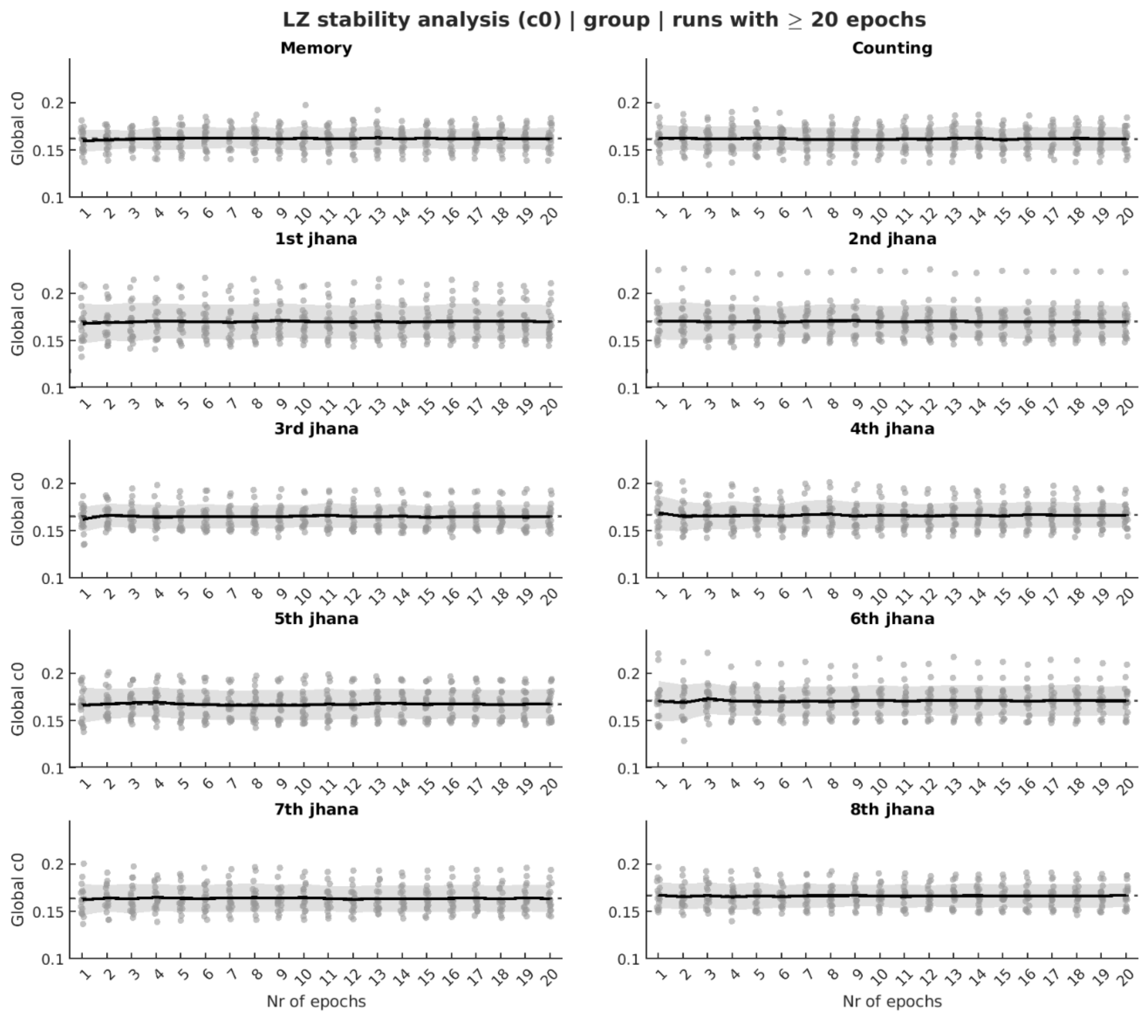
LZ stability analysis (c0) | group | runs with ≥ 20 epochs
Memory
Counting
1st jhana
2nd jhana
3rd jhana
4th jhana
5th jhana
6th jhana
7th jhana
8th jhana
Global c0
Nr of epochs

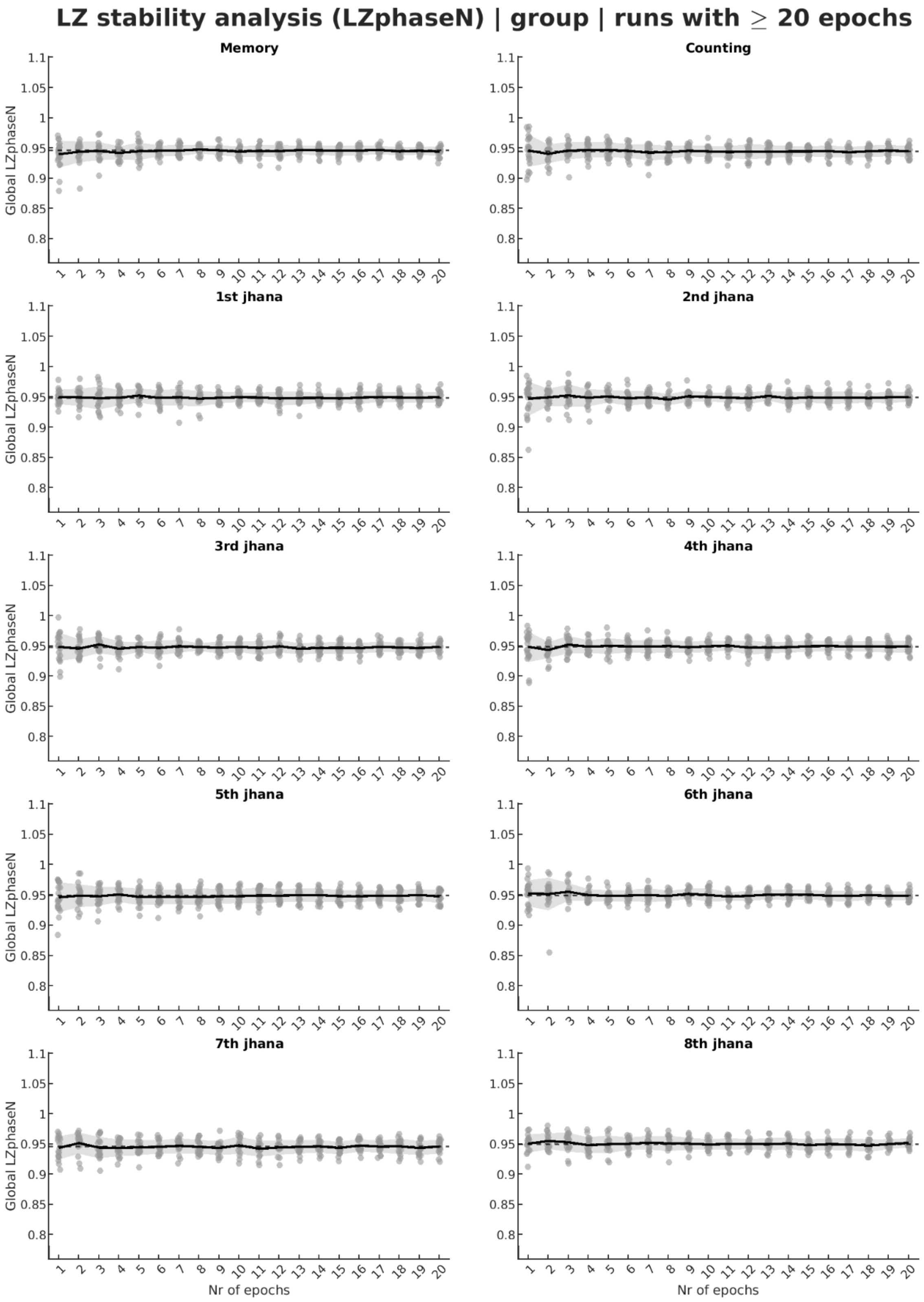
LZ stability analysis (LZphaseN) | group | runs with ≥ 20 epochs
Memory
Counting
1st jhana
2nd jhana
3rd jhana
4th jhana
5th jhana
6th jhana
7th jhana
8th jhana
Global LZphaseN
Nr of epochs

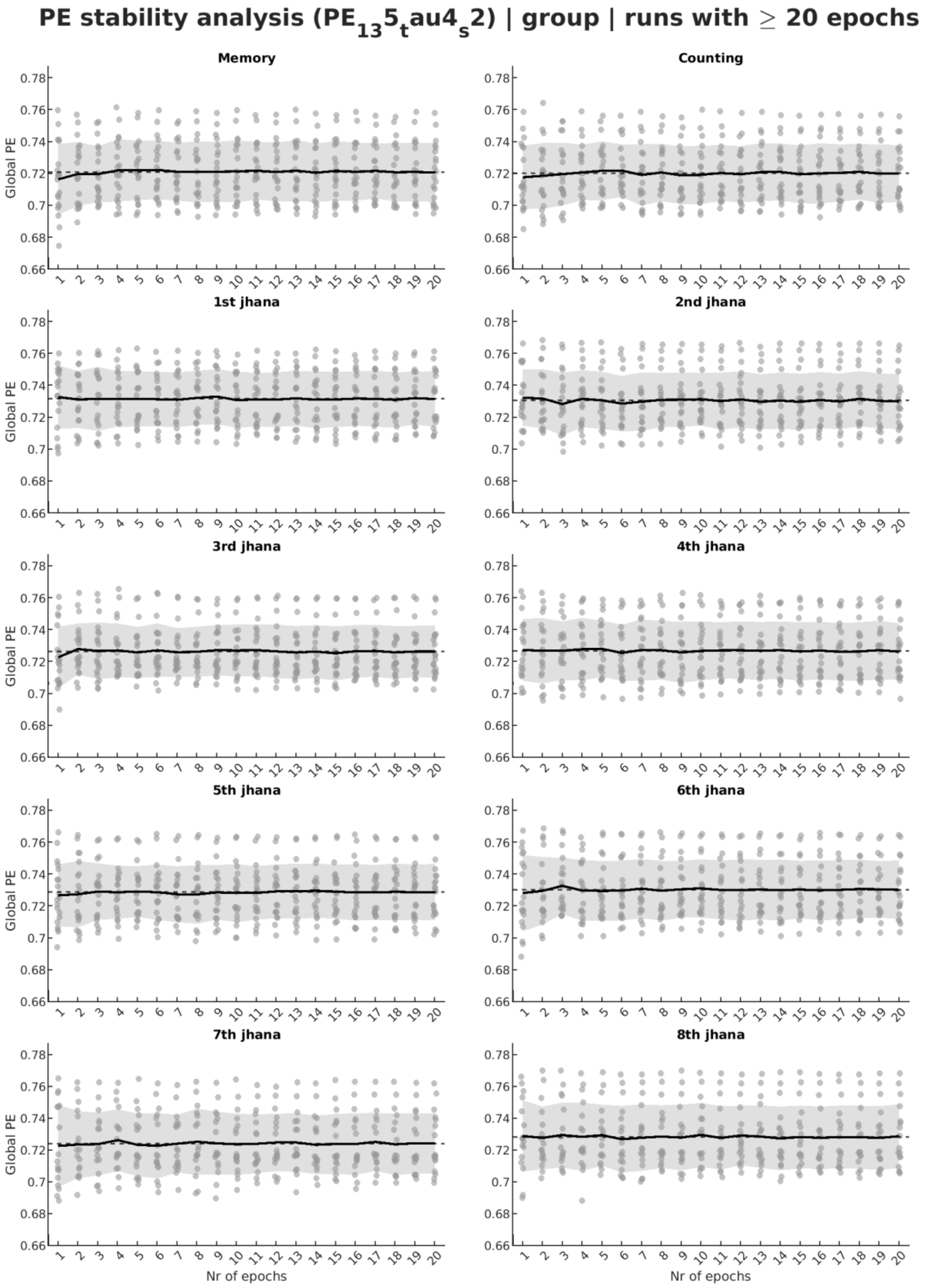
PE stability analysis (PE_135_tau4_s2) | group | runs with ≥ 20 epochs
Memory
Counting
1st jhana
2nd jhana
3rd jhana
4th jhana
5th jhana
6th jhana
7th jhana
8th jhana
Global PE
Nr of epochs

tral entropy stability analysis (SpecEn$_1$to35Hz) | group | runs with ≥ 20 e

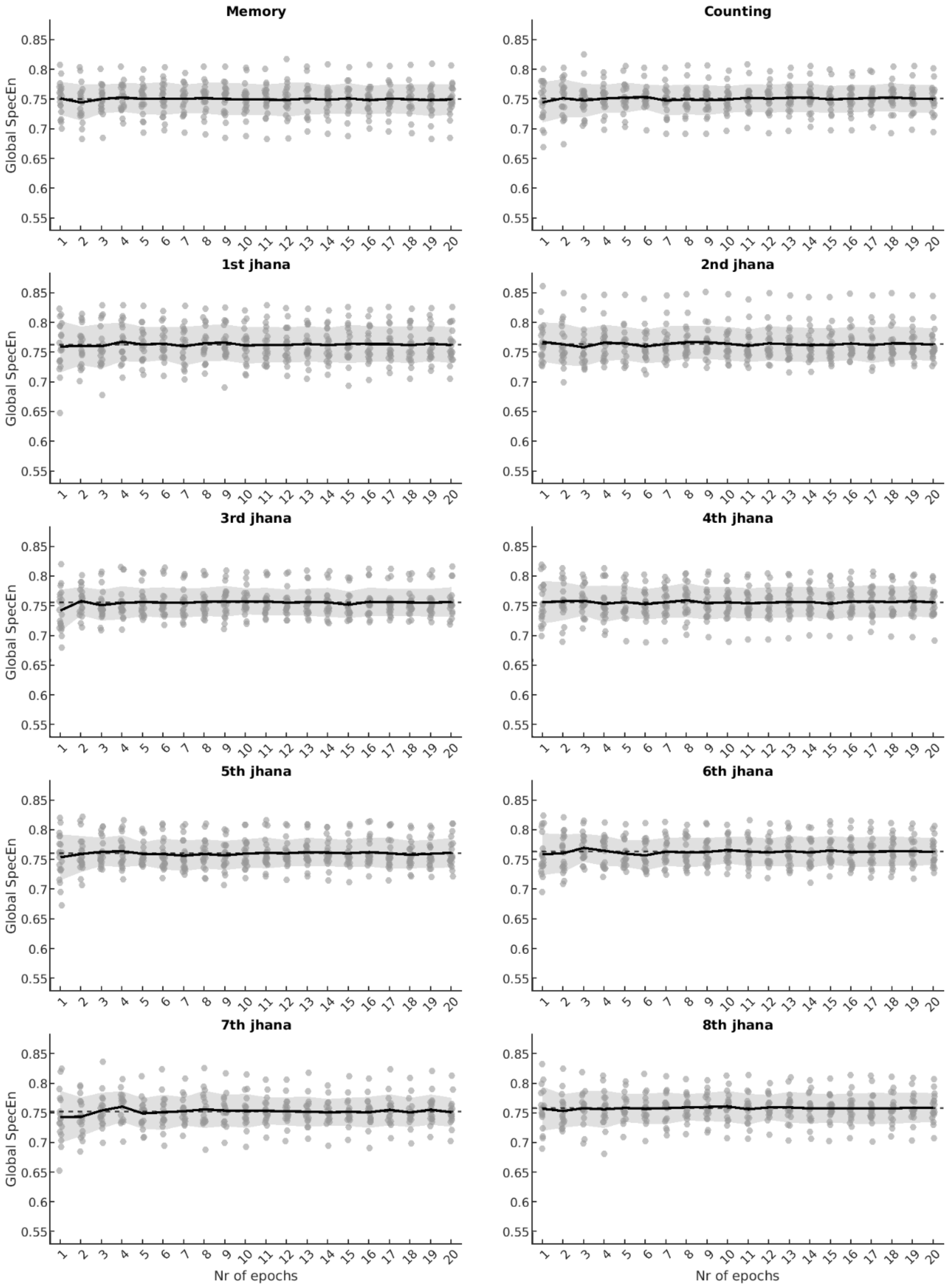

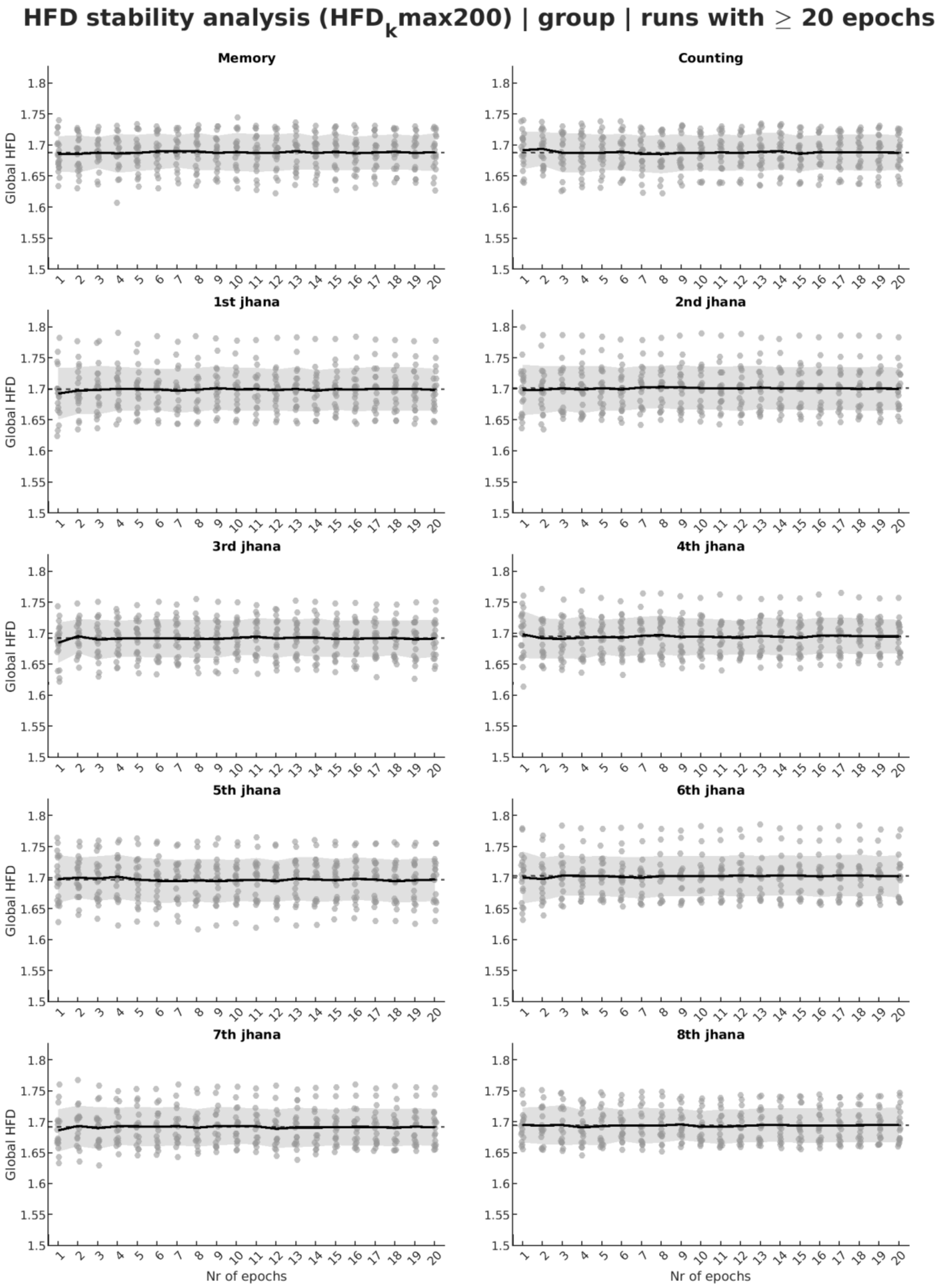
HFD stability analysis (HFDkmax200) | group | runs with ≥ 20 epochs
Memory
Counting
1st jhana
2nd jhana
3rd jhana
4th jhana
5th jhana
6th jhana
7th jhana
8th jhana
Global HFD
Nr of epochs

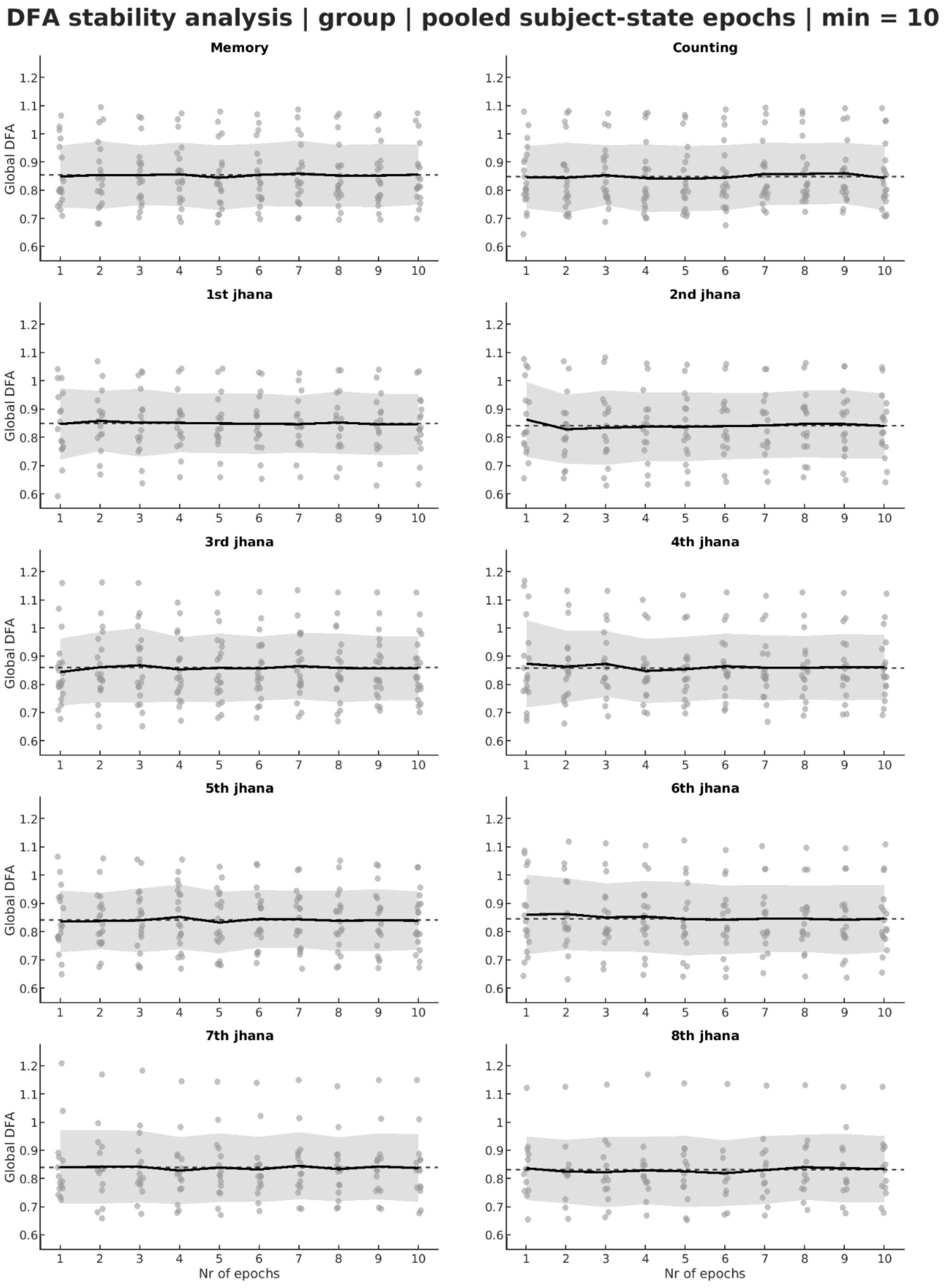
DFA stability analysis | group | pooled subject-state epochs | min = 10
Memory
Counting
1st jhana
2nd jhana
3rd jhana
4th jhana
5th jhana
6th jhana
7th jhana
8th jhana
Global DFA
Nr of epochs
